# Haemagglutinin 162–164 deletions enhance influenza B/Victoria virus fitness and virulence in vivo

**DOI:** 10.64898/2026.01.08.698527

**Authors:** Giselle G.K. Ng, Peter Cronin, Xiang Li, Carla Bianca L. Victorio, Dong-Qiang Cheng, Rong Zhang, Foong Ying Wong, Arun Ganasarajah, Ann-Marie Chacko, Richard J. Webby, Paul A. Tambyah, Gavin J.D. Smith, Yvonne C.F. Su

## Abstract

Influenza B viruses cause substantial respiratory disease and seasonal outbreaks. Despite decades of circulation in humans, only the B/Victoria lineage persisted after the COVID-19 pandemic. Continual evolution has generated hemagglutinin deletion variants at residues 162–164 that drive successive epidemics, yet their functional consequences remain poorly understood. Using integrated phylodynamics and reverse genetics, we show that Clade V1A.1 viruses carrying a two-amino acid deletion exhibit enhanced replication and increased virulence compared with ancestral viruses lacking deletions. The recently prevailing Clade V1A.3, which harbors a three-amino acid deletion together with the K136E substitution, has completely displaced V1A.1 and causes more severe disease in mice. Both clades bound efficiently to alpha 2-3 and 2-6 sialylated glycans and exhibited broad tolerance to acidic pH and elevated temperatures. These findings reveal that specific combinations of HA deletions and substitutions confer pronounced fitness advantages to emerging variants, driving global selective sweeps, evolutionary success and long-term persistence of B/Victoria lineage, and posing challenges for vaccine efficacy and influenza control.

## INTRODUCTION

Long-term evolution and persistence of influenza B virus (IBV) continue to cause seasonal epidemics and pose a substantial disease burden, compounded by recurring challenges of vaccine mismatch. Globally, IBV causes approximately 7.9 million lower respiratory tract infections and 1.4 million hospitalizations each year (*1, 2*). Young children, adolescents, older adults above 65 years of age and immunocompromised individuals are particularly vulnerable, as infection with IBV can lead to severe illness and death, frequently complicated by secondary bacterial pneumonia and cardiovascular events (*3*). IBV infection accounts for 16–52% of all influenza-associated paediatric deaths, underscoring its significant public health relevance (*4*).

Influenza B virus was first identified in 1940 and by the 1970s had diverged into two distinctive genetic lineages, B/Victoria and B/Yamagata (*5–7*). Among the two lineages, the B/Victoria lineage exhibited stronger seasonal fluctuations and greater genetic drift, whereas B/Yamagata showed weaker genetic diversification. IBV evolves at relatively rapid evolutionary rates, comparable to influenza A, accumulating mutations that drove the emergence and expansion of multiple clades between 2000 and 2015 (*6, 7*). Prior to 2020, rapid and continual evolution of B/Victoria lineage viruses triggered substantial global outbreaks associated with the emergence of genetic variants carrying 2- and 3-amino acid (aa) deletions in the hemagglutinin (HA) protein. These mutations gave rise to multiple co-circulating clades (V1A.1–V1A.4) as we previously described (*8*). Clade V1A.1 is characterized by 2aa deletion (2aaΔ) at HA residues 162–163, together with three unique HA substitutions (D129G, I180V and R498K), whereas the remaining clades carried a 3aa deletion (3aaΔ) at HA residues 162–164 (*8*). During this period, the 2aaΔ variants replaced ancestral B/Victoria viruses lacking this deletion and rapidly became predominant globally, while the 3aaΔ variants circulated at lower frequencies. Consequently, an update of the B/Victoria influenza vaccine strain from B/Brisbane/60/2008 (no HA deletion) to B/Colorado/06/2017 (2aaΔ variant) was recommended. The COVID-19 pandemic ultimately led to the global extinction of B/Yamagata (*9*), while B/Victoria re-emerged and resumed widespread circulation of Clade V1A.3 (3aaΔ) variants, coinciding with a sharp population decline and complete disappearance of Clade V1A.1 (2aaΔ) variants, indicative of a global selective sweep.

Understanding the biological effects of gene mutations is fundamental to deciphering influenza virus evolution, transmission dynamics and by extension, vaccine strain selection. Substitutions in the HA protein can modulate viral infectivity, replication and pathogenicity; however, most mechanistic studies have focused on influenza A viruses with zoonotic potential at the animal-human interface (*10–13*). The functional impact of these HA deletions in B/Victoria lineage and how they shaped global epidemiology and transmission remain poorly understood. Here, we explored the virological impact of HA mutations in B/Victoria V1A.1 (2aaΔ) and V1A.3 (3aaΔ) viruses. Using robust reverse genetics system, we generated 19 recombinant viruses and systematically characterized the effects of single and combined mutations using *in vitro* and *in vivo* systems. Our integrated phylodynamic and virological analyses show that specific combinations of HA deletions and substitutions that markedly amplified viral replication, virulence, and pathogenesis and altered antigenicity. These genetic configurations underpin the increased fitness of emerging B/Victoria variants and provide a mechanistic explanation for the recent global sweep, accelerated diversification and increased severity of Clade V1A.3 viruses.

## RESULTS

### Global epidemiology and population dynamics of B/Victoria lineage

Since 2017, global influenza B epidemics have been driven by the rapid emergence and co-circulation of B/Victoria 2aaΔ and 3aaΔ variants (Fig. 1A). The 2aaΔ variant, carrying a 2-amino-acid deletion along with unique HA substitutions D129G, I180V and R498K, was first detected on 07 October 2016 (B/New York/52/2016) and gave rise to Clade V1A.1. The estimated mean time to most recent common ancestor (TMRCA) was August 2016 (95% HPD: 2016.33–2016.92; table S1). This variant predominated during 2017–2018 (Fig. 1A, denoted by vertical blue dotted line), with overall genetic diversity peaking in late 2016 to early 2017 (Fig. 1B), and continued circulating globally until March 2020, when influenza transmission sharply declined due to control measures implemented during the COVID-19 pandemic.

**Fig. 1.**
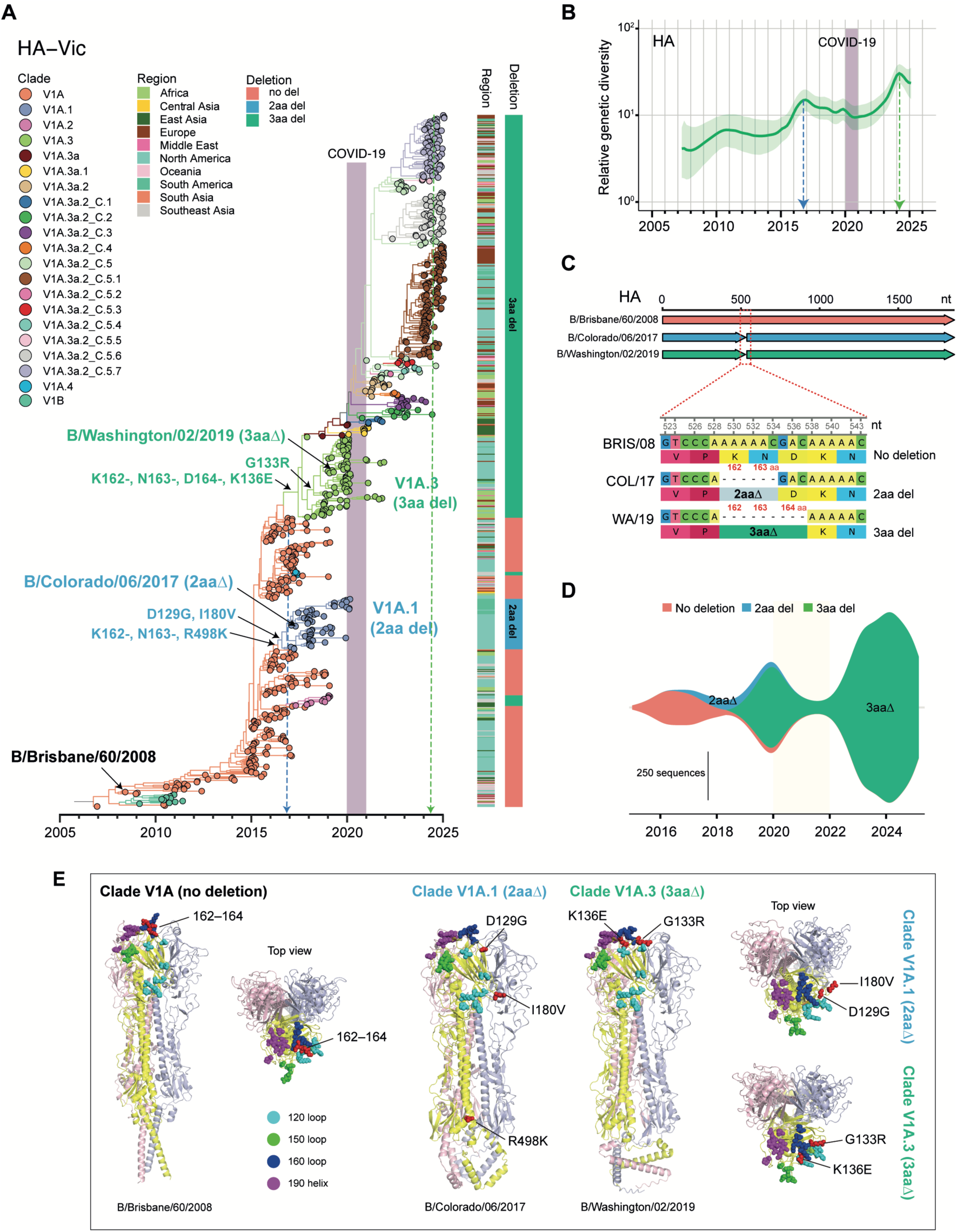
Global epidemiology and phylogenetic inference of influenza B/Victoria viruses. (**A**) Maximum clade credibility tree of HA gene sequences inferred using a Bayesian Gaussian Markov Random model in BEAST, based on globally subsampled viruses. Coloured branches representing genetic clades. Two-amino-acid and three-amino-acid deletions, along with major substitutions are highlighted for Clade V1A1 (2aaΔ) and V1A.3 (3aaΔ), as denoted by blue and green branches, respectively. Purple shading denotes the COVID-19 pandemic period. Blue and green vertical dotted lines indicate epidemic peaks. (**B**) Temporal changes in relative genetic diversity in B/Victoria based on HA gene sequences. (**C**) Comparison of HA sequences among three strains – ancestral B/Brisbane/60/2008 and two deletion variants represented by B/Colorado/06/2017 (Clade V1A.1), and B/Washington/02/2019 (Clade V1A.3) – illustrating characteristic two-amino-acid (positions 162–163) and three-amino-acid (162–164) deletions. (**D**) Global prevalence of ancestral and deletion variants based on all available sequences from 2016–2024. (**E**) Predicted structures of HA proteins from different B/Victoria clades. Side and top views of trimeric HA are shown for B/Brisbane/60/2008 (Clade V1A, no deletion), B/Colorado/06/2017 (Clade V1A.1, 2-aa deletion) and B/Washington/02/2019 (Clade V1A.3, 3-aa deletion). Deleted amino acid residues at 162–164 and clade-defining substitutions are highlighted as red spheres.

B/Victoria viruses re-emerged following the relaxation of COVID-19 pandemic restrictions; however, Clade V1A.1 was no longer detected, while Clade V1A.3 3aaΔ viruses with a K136E substitution in the HA – and subsequent acquisition of G133R – underwent rapid lineage expansion (indicated by vertical green dotted line in Fig. 1A). The estimated mean TMRCA of Clade V1A.3 is July 2017 (2017.50; 95% HPD: 2023.79–2024.71; table S1), with genetic diversity peaking in March 2024 (Fig. 1B), surpassing previous years (Fig. 1B). Clade V1A.3 viruses likely originated in Africa, notably in Mali, Nigeria and Sierra Leone, followed by multiple independent introductions into North and South Americas, Europe, East Asia (e.g. China) and Southeast Asia (e.g. Singapore) (Fig. 1A). To combat these evolving dynamics, the World Health Organization (WHO) updated the B/Victoria vaccine component from B/Colorado/06/2017 (COL/17, Clade V1A.1 2aaΔ at residues 162–163) Northern Hemisphere season to B/Washington/02/2019 (WA/19, Clade V1A.3 3aaΔ at 162–164) for the 2020–2021 season (Fig. 1C). Since 2023, Clade 1A.3 viruses further diversified into multiple co-circulating subclades and remain globally predominant (Fig. 1D).

To compare the HA structures of B/Victoria Clade V1A.1 and V1A. 3 viruses relative to B/Brisbane/60/2008, we applied AlphaFold to predict HA conformations of successive vaccine strains (Fig. 1E). The predicted HA structures of Clade V1A.1 COL/17 and Clade V1A.3 WA/19 showed substantial divergence from B/Brisbane/60/2008 (Clade V1A), with pairwise root mean square deviation (RMSD) values exceeding 20 Ångström (Å). In contrast, the HA structures of the two more recent clades were highly similar to each other, with a pairwise RMSD of ∼0.65 Å. Notably, deletions at residues 162–164 within the 160-loop are located proximal to the sialic acid receptor-binding site (Fig. 1E), a well-known structural hotspot for antigenicity and antibody escape.

To characterize how these HA deletions and associated substitutions influence viral phenotype and virulence, we first generated recombinant B/Victoria wild-type and mutant viruses (7+1) using an eight-segment reverse genetics system. Wild-type (WT) virus was rescued from all eight gene segments of the ancestral B/Brisbane/60/2008 strain (rgBRIS/08, denoted as Mutant 1 in Table 1), which lacks deletions. Mutant viruses (7+1) contained seven BRIS/08 segments and the HA segment of COL/17 (Clade V1A.1 2aaΔ, Mutant 2) or WA/19 (Clade VIA.3 3aaΔ, Mutant 18). In total, a panel of 19 recombinant viruses with different combinations of mutations was generated for subsequent experiments, albeit with four that were non-viable, likely due to incompatibility (Tables 1 and 2).

**Table 1.** Characterisation of B/Victoria V1A.1 mutant viruses with single and combinatory HA mutations.

| ID | Recombinant virus | Strains | Amino acid substitutions (HA) |  |  |  |  | Lung viral titers<br>(Log<sub>10</sub>TCID<sub>50</sub>/g) |  | BRIS/08<br>antiserum<br>HI titer |
| --- | --- | --- | --- | --- | --- | --- | --- | --- | --- | --- |
|  |  |  | 162 | 163 | 129 | 180 | 498 | Day 3 | Day 5 |  |
| 1 | rgBRIS/08 wild-type | B/Brisbane/60/2008 | K | N | N | I | R | 3.01±0.12 | 3.25±0.16 | 5120 |
| 2 | rgCOL/17 (2aaΔ) | B/Colorado/06/2017<br>HA | - | - | G | V | K | 5.44±0.20 | 4.9±0.12 | 320 |
| 3 | rgIns2aa | Single mutant | Ins2aa(162K/163N)* |  | G | V | K | 3.99±0.48 | 2.98±0.15 | 1280 |
| 4 | rg2aaΔ-129D | Single mutant | - | - | D* | V | K | x | x | x |
| 5 | rg2aaΔ-180I | Single mutant | - | - | G | I* | K | 4.40±0.24 | 3.15±0.08 | 1280 |
| 6 | rg2aaΔ-498R | Single mutant | - | - | G | V | R* | 4.73±0.16 | 3.29±0.35 | 2560 |
| 7 | rg2aaΔ-129D/180I | Double mutant | - | - | D* | I* | K | 5.73±0.22 | 5.26±0.24 | 160 |
| 8 | rg2aaΔ-180I/498R | Double mutant | - | - | G | I* | R* | 5.06±0.13 | 4.8±0.45 | 320 |
| 9 | rg2aaΔ-129D/498R | Double mutant | - | - | D* | V | R* | x | x | x |
| 10 | rgIns2aa/129D | Double mutant | Ins2aa(162K/163N)* |  | D* | V | K | 4.66±0.25 | 5.85±0.07 | 320 |
| 11 | rgIns2aa/180I | Double mutant | Ins2aa(162K/163N)* |  | G | I* | K | 4.56±0.24 | 3.3±0.25 | 1280 |
| 12 | rgIns2aa/498R | Double mutant | Ins2aa(162K/163N)* |  | G | V | R* | 4.00±0.14 | 4.19±0.21 | 1280 |
| 13 | rg2aaΔ-129D/180I/498R | Triple mutant | - | - | D* | I* | R* | 4.5±0.29 | 4.05±0.19 | 2560 |
| 14 | rgIns2aa/129D/180I | Triple mutant | Ins2aa(162K/163N)* |  | D* | I* | K | x | x | x |
| 15 | rgIns2aa/129D/498R | Triple mutant | Ins2aa(162K/163N)* |  | D* | V | R* | 5.41±0.18 | 3.95±0.43 | 2560 |
| 16 | rgIns2aa/180I/498R | Triple mutant | Ins2aa(162K/163N)* |  | G | I* | R* | 5.04±0.10 | 3.93±0.12 | 2560 |
| 17 | rgIns2aa/129D/180I/498R | 4-amino-acid mutant | Ins2aa(162K/163N)* |  | D* | I* | R* | x | x | x |

Abbreviations: \* denote changes in amino acid residues at a specific position in the HA gene of B/Colorado/06/2017, while x denote non-viable virus.

**Table 2.** Characterisation of B/Victoria V1A.3 mutant viruses with single and combinatory HA mutations.

| ID | Recombinant virus | Strains | Amino acid substitutions (HA) |  |  |  |  | Lung viral titers<br>(Log<sub>10</sub>TCID<sub>50</sub>/g) |  | BRIS/08<br>antiserum<br>HI titer |
| --- | --- | --- | --- | --- | --- | --- | --- | --- | --- | --- |
|  |  |  | 162 | 163 | 164 | 133 | 136 | Day 3 | Day 5 |  |
| 18 | rgWA/19 (3aaΔ) | B/Washington/02/2019<br>HA | - | - | - | R | E | 5.94±0.21 | 4.62±0.15 | 2560 |
| 19 | rgIns1aa-164D | Single mutant | - | - | Ins1aa(164D)* | R | E | 3.25±0.09 | 3.44±0.27 | 1280 |
| 20 | rgIns3aa | Double mutant | Ins3aa(162K/2163N/164D)* |  |  | R | E | 3.53±0.18 | 3.78±0.19 | 5120 |
| 21 | rg3aaΔ-133G | Single mutant | - | - | - | G* | E | 3.36±0.62 | 3.17±0.07 | 5120 |
| 22 | rg3aaΔ-136K | Single mutant | - | - | - | R | K* | 3.90±0.42 | 3.48±0.28 | 5120 |
| 23 | rg3aaΔ-133G/136K | Double mutant | - | - | - | G* | K* | 3.53±0.38 | 3.72±0.20 | 5120 |

Abbreviations: \* denote changes in amino acid residues at a specific position in the HA gene of B/Washington/02/2019.

### A two-amino acid deletion in HA of B/Victoria Clade V1A.1 viruses alters replication and virulence

To evaluate the virological impact of the two-amino-acid (2aaΔ) deletion in the HA protein of the B/Victoria Clade V1A.1 *in vivo*, a mutant virus with residues 162K and 163N reinserted into the HA of COL/17 was constructed (rgIns2aa, denoted as Mutant 3 in Table 1). Mice were intranasally inoculated with rgCOL/17-Ins2aa (Mutant 3) and compared with WT viruses rgBRIS/08 (no deletion) and rgCOL/17 (2aaΔ). All infected mice were monitored for weight loss and survival for 14 days post-infection (dpi), and lung viral titers were measured at days 3 and 5 post-infection (Fig. 2A, fig. S1).

**Fig. 2.**
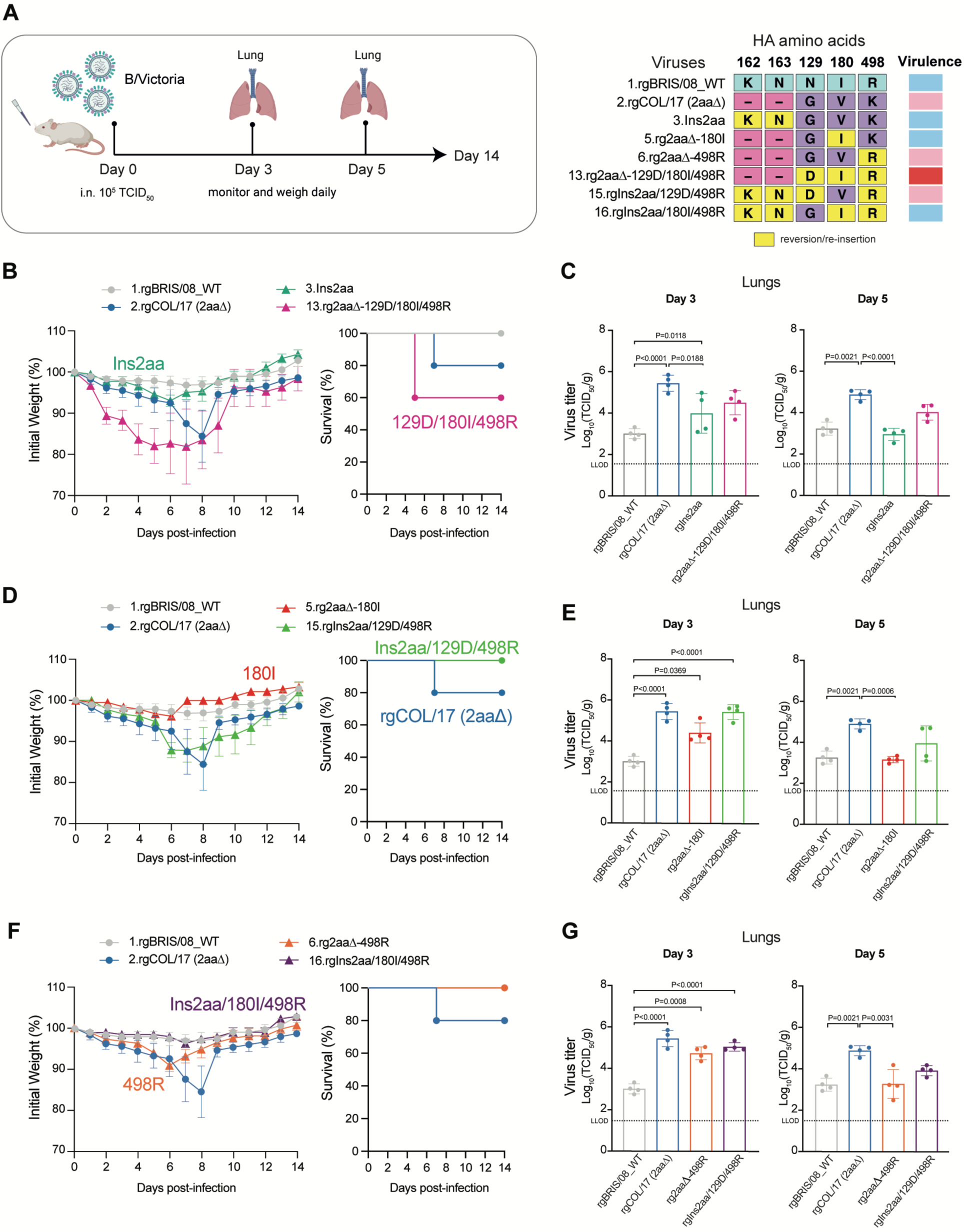
The effect of HA mutations on B/Victoria Clade V1A.1 viruses in mice. (**A**) Schematic diagram illustrating animal study design. Three- to five-week-old female BALB/c mice (n=13 per group) were intranasally inoculated with 10^5^ TCID_50_ of each virus. The diagrams were created with BioRender. The right panel shows a subset of recombinant B/Victoria viruses generated by reverse genetics. The wild-type BRIS/08 refers to ancestral B/Brisbane/60/2008 strain, with all gene segments derived from same strain, whereas the mutant viruses were generated using a 7+1 (HA) plasmid system. The rgCOL/17 virus (2aaΔ) carries the HA gene from the B/Colorado/06/2017 strain, representing Clade V1A.1 viruses with a two-amino-acid deletion at positions 162–163. Each recombinant virus contains single or combinational HA substitutions, with amino acid changes indicated at positions 162, 163, 129, 180, and 498. Yellow boxes highlight yellow re-insertion or reversion sites, and black dashes indicate amino acid deletions. Virulence levels are summarized in the right bars (light blue: low; pink: moderate and red: high). (**B, D, F**) Weight changes and survival of virus-infected mice (n=5 per group) were monitored daily for 14 days post-infection (dpi). (**C, E, G**) Lung tissues from infected mice (n=4 per group) were collected at 3 and 5 dpi, and viral titers were determined in MDCK cells using the Reed-Muench 50% endpoint, expressed as the log_10_TCID_50_/g. Mean and error bars are shown, and the lowest limit of detection is represented by dotted horizontal lines. Statistical significance was assessed using one-way ANOVA followed by Tukey test for body weight and viral titers and log-rank (Mantel-Cox) test for survival analysis. Data represent mean ± SEM from biological independent replicates (n=5 for body weight and survival; n=4 for viral titers).

Mice infected with rgBRIS/08 (WT) developed only mild disease, with stable body weight maintained throughout infection (Fig. 2B). Corresponding lung viral loads remained low, with titers of approximately 10^3^ TCID_50_/g at both 3 and 5 dpi (Fig. 2C). In contrast, infection with rgCOL/17, which carries the 2aa deletion along with 129G/180V/498K substitutions, led to progressive weight loss beginning at 2 dpi, increasing further to 15% of baseline by 7 dpi, with one of five mice succumbing (20% mortality) by 7 dpi (Fig. 2B). Lung viral titers were significantly elevated (∼10^5^ TCID_50_/g) at both 3 and 5 dpi (P<0.0001, Fig. 2C) compared with rgBRIS/08.

By comparison, mice infected with rgCOL/17-Ins2aa (reinserted residues 162–163 highlighted by yellow boxes in Fig. 2A) exhibited minimal weight loss (±5%) and 100% survival, in stark contrast to the pronounced weight loss observed in rgCOL/17-infected mice. The disease course of rgCOL/17-Ins2aa infected mice closely resembled the mild phenotype of rgBRIS/08 infection, with slight weight loss appearing at 6 dpi and full recovery to baseline by 10 dpi (Fig. 2B). Viral replication of rgCOL/17-Ins2aa was significantly lower at 3 dpi (P=0.0188) and 5 dpi (P<0.0001) compared with rgCOL/17 (Fig. 2C). These results indicate that re-insertion of residues 162–163 substantially attenuates the virulence of the B/Victoria Clade V1A.1 virus in mice.

To further assess the specific effect of the 2aa deletion, we generated Mutant 13 (rg2aaΔ-129D/180I/498R), in which three substitutions (G129D, V180I and K498R) were reverted to their pre-emergent residues in the rgCOL/17 HA while retaining the 2aa deletion (Fig. 2A). Infection with Mutant 13-rgCOL/17 induced 5% weight loss in mice at 1 dpi and 20% by 6 dpi, substantially higher compared with the more delayed weight loss observed in rgCOL/17 (Fig. 2B). By 5 dpi, two of the five mice met human endpoints, while the remaining three recovered by 9 dpi, resulting in an overall mortality rate of 40% (Fig. 2B). Lung viral titers at both 3 and 5 dpi were higher those of rgBRIS/08 WT but lower than rgCOL/17 (Fig. 2C). These findings demonstrate that the 2aa deletion in HA plays a critical role in modulating B/Victoria virulence *in vivo*, but in a context dependent manner. Re-insertion of the deleted residues attenuated viral replication and disease severity, whereas the retention of the deletion coupled with the three historical substitutions (129D/180I/498R) increased viral replication and mortality, highlighting the synergistic contribution of these HA changes.

We also evaluated the antigenic properties of these mutant viruses relative to rgCOL/17 and wild-type viruses using hemagglutination inhibition (HI) assays against BRIS/08 antiserum. rgCOL/17 (2aaΔ) virus exhibited a 16-fold reduction in HI titer (1:5120 to 1:320) compared with rgBRIS/08 WT when tested against BRIS/08 antisera (Table 1), consistent with expected antigenic drift. Mutant 3 (rgIns2aa), Mutant 5 (180I) and 6 (498R) – each showed at least 2-fold less reduction (1:5120 to 1:2560) compared with rgCOL/17, indicative of a partial restoration toward BRIS/08-like antigenicity. Further reversion to ancestral residues in Mutant 13 (129D/180I/498R) resulted in an antigenic profile more closely resembling rgBRIS/08 than rgCOL/17 (HI titer 1:2560). These findings suggest that both the 2aa deletion and specific mutations impacted the antigenic relationship of rgBRIS/08 and rgCOL/17 viruses.

### Reversion to V180I or K498R mutation attenuates virus virulence

We investigated the contributions of additional HA substitutions in COL/17 (2aaΔ) by introducing single reversions (G129D, V180I, K498R), generating three mutant viruses (Mutants 4–6, Table 1). Mutant 4, carrying only the G129D reversion, was non-viable, indicating that the 2aa deletion is incompatible with the ancestral 129D residue and that 129G is essential for maintaining the viability of the 162–163 deletion variant. Mutant 5, with the 180I reversion, caused minimal weight loss (±5%) and 100% survival (Fig. 2D) with a corresponding ≥10-fold reduction in lung viral titers at both 3 and 5 dpi, significantly lower than rgCOL/17 at 5 dpi (P=0.0006, Fig. 2E). In contrast, Mutant 15, which carries 129D/498R substitutions along with re-insertion of residues 162–163, induced only transient weight loss (5% at 6 dpi), with all mice surviving 14 days (Fig. 2D). Lung viral replication for Mutant 15 was comparable to rgCOL/17 at 3 dpi but ∼10-fold lower at 5 dpi, and titers were ∼20-fold higher at 3 dpi and ∼10-fold higher at 5 dpi relative to rgBRIS/08 WT (Fig. 2E). These results indicate that the I180V residue in rgCOL/17 contributes to enhanced virulence, while reversion to 180I attenuates pathogenesis, as reflected by reduced viral replication and complete survival in Mutant 5-infected mice.

Similarly, Mutant 6, carrying only the 498R substitution, caused a transient 10% weight loss at 6 dpi with all mice survived (Fig. 2F). Lung viral titers were initially high (∼10^5^ TCID_50_/g) at 3 dpi but declined to levels comparable to rgBRIS/08 WT (∼10^3^ TCID_50_/g) by 5 dpi (Fig. 2G). Mutant 16, combining the 180I/498R with the 2aa re-insertion, maintained 100% body weight, resembling rgBRIS/08 WT (Fig. 2F), with titers decreasing from ∼10^5^ to 10^4^ TCID_50_/g by 5 dpi (Fig. 2F, G). These results suggest that the historical K498R residue does not enhance pathogenicity in mice, as expected given its position in the HA stem region (Fig. 1E).

### Dual HA substitutions synergistically module disease severity and infection dynamics

We next assessed mutants carrying different combinations of amino acid substitutions *in vivo*. Mutant 7, harboring the 129D and 180I substitutions, induced significantly accelerated weight loss from 2 to 4 dpi compared with rgCOL/17 (P<0.005), with all mice succumbing by 5 dpi (P=0.039), compared with rgCOL/17 (Fig. 3A). Consistently, Mutant 7 exhibited enhanced lung viral replication at both 3 and 5 dpi, similar to rgCOL/17 titers (Fig. 3C). Mutant 10, which carries the 2aa re-insertion and 129D substitution, also caused accelerated weight loss (P<0.005) and increased mortality (P=0.0031), with higher lung viral titers at 5 dpi and complete mortality

**Fig. 3.**
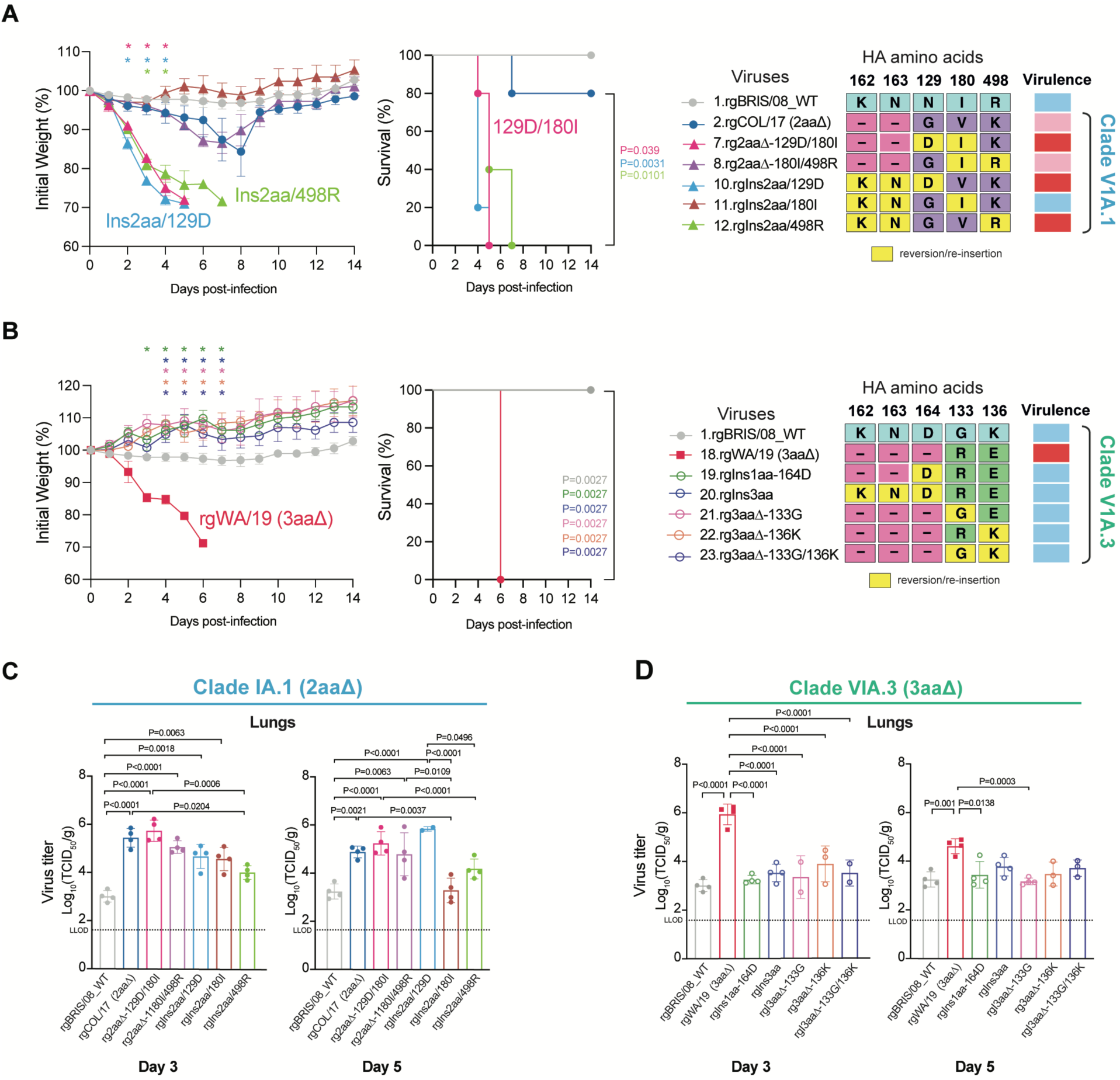
Weight loss and mortality of Clade V1A.1 and V1A.3 viruses in mice. Female BALB/c mice (3–5 weeks old, n=13 per group) were intranasally inoculated with 10^5^ TCID_50_ of each virus. (**A**) Body weight and survival of virus-infected mice (n=5 per group) were monitored daily for 14 days post-infection (dpi). The right panel shows a subset of recombinant B/Victoria Clade V1A.1 viruses generated by reverse genetics. Yellow boxes highlight re-insertion or reversion sites at amino acid positions 162, 163, 129, 180 and 498, and black dashes indicate amino acid deletions. Virulence levels are summarized in the right bars (light blue: low; pink: moderate and red: high). (**B**) Body weight and survival of mice (n=5 per group) infected with Clade V1A.3 viruses, including rgWA/19 (3aaΔ) and mutant viruses carrying clade-specific amino acid changes at positions 162, 163, 164, 133, and 136. The rgWA/19 virus (3aaΔ) carries the HA gene from the B/Washington/02/2019 strain, representing Clade V1A.3 viruses with a three-amino-acid deletion at positions 162–164. (**C, D**) Lung tissues from infected mice (n=4 per group) were collected at 3 and 5 dpi. Statistical significance was assessed using one-way ANOVA with Tukey’s post hoc test for body weight and viral titers, and the log-rank (Mantel-Cox) test for survival analysis. The lowest limit of detection is represented by dotted horizontal lines. Data represent mean ± SEM from biological independent replicates (n=5 for body weight and survival; n=4 for viral titers). *P< 0.05 indicates significance.

(Fig. 3A, C), highlighting the contribution of these mutations to virulence. Mutant 12, containing the 2aa re-insertion and 498R substitution, induced accelerated weight loss between 2–4 dpi (P<0.01) and increased mortality (P=0.01), with lung titers of ∼10^4^ TCID_50_/g at 3 dpi that remained stable at 5 dpi, indicating sustained viral replication (Fig. 3A, C).

In contrast, Mutant 8 (180I/498R) and Mutant 11 (Ins2aa/180I) led to attenuated disease compared with rgCOL/17 (Fig. 3A). Mutant 8 maintained lung titers of ∼10^5^ TCID_50_/g at both 3 and 5 dpi, whereas Mutant 11 reached ∼10^4.5^ TCID_50_/g at 3 dpi but declined 15-fold to ∼10^3^ TCID_50_/g by 5 dpi (Fig. 3C). Most double mutant viruses showed reduced HI titer (16- to 32-fold difference) reactivity relative to homologous rgBRIS/08 WT titers, similar to rgCOL/17 (Table 1). Together, these results indicate that the G129D substitution, especially when combined with V180I or the 2aa re-insertion, enhances virulence and accelerates mortality, whereas K498R combined with V180I attenuates viral replication. As such, dual substitutions (G129D/V180I or Ins2aa/G129D) markedly alter disease severity and infection outcomes *in vivo*.

### HA K136E and the 3-amino-acid deletion in Clade V1A.3 viruses increase mortality in mice

We further compared Clade V1A.3 (rgWA/19) viruses, which harbors a 3-amino-acid deletion and K136E substitution, with Clade V1A.1 (rgCOL/17) and ancestral rgBris/08 WT strains.

Mice infected with rgWA/19 exhibited markedly greater disease severity, with early weight loss (∼7%) at 2 dpi that rapidly progressed to >75% by 6 dpi (P<0.05) and humane endpoints in all animals (Fig. 3B). Consistently, lung viral titers were elevated, reaching ∼10^6^ TCID_50_/g at 3 dpi and remaining ∼10^5^ TCID_50_/g at 5 dpi (Fig. 3D).

To dissect the contributions of HA mutations in WA/19, six different mutants (Mutant 18–23), with single, double or triple amino acid changes were evaluated (Table 2). Mutant 19-rgWA/19, with re-insertion at residue 164, caused no obvious weight loss (Fig. 3B) and displayed markedly attenuated replication, with lung titers reduced >1000-fold at 3 dpi (P<0.0001) and 100-fold at 5 dpi (P<0.0138), relative to rgWA/19 (Fig. 3D). Similarly, Mutant 20 with re-insertion at residues 162–164, also exhibited an attenuated phenotype, with body weight unchanged throughout the infection and lung titers ∼10-fold lower than rgWA/19 at both 3 dpi (P<0.0001) and 5 dpi (Fig. 3D). Likewise, Mutants 21–23, carrying 133G, 136K, or combined 133G/136K substitutions, induced minimal disease and substantially reduced viral replication in infected mice, with viral titers at least 20-fold lower at 3 dpi and 10-fold lower at 5 dpi compared with rgWA/19 (Fig. 3D). As such, the 3-amino acid deletion at residues 162–164, together with the K136E and G133R substitutions, are critical for enhancing viral replication and virulence compared with the preceding B/Victoria viruses.

### Clade 1A.3 viruses induce elevated pulmonary inflammation in vivo

Longitudinal non-invasive positron emission tomography/computed Tomography (PET/CT) imaging with ^18^F-Fluorodeoxyglucose (^18^F-FDG) radiotracer was employed to assess and compare the extent of lung inflammation and metabolic activity in mice infected with two influenza B clades: Clade V1A.1 (2aaΔ) and Clade V1A.3 (3aaΔ). ^18^F-FDG is a glucose analog irreversibly accumulated in highly metabolic tissues, including inflammatory lesions. Mice were intranasally infected with rgCOL17 or rgWA/19 and subjected to by PET/CT imaging at days 3 and 5 post-infection (Fig. 4A). Maximum intensity projection (MIP) of ^18^F-FDG-PET/CT images showed higher ^18^F-FDG uptake in the lungs of infected mice compared to pre-infection, with foci of increased update indicative of inflammatory lesions (indicated by white arrows in Fig. 4B). Notably, rgWA/19 (3aaΔ) infection had a higher level of lung inflammation at both 3 and 5 dpi than those of rgCOL/17 (2aaΔ) infected mice (Fig. 4C), consistent with the higher mortality observed in rgWA/19-infected mice (100%) versus rgCOL/17-infected mice (20%) in the independent experiments described above. Regions of elevated ^18^F-FDG uptake on PET images corresponded closely with CT-defined lung lesions (white dashed outlines in Fig. 4D). Quantification of standardized lung ^18^F-FDG uptake – reflecting lung metabolic activity – showed the greatest tracer accumulation over time in rgWA/19-infected mice (3aaΔ) (Fig. 4E). By 5 dpi, lung ^18^F-FDG uptake in rgWA/19-infected mice was 1.9-fold higher than pre-infection (*p* = 0.04) and 2.5-fold higher than in rgCOL/17-infected mice (*p* = 0.009) (Fig. 4E). Moreover, uptake in rgWA/19-infected mice increased by 1.4-fold from 3 dpi to 5 dpi (Fig. 4E, table S2). These observations were broadly concordant with metabolic lung volume measurements (MLV, Fig. 4F). At 3 dpi, rgWA/19-infected mice exhibited a 2.3-fold increase relative to pre-infection (*p* = 0.03), with a further marked increase by 5 dpi (13.6-fold, *p* = 0.008, fig. S2). Conversely, MLV in rgCOL/17 remained relatively unchanged (fig. S2). These data suggest that rgWA/19 (3aaΔ) results in elevated metabolic activity and inflammation in the lungs.

**Fig. 4.**
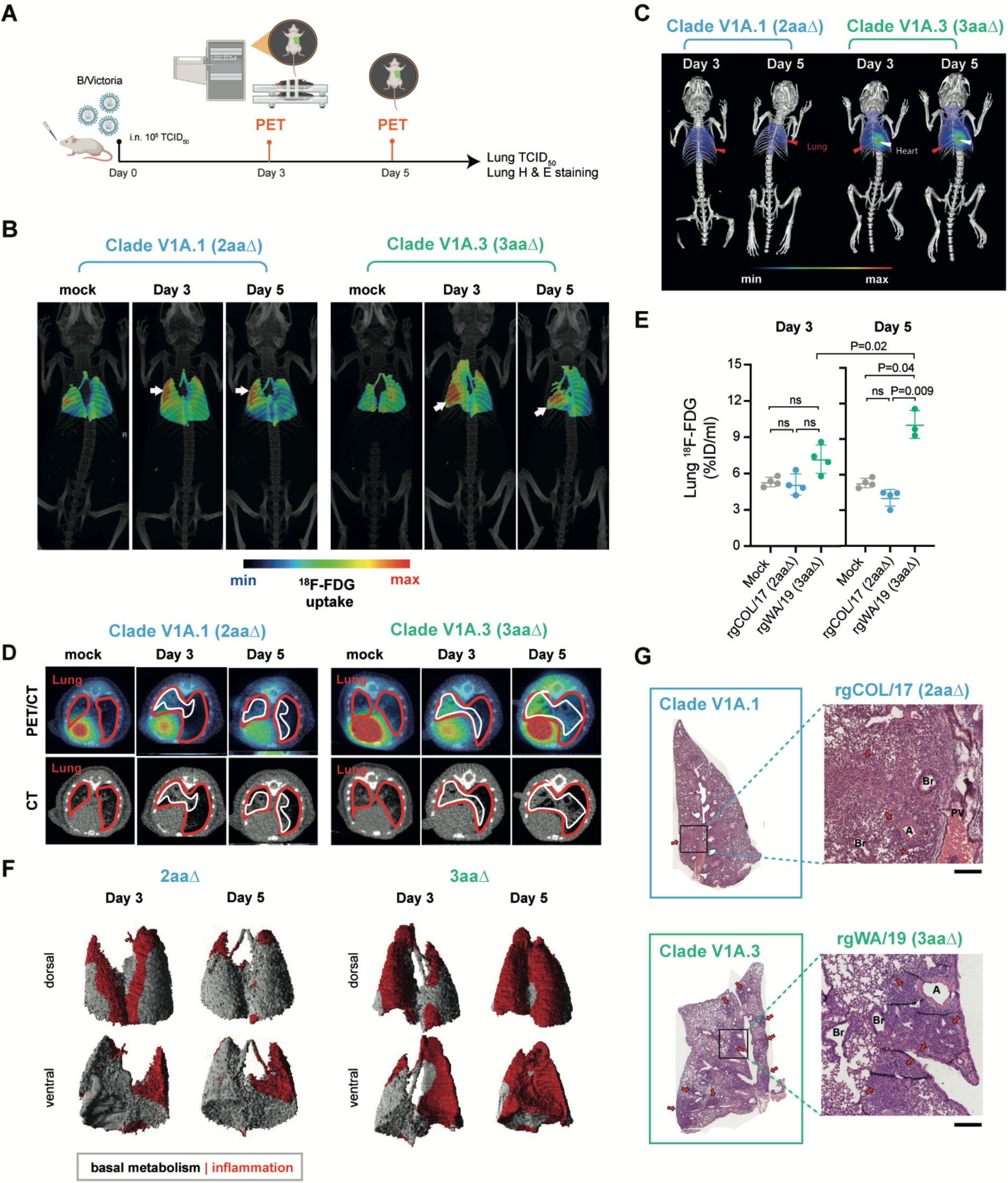
Comparison of lung inflammation and pathology between Clade V1A.1 and V1A.3 viruses using longitudinal PET/CT imaging. **(A)** Schematic diagram illustrating independent mouse experiments for PET/CT imaging, created with BioRender. Mice were intranasally infected with rgCOL/17 (Clade V1A.1) or rgWA/19 (Clade V1A.3) viruses and subjected to ^18^F-FDG -PET/CT scans on days 3 and 5 post-infection (dpi). (**B, C**) Maximum intensity projection (MIP) images of ^18^F-FDG-PET/CT scans. Foci of sustained pulmonary ^18^F-FDG uptake are shown with white arrows. (**D**) Transverse image slices of PET/CT overlay and CT images highlighting areas of elevated lung uptake (outlined by red solid lines). Pulmonary lesions seen in lung CT images (outlined in dashed white lines) colocalize with intense ^18^F-FDG uptake in PET images (outlined in white solid lines). (**E**) Quantification of lung ^18^F-FDG uptake in pre-infection, rgCOL/17- and rgWA/19-infected mice. (**F**) Surface-rendered 3D reconstructions of metabolic lung volumes (MLV) based on a baseline threshold (6.65% ID/ml) from pre-infection. Grey regions indicate baseline ^18^F-FDG uptake, while red regions represent areas of elevated ^18^F-FDG activity. (**G**) Hematoxylin and eosin-stained sections of the left lung lobe collected at 5 dpi, with red arrows indicating inflammatory foci. Selected areas are shown at higher magnification, highlighting alveolar and airway structures, including artery (A), bronchiole (Br), and pulmonary vein (PV). Scale bars = 1mm (whole lung lobe) and 200 μm (enlarged regions). Data for 3 and 5 dpi are presented as mean ± SEM, with individual data points representing individual biological replicates. Statistical analysis was performed using Welch’s ANOVA or Mann-Whitney test, as indicated. ns, not significant.

Histopathological examination on 5 dpi further indicated that mice infected with rgCOL/17 induced mild inflammation, with largely preserved lung architecture and few inflammatory lesions (Fig. 4G). In contrast, rgWA/19 infection in mice induced markedly severe pathology involving widespread loss of normal alveolar structure, dense inflammatory infiltrates, and extensive filling of alveoli with fluid and cellular debris (Fig. 4G, fig. S3 and S4).

Immunofluorescence staining of the lungs for IBV NP and CD45^+^ cells revealed exacerbated virus infection and infiltration of inflammatory cells into the lungs of rgWA/19-mice compared to the other groups (fig. S5). These findings indicate that Clade V1A.3 viruses (3aaΔ) elicit more robust inflammatory response and more extensive pulmonary damage than Clade V1A.1 viruses (2aaΔ).

### B/Victoria deletion variants show robust *in vitro* replication on α2,3- and α2,6-sialylated glycans

In addition to animal infection experiments, we evaluated the effects of B/Victoria lineage HA mutations on virus replication *in vitro* and receptor binding profiles. MDCK cells were infected with wild-type or mutant B/Victoria viruses (MOI=0.01) representing Clade V1A.1 and V1A.3 viruses, and viral titers were quantified over a 72-hour time course (Fig. 5). rgCOL/17 (Clade V1A.1; 2aaΔ) virus replicated efficiently in MDCK cells and reached titers of up to ∼10^6^ TCID_50_/ml as early as 48 hours-post infection (hpi); in comparison, wild-type rgBRIS/08 had delayed replication kinetics although reaching similar peak titers by 72hpi (Fig. 5A).

**Fig. 5.**
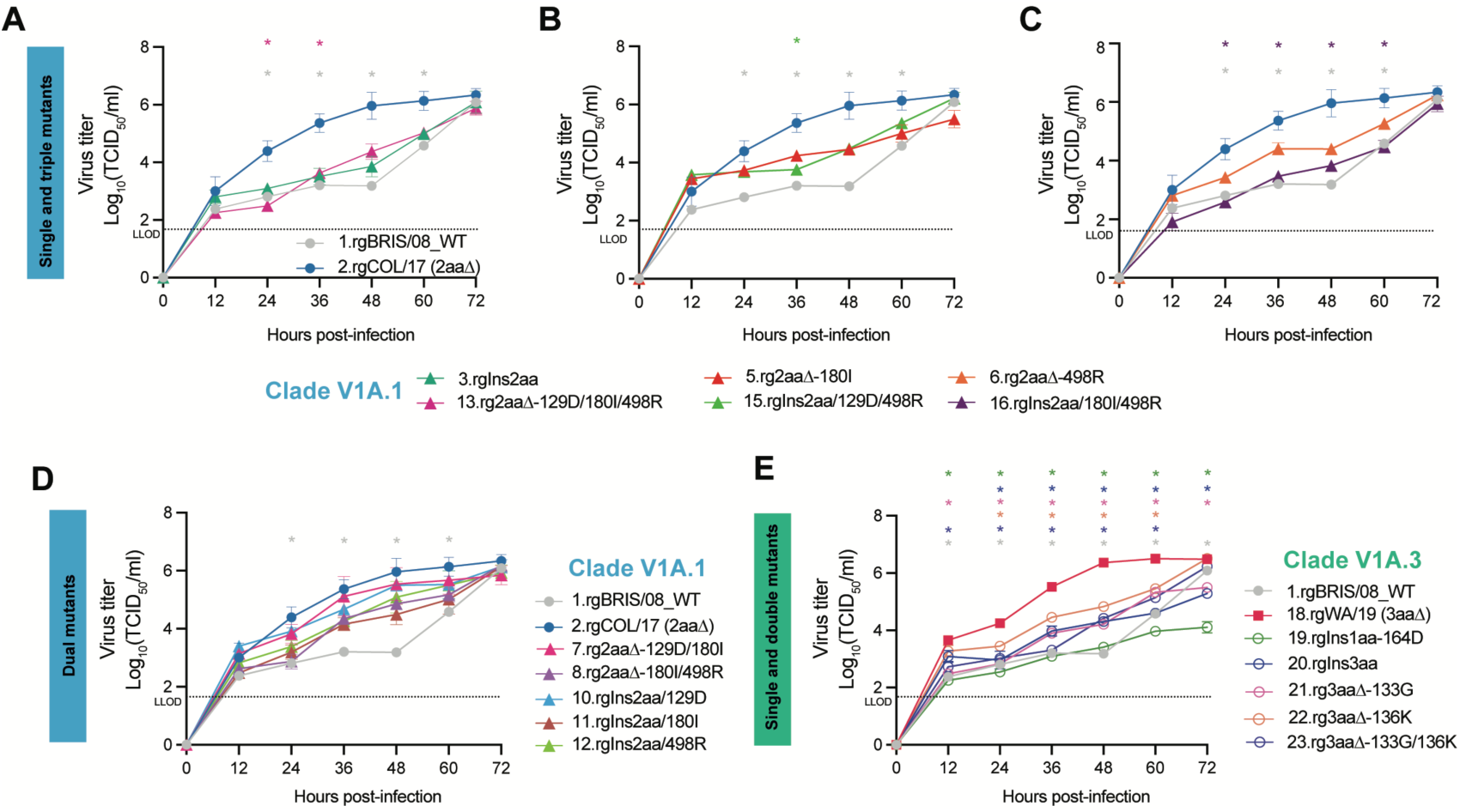
HA substitutions differentially modulate *in vitro* replication kinetics of Clade V1A.1 and V1A.3 viruses. Replication kinetics of recombinant B/Victoria viruses were assessed in MDCK cells infected at a multiplicity of infection (MOI) of 0.01. Viral titers were quantified at 12, 24, 36, 48, 60, and 72 hours post-infection using the Reed-Muench 50% endpoint and expressed as log_10_TCID_50_/ml. Panels show replication profiles of (**A, B, C**), single and triple mutants of Clade V1A.1. (**D**) dual mutants of Clade V1A.1, and (**E**) mutants of Clade V1A.3 viruses. Statistical significance was assessed using one-way ANOVA with Tukey’s post hoc test for viral titers. *P<0.05. Data represent mean ± SEM from biological independent replicates (n=3).

Mutants 3 (rgIns2aa), 13 (rg2aaΔ-129D/180I/498R) and 16 (rgIns2aa/180I/498R) exhibited slower replication kinetics like rgBRIS/08 WT, while Mutants 5, 6 and 15 showed intermediate kinetics of up to ∼10^5^ TCID_50_ by 72 hpi (Fig. 5A–C). As such, single and triple mutants of Clade V1A.1 carrying residues re-insertions at the 2aa deletion site or individual substitutions (G129D, V180I, K498R) replicated less efficiently than rgCOL/17 (Fig. 5B, C). However, double mutants (e.g. Mutant 7 with 129D/180I) achieved titers comparable to ∼10^6^ TCID_50_ by 72 hpi (Fig. 5D).

The rgWA/19 (Clade V1A.3; 3aaΔ) virus also replicated efficiently, reaching ∼10^4^ TCID_50_ by 12 hpi and peaking at ∼10^6^ TCID_50_ by 48 hpi (Fig. 5E). However, most Clade V1A.3 mutants (Mutants 18–23) showed reduced replication kinetics relative to rgWA/19, except Mutants 22 and 23 which reached comparable peak titers (Fig. 5E), suggesting partial restoration of replicative fitness due to these mutations.

To further evaluate receptor binding, we tested the virus panel against glycan binding assays. Recombinant viruses including the wild-type virus bound efficiently to α2,3-linked and α2,6-linked sialylated glycans (Fig. 6, fig. S6). Relative to rgCOL/17, Mutants 6 (498R), 12 (Ins2aa/498R), 13 (129D/180I/498R) and 16 (Ins2aa/180I/498R) displayed relatively weaker binding to both α2,3-linked and α2,6-linked at 9 ug/ml (Fig. 6A), however, the differences did not reach significance. Among all the Clade V1A.1 viruses, Mutant 10 (Ins2aa/129D) also showed lower binding to both receptors, whereas Mutant 15 (Ins2aa/129D/498R) exhibited the strongest 6’SLN avidity (Fig. 6B).

**Fig. 6.**
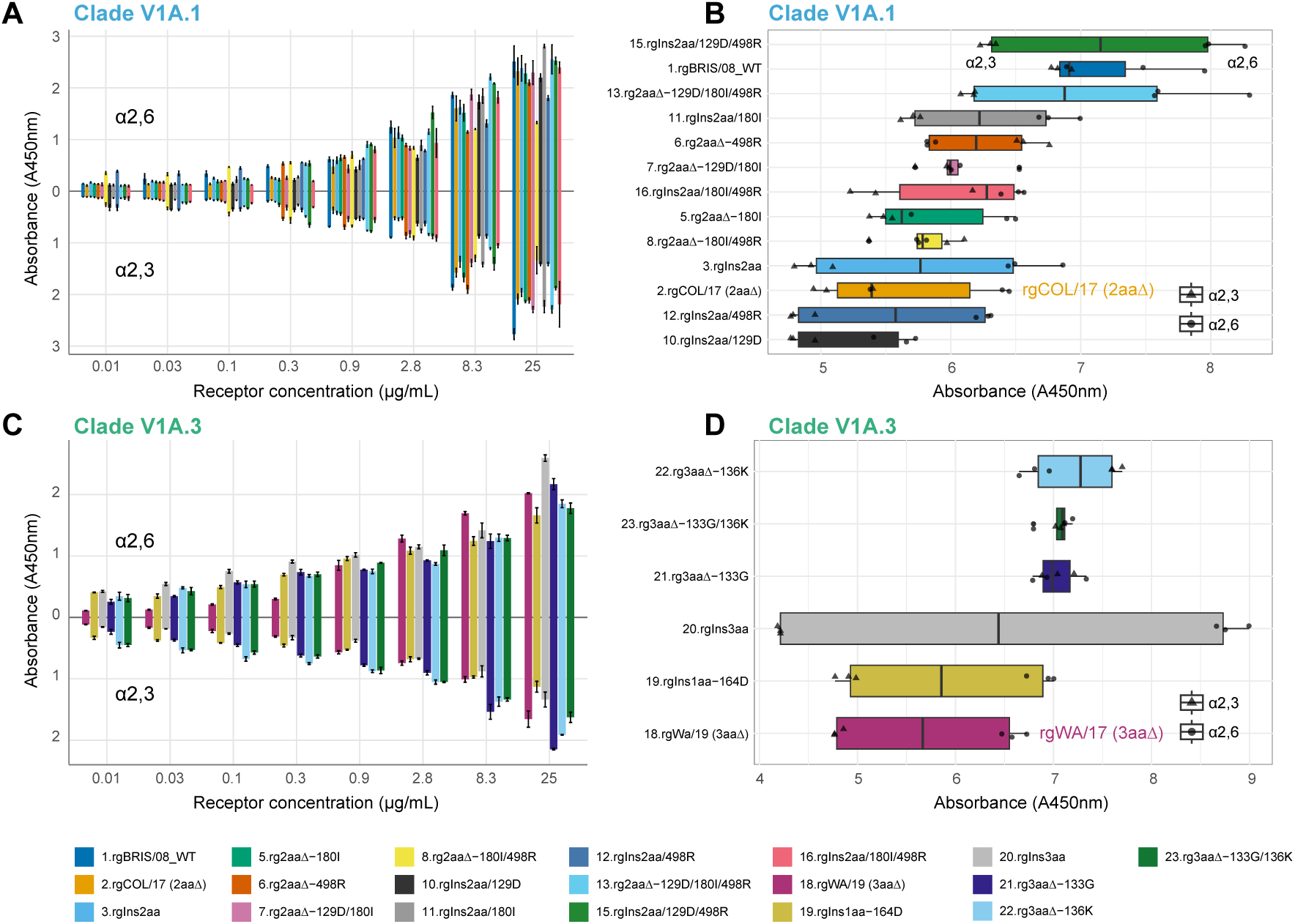
Receptor binding profiles of recombinant viruses. Virus binding to α2,6-linked (6’SLN-PAA) and α2,3-linked (3’SLN-PAA) sialyglycopolymers was assessed by solid-phase direct binding assays. Serial dilutions of biotinylated glycans were incubated with equal amounts of virus at 64 HA units (HAU). Bound virus was detected using HRP-conjugated streptavidin and TMB substrate, and absorbance was measured at 450 nm. (**A, C**) Binding profiles of Clade V1A.1 and Clade V1A.3 recombinant viruses, with absorbance values above and below the x-axis corresponding to α2,6-linked and α2,3-linked glycans, respectively. Vertical bar plots indicate mean absorbance values, with error bars indicating standard deviation. (**B, D**) Horizontal boxplots summarize total absorbance across all glycan concentrations for each biological replication (n=3), with individual points overlaid (circles, α2,6-linked; triangles, α2,3-linked). Statistical significance was determined using one-way ANOVA.

All Clade V1A.3 (3aaΔ) viruses also bound efficiently to α2,3-linked and α2,6-linked sialylated glycans, even at a lower glycan concentration of 0.9 μg/ml (Fig. 6C). Mutants 21 (133G), 22 (136K) and 23 (133G and 136K) displayed enhanced α2,3-linked binding, compared with rgWA/19 (Fig. 6D). Interestingly, mutants 20 (Ins3aa) showed the greatest difference in binding between α2,3-linked and α2,6-linked glycans, with a preference for α2,6 (Fig. 6D). As such, all B/Victoria viruses efficiently bound to both α2,3-linked and α2,6-linked glycans, indicating broad receptor susceptibility that likely supports their sustained circulation in human populations.

### HA deletions and substitutions differentially influence pH and thermal stability

We next assessed the pH and temperature stability of the wild-type and mutant viruses. rgBRIS/08 WT displayed lower tolerance to acidic conditions at pH 5.0 (Fig 7a; P=0.027, table S8) and reduced thermal stability compared with rgCOL/17 (Fig. 7C). rgCOL/17 exhibited a loss of infectivity only at pH 5.0 (Fig. 7A, P<0.0001, table S12) and maintained its initial hemagglutinin activity (64 HA units) across temperatures from 33°C to 45°C (Fig. 7C). These observations were consistent with high-temperature stability assays in which viruses were incubated at a fixed temperature of 50°C for 12 hours: rgCOL/17 retained 16 HA units for up to 6 hours before activity declined (fig. S8A). In contrast, Clade V1A.1 mutants carrying single or triple substitutions were less thermally stable, showing a marked decline in HA activity to 8 units after 5 hours at 50°C (fig. S8B–D).

**Fig. 7.**
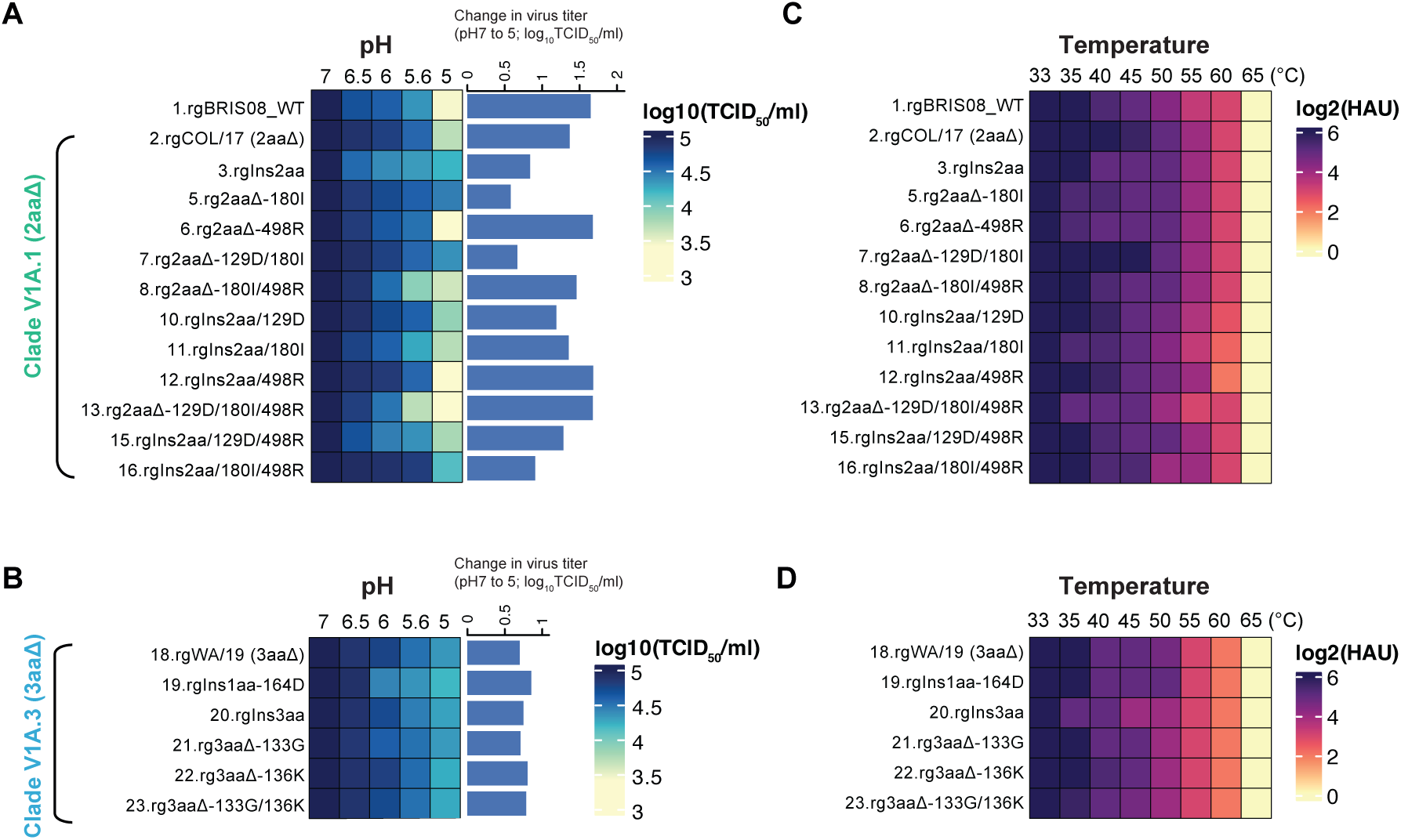
Acidic and thermal stability profiles of recombinant B/Victoria viruses. (**A**) Acid stability of Clade V1A.1 recombinant viruses. Viruses were incubated in buffers with pH ranging from pH 5.0 to 7.0 and subsequently used to infect MDCK cells. At 72 hours post-infection (hpi), cells were fixed and stained for NS1 protein, and viral titers were quantified by TCID_50_ assay. (**B**) Acid stability of Clade V1A.3 recombinant viruses, assessed as described for panel a. (**C**) Thermal stability of Clade V1A.1 recombinant viruses. Viruses normalized to 64 hemagglutination units (HAU) were incubated at 33°C-65°C for 20 minutes, and haemagglutination activity was measured using 1% guinea pig red blood cells and expressed as log_2_HAU. (**D**) Thermal stability of Clade V1A.3 recombinant viruses, evaluated using the same conditions described in panel c. Statistical significance was determined using linear mixed-effects models and one-way ANOVA with Tukey’s post hoc test. Data represent the mean ± SEM (pH assay: n=2; temperature assay: n=3).

Overall, Clade V1A.1 mutants were more sensitive to acidic pH and heat compared with rgCOL/17. Several mutants – including Mutant 3 (2aa re-insertion), Mutant 5 (180I), Mutant 6 (498R), Mutant 11 (2aa re-insertion/180I), Mutant 13 (129D/180I/498R) and Mutant 15 (2aa re-insertion/129D/498R) – exhibited markedly reduced infectivity at pH 6.5 and loss of HA activity above 40°C (Fig. 7A, C). Notably, at the more acidic pH of 5.0, rgCOL/17 showed a significant 10-fold reduction in infectivity relative to pH 7.0 (longer horizontal bar in Fig. 7A; P<0.0001, table S12), whereas Mutants 3, 5 and 7 retained higher acid tolerance (shorter horizontal bars, Fig. 7A).

Compared with rgCOL/17 (2aaΔ), re-insertion of the 2-aa alone (Mutant 3) significantly reduced viral titers at pH below pH 6.5 (indicated by vertical bars in Fig. S7A; P=0.0007, table S11) and markedly decreased HA activity at 40°C (indicated by vertical bars in Fig. S8E; P=0.0001, table S14). Reversion of K498R (Mutant 6) preserved acid tolerance similar to rgCOL/17, but HA activity was reduced twofold between 35°C and 45°C (Fig. 7, fig. S7B, F).

Mutant 13 (129D/180I/498R) exhibited pronounced instability, with ∼10-fold lower infectivity at pH 5.0 (Fig. 7A; P<0.0001, table S8) and a ∼2–3-fold reduction in HA activity between 35°C and 55°C (Fig. 7C; P=0.0001, tables S13 and S17), compared with Mutant 7 (Fig. 7, fig. S7C, G). Notably, Mutant 7 with 129D/180I substitutions but retaining 498K maintained high infectivity across varying acidity even at pH 5.0 and thermal stability up to 45°C, similar to rgCOL/17 (Fig. 7, fig. S7C, G). However, Mutant 13 with ancestral substitutions (129D/180I/498R) are pH and temperature sensitive, compared to rgCOL/17 (fig. S7D, H). As such, our data suggest that 2aa deletion or 498K enhance overall HA stability in Clade V1A.1 viruses.

In contrast, all Clade V1A.3 mutant viruses showed pH tolerance and thermal stability comparable to rgWA/19 (3aaΔ; Fig. 7C, D). Relative to rgCOL/17 (2aaΔ), 3aa mutant viruses displayed higher pH tolerance but were slightly more temperature-sensitive (Fig. 7B, D).

Notably, re-insertion of 3aa (Mutant 20) resulted in a reduction in HA activity at 35°C, whereas other Clade V1A.3 mutants only showed reduced activity at 40–45°C. Heat duration assays at 50°C confirmed similar decline in HA activity among Clade V1A.3 mutants and rgWA/19 (fig. S8E, F). As such, Clade V1A.3 viruses (3aaΔ) with enhanced pH tolerance and thermal stability may persist longer in humans and environment, increasing transmission likelihood.

## DISCUSSION

Rapid gene mutations within B/Victoria have led to the emergence of new clades, sparking waves of epidemics globally. Through integrated phylodynamic and functional analyses, we show that the displacement of two-amino acid deletion (2aaΔ) strains by 3-amino acid deletion (3aaΔ) HA variants drove major lineage turnover of B/Victoria viruses, resulting in strong selective sweeps and successive waves of influenza B epidemics. These deletions at positions 162–164 lie within the HA 160-loop, one of the four key antigenic sites (120-loop, 150-loop, and 190-helix) (*14*). In this study, we investigated the virological consequences of HA deletions and associated substitutions among two prominent B/Victoria clades. Compared with the ancestral B/Victoria virus lacking deletions, both Clade V1A.1 (2aaΔ) and V1A.3 (3aaΔ) viruses demonstrated enhanced replication and fitness, replicating efficiently both *in vivo* and *in vitro*. In Clade V1A.1, the 2aa deletion synergized with D129G, I180V, and R498K substitutions were shown to enhance replication efficiency, increase lung viral loads and exacerbate disease severity in mice. Reintroduction of residues 162–163 attenuated virulence, while individual reversions revealed distinct functional roles: D129G was essential for virus viability, whereas V180I and K498R attenuate replication and virulence. Dual substitutions, specifically G129D/V180I, Ins2aa+129D and Ins2aa+498R, restored virulence and altered antigenicity, highlighting the complex epistatic interactions that stabilize HA and modulate disease severity.

Both Clade V1A.1 and V1A.3 displayed robust replication with dual α2,6- and α2,3-linked sialic acid binding, contrasting with B/Yamagata-lineage viruses that preferentially bind to α2,6 receptors (*7, 15*). However, V1A.3 viruses, the 3-amino-acid deletion together with G133R/K136E substitutions exhibited greater virulence and pathogenesis in mice, causing severe lung inflammation and 100% mortality. Reverting these deletions or substitutions attenuated replication and virulence, underscoring their critical role in fitness. Stability assays further showed that rgCOL/17 (2aaΔ) and rgWA/19 (3aaΔ) HAs were resilient to acidic pH and elevated temperatures (up to 45°C), whereas ancestral reversions exhibited increased susceptibility to inactivation. HA stability is a key determinant of viral pathogenicity, adaptability, host range, and transmissibility, and is typically assessed by infectivity or hemagglutination activity (*16, 17*) after exposure to defined pH and temperature conditions (*18, 19*). Both viruses replicated efficiently at 33°C in MDCK cells, consistent with previous reports for the B/Washington/2019 strain (*20, 21*). Notably, rgWA/19 and its variants retained infectivity following low-pH exposure. In comparison with influenza A viruses, HA fusion pH influences host adaptation, with human seasonal viruses typically fusing at lower pH (∼5.0–5.5), swine viruses at intermediate pH (∼5.5–5.9), and avian viruses require higher pH thresholds (∼5.6–6.0) (*22, 23*). Current Clade V1A.3 viruses have undergone rapid genetic and antigenic diversification and remained a persistent public health burden. Post-vaccination sera from individuals exposed to Clade V1A.1 viruses show poor reactive to Clade V1A.3 viruses, and younger children (0–18 years) have been reported to have lower levels of B/Victoria protection to adults due to a lack of preexisting immunity (*24–26*).

Amino acid deletions in the HA has been reported to occur during the very early evolution of IBV strains, exclusively in very early IBV viruses from 1940 to 1993, and in early B/Yamagata lineage viruses (*27, 28*). These deletions typically involved one to three amino acid deletions at positions 163–165 (*28*). As recent study indicated these amino acid deletions had minimal impact on antigenicity (*29*). Since the 2000s, however, B/Victoria lineage exhibits higher genetic diversity, more pronounced antigenic drift and greater seasonal fluctuations than B/Yamagata lineage (*6–8*). Consistent with this, we observed antigenic differences in Clade V1A.1 and V1A.3 viruses compared with B/Brisbane/60/2008, aligning with prior study (*30*). Current influenza B vaccine strategies include a single B/Victoria candidate, but given the rapid evolution and diversification of deletion variants, we speculate that clade-specific vaccines or broadly neutralizing vaccines may improve protection and reduce risk of vaccine mismatch against co-circulating strains. Comprehensive studies of adaptive T-cell responses will be critical to assess protective capacity following immunization. Improved predictive tools are essential for anticipating lineage dominance, antigenic change, and potential vaccine impact.

We note several limitations in this study: (1) the use of mice, while susceptible to influenza B and easier to handle than ferrets or hamsters, may not fully recapitulate human disease; (2) our PET imaging was limited to a subset of strains due to cost, although this non-invasive approach allowed longitudinal assessment of pulmonary inflammation in live animals.

In summary, our study integrates molecular, phenotypic, and phylodynamics analyses to illustrate the emergence and evolution of B/Victoria-lineage deletion variants dominating global circulation. We show that a few nucleotide deletions in the HA head domain – through subtle in structure – profoundly affect viral fitness, replication and pathogenesis. The sequential dominance of 2aa and 3aa deletion variants reflects ongoing antigenic drift coupled with specific genetic mutations under immune pressure and/or vaccinations. Our work highlights the importance of continued genomic surveillance and functional characterization of emerging virus variants to guide vaccine strain selection. Together, this study reveal key drivers reshaping contemporary evolution and transmission of influenza B viruses, providing a mechanistic explanation for its rapid resurgence and dominance, and highlighting challenges for effective influenza control strategies.

## MATERIALS AND METHODS

### Ethics approval

All animal experiments were performed in an animal biosafety level 2+ facility at Duke-NUS Medical School, Singapore with the approval of the NUS Institutional Animal Care and Use Committee (NUS-IACUC 2020/SHS/1606).

### Phylogenetic analysis

All available influenza B sequences (n=33,779) were downloaded from GISAID and the National Center for Biotechnology Information (NCBI) Virus databases from January 2007 to December 2024 (assessed as of 19 February 2025). Multiple sequence alignment of hemagglutinin (HA) gene sequences were performed using MAFFT v7.522 (*31*), and a large preliminary phylogenetic tree was reconstructed using FastTree v2.2 (*32*). The dataset was then downsampled to 768 sequences using SMOT method (*33*) and maximum likelihood tree was reconstructed using RAxML v8.2.12 (*34*). Temporal signal was assessed and outlier sequences were removed using TempEST (*35*). Time-scaled phylogenetic trees were reconstructed using BEAST X (*36*) under an HKY+G substitution model with an uncorrelated relaxed molecular clock. The Gaussian Markov Random Field (GMRF) Bayesian Skyride model (*37*) was applied to estimate changes in effective population size (i.e., relative genetic diversity) over time. Four independent Markov chains of 100 million states, sampled every 10,000 states, were run and combined using LogCombiner after removing the first 10% as burn-in. All parameters of estimated sample sizes (ESS) exceeded 200, indicating adequate mixing and convergence of runs. Maximum clade credibility (MCC) tree was generated using TreeAnnotator. Amino acid substitution analysis was performed using Treesub (*38*) using BASEML from the PAML package (*39*). Phylogenetic tree was illustrated using R package ggtree (*40*) and ggplot2 (*41*). Relative genetic diversity of B/Victoria lineage based on HA gene was plotted using ggplot2.

The distribution and prevalence of no deletion, 2 amino acids deletion and 3 amino acids deletion was plotted with ggplot2. Notably, the clade designated as V1A.3 in this study corresponds to the clade previously referred to as V1A.2 in our earlier work, reflecting updated Nextstrain nomenclature. The hemagglutinin protein structures of B/Brisbane/60/2008 (Clade V1A, no deletion), B/Colorado/06/2017 (Clade V1A.1, 2-aa deletion) and B/Washington/02/2019 (Clade V1A.3, 3-aa deletion) were predicted using Alphafold3 (*42*). Pairwise root mean square deviation (RMSD) values (in Ångströms, Å) between the predicted structures were calculated from the output files. The final structures were visualized and annotated using PyMOL (*43*).

### Cells

Madin-Darby canine kidney (MDCK) and human embryonic kidney x3T (HEK-293T) cells were maintained in high glucose, GlutaMAX supplement Dulbecco’s Modified Eagle Medium (DMEM, Gibco, New York, USA) supplemented with 10% FBS (S181H Heat inactivated, Biowest), 1% penicillin-streptomycin (P/S, Cytiva) and buffered with 25mM HEPES (Gibco, New York, USA). The cells were cultured in 37°C in 5% CO_2_.

### Generation of recombinant B/Victoria viruses by reverse genetics

All recombinant viruses, including wild-type B/Brisbane/60/2008, B/Colorado/06/2017 HA and B/Washington/02/2019 HA were generated by using reverse genetics methods. All experiments were performed in Biosafety Level 2 plus (BSL2+) laboratory approved by National University of Singapore. Briefly, the HA sequences were cloned into the pAD3000 bidirectional vector (*44*) by Genscript (Biotech Corporation, China). The B/Brisbane/60/2008 (BRIS/08) wild-type strain was used as B/Victoria backbone, comprising eight plasmids, each carrying one gene segment flanked by both RNA polymerase I (pol I) and II (pol II) promoters. For the BRIS/08 wild-type recombinant virus, all eight gene segments (PB2, PB1, PA, HA, NP, NA, M and NS) were derived from B/Brisbane/60/2008. For all other recombinant viruses, we used an eight-gene plasmid (7+1) system, containing seven segments (PB2, PB1, PA, NP, NA, M and NS) from B/Brisbane/60/2008, whereas the HA segment was either B/Colorado/06/2017 for Clade V1A (2aaΔ) or B/Washington/02/2019 for Clade V1A.3 (3aaΔ) viruses. Mutant viruses were generated by introducing specific HA amino-acid substitutions onto the HA gene. All recombinant viruses were generated by cotransfecting MDCK and 293T cells with eight different plasmids using Fugene 6 transfection agent (Promega, Wisconsin, USA) according to the manufacturer’s instructions. All rescued viruses were passaged twice onto MDCK cells virus supernatants were harvested followed by RNA extractions using Direct-zol^TM^ RNA Miniprep (Zymo Research Corporation, Irvine, USA). The sequences of all recombinant viruses were confirmed by next-generation sequencing (NGS) using previously described methods (*8*). The corresponding virus titers were assessed using TCID_50_ assay on MDCK cells.

### 50% Tissue culture infectious dose (TCID_50_) assay

MDCK cells were seeded in DMEM supplemented with 10% FBS, 1% P/S and 25mM HEPES in a 96-well flat-bottom plate. Virus samples were prepared by 10-fold serial dilution in DMEM containing 1% P/S, 1μg/ml TPCK-treated trypsin (Sigma-Aldrich, Missouri, USA) and 3% bovine serum albumin (BSA, ThermoFisher Scientific, Massachusetts, USA). After 1 hour adsorption at 33°C in 5% CO_2_, 100μl of the virus medium was added to each well, and plates were incubated for 72 hours.

Cells were fixed with 80% cold acetone for 10 minutes, washed with 1X PBST (Tween 20, Sigma-Aldrich, Missouri, USA) and stained with anti-influenza B nucleoprotein mouse monoclonal antibody (Abcam, UK) followed by HRP conjugated goat-anti-mouse secondary antibody (Abcam, UK). The *O*-phenylenediamine dihydrochloride (OPD) substrate (Sigma-Aldrich, Missouri, USA) dissolved in phosphate-citrate buffer with sodium perborate (Sigma-Aldrich, Missouri, USA) was added in the plates and the reaction was stopped with 0.5M sulfuric acid (ThermoFisher Scientific, Massachusetts, USA). Absorbance at 490nm was measured using Tecan Spark 10M microplate plate reader (Tecan, Mannedorf, Switzerland). The TCID_50_ values were calculated using the Reed and Muench method (*45*).

### Haemagglutination and haemagglutination inhibition assay

Haemagglutination (HA) and haemagglutination inhibition (HI) assays were performed following standard procedures(*46*). Viruses were serially diluted 2-fold and mixed with 1% guinea pig red erythrocytes. After 1 hour of incubation at 4°C, HA titers were determined as the highest dilution that produced complete agglutination of erythrocytes. For HI assays, 8 HA units of virus were incubated with B/Victoria reference goat antisera B/Brisbane/60/2008 (International Reagent Resource) at room temperature for 30 minutes and inhibition of haemagglutination was assessed.

### Mice experiments

Three- to four-week-old female BALB/c mice (InVivos, Singapore) were initially anesthetized by intraperitoneal injection of 70 mg/kg ketamine and 10 mg/kg xylazine and then intranasally infected with 1×10^5^ TCID_50_ (50μl) of virus. Survival and weight loss were monitored daily for 14 days, and animals that lost 25% or more of their initial body weight were euthanized. For each recombinant virus, group of 5 mice were used. Negative control mice were intranasally inoculated with 50μl of phosphate-buffered saline (PBS, Lonza, Switzerland). To determine the virus lung titers, mice were euthanized and lungs were harvested on days 3 (n=4 mice/group) and 5 (n=4 mice/group) after infection. Lung virus titers were determined by TCID_50_ assay in MDCK cells.

### Non-invasive PET/CT imaging of infected mice

Positron emission tomography/computed tomography (PET/CT) imaging was performed longitudinally to compare the extent of lung inflammation between Clade V1A.1 and V1A.3 viruses, represented by rgCOL/17 and rgWA/19, respectively. Groups of mice (n=4) were anesthetized and inoculated with virus as described above, and baseline measurements were obtained from each animal prior to infection (pre-infection). Whole-body PET/CT imaging was conducted on days 3 and 5 post-infection as described previously (*47*). Briefly, mice were intravenously injected with ^18^Fluorodeoxyglucose (^18^F-FDG; 20 MBq in 100 µL) and, following 60 min uptake period, subjected to PET/CT scans under continuous anesthesia with 2% (^v^/_v_) isofluorane. Four mice were simultaneously imaged using VECTor^4^CT (MILabs, Netherlands) multi-bed scanner equipped with a HE-GP-RM 3.6mm collimator. Thoracic PET images were acquired for 20 min(*48*), followed by 2 min CT scan (tube current: 0.24 mA; tube voltage: 50 kV). PET images were reconstructed at 0.8-mm voxel resolution using a SROSEM algorithm (16 subsets, 24 iterations, 1.5-μm Gaussian filter) using the MILabs reconstruction software. PET and CT images were automatically co-registered after image reconstruction, and PET attenuation correction was manually performed using the corresponding CT images.

Quantitative image analysis and image data visualization were carried out using PMOD software (v4.4, PMOD Technologies, Switzerland). Volumes of interest (VOIs) were drawn on CT images scaled to Hounsfield Units (HU) using a semi-automated thresholding approach to segment lung space from surrounding tissues. VOIs corresponding to total lungs were manually delineated around the ribcage and refined using the cold iso-contouring function (HU ≤ -300 HU) in PMOD. The final lung VOIs were visually inspected and manually corrected as needed to exclude the heart, major airways, and vessels to ensure consistent and reproducible definition of the lung space for quantitative analysis. The same VOIs were transposed onto PET images and used to quantify lung ^18^F-FDG uptake reported as % injected dose per volume of lung (%ID/mL). We also quantified metabolic lung volume (MLV) –defined as the volume of lung tissue exhibiting ^18^F-FDG uptake above a basal threshold – calculated as the mean plus two standard deviations of pre-infection lung uptake. MLV calculations and image 3D-rendering were performed in PMOD.

On 5 dpi and immediately following PET/CT scans, lungs were harvested and fixed in 4% paraformaldehyde, and processed for embedding in paraffin blocks. Tissue sections (5 µm thick) were stained with hematoxylin & eosin (H&E) for histopathological examination (n=5 mice per group). Contiguous slides were subjected to immunofluorescence staining by overnight incubation with mouse anti-influenza B nucleoprotein (NP) clone B017 (B35G) (Bio-Rad cat. no. MCA403) and rabbit anti-CD45 (Abcam cat. no. ab10558). Tissue sections were counterstained with DAPI, and autofluorescence was quenched using the with Vector^®^ TrueVIEW^®^ kit (Vector Laboratories, USA). Slides were examined and imaged on Nikon Ni-E inverted fluorescence microscope (Nikon, Tokyo, Japan).

### Replication kinetics in MDCK cells

Wild-type or recombinant viruses were inoculated into MDCK cells at a multiplicity of infection (MOI) of 0.01. After 1h of exposure to inoculum, cells were incubated with virus media. Supernatants were collected at 12, 24, 36, 48, 60, and 72 hours post-infection and were stored at -80°C until subsequent titration by TCID_50_ assays in MDCK cells. The TCID_50_ values were obtained from three independent experiments to counter for the biological replicates.

### Receptor binding assay

Receptor-binding specificity was determined by a solid-phase direct binding assay with few modifications, as previously described (*49*). Serial dilutions (0.01, 0.03, 0.1, 0.3, and 0.9μg/ml) of biotinylated glycans 3’sialyllactosamine (3’SLN-C3-PAA-BIOT, GNZ-0036-BP, Sigma Aldrich, Missouri, USA) and 6’sialyllactosamine (6’SLN-C3-PAA-BIOT, GNZ-0997-BP, Sigma-Aldrich, Missouri, USA), were prepared in PBS. A 100μl aliquot of each dilution was added to 96 well flat-bottom plate and allowed to attach overnight at 4°C. After removing the glycan solution, wells were washed with ice-cold PBS containing 0.1% Tween 20 (PBST) and blocked with 2% BSA (ThermoFisher Scientific, Massachusetts, USA) containing Oseltamivir NA inhibitor (AOBIOUS INC, Massachusetts, USA) for 1 hour at room temperature. Plates were then incubated with 64 HA units of recombinant viruses overnight at 4°C. Following washes with PBST, wells were incubated with HRP-Streptavidin (Sigma-Aldrich, Missouri, USA) for 2 hours at 4°C. After a final wash, 50μl of 3,3’,5,5’-Tetramethylbenzidine (TMB, Sigma-Aldrich, Missouri, USA) was added for 10 minutes at room temperature in the dark. The reaction was stopped with 50μl of 1M sulphuric acid and absorbance was measured at 450nm using a microplate reader (Tecan, Mannedorf, Switzerland).

### Thermostability assay

HA thermostability was determined by measuring the temperature at which the viruses lost their ability to agglutinate red blood cells. The recombinant viruses were diluted to 64 HA units and incubated at an accelerated temperature (i.e. 33°C, 37°C, 40°C, 45°C, 50°C, 55°C, 60°C and 65°C) for 20 minutes or incubated at 50°C for 12-hour duration (i.e. 1, 2, 3, 4, 5, 6 and 12 hours). After incubation, HA assays were performed according to standard methods using 1% guinea pig red blood cells (Innovative Research, Michigan, USA).

### pH stability

The acid stability of the viruses was evaluated by measuring their infectivity after acid treatment. The recombinant viruses were diluted to 10^5^ TCID_50_/ml in PBS at varying pH levels (pH 7.0, 6.5, 6.0, 5.6 and 5.0), adjusted by adding 0.1M hydrochloric acid. After 15 minutes incubation at 33°C, the treated viruses were used to infect MDCK cells and infectivity was determined by TCID_50_ assay.

### Immunofluorescence assay

Transwell inserts were fixed in 10% neutral buffered formalin for 30 minutes, washed with PBS and permeabilized in 0.1% Triton X-100 (Sigma-Aldrich, Missouri, USA) in PBS. Cells were blocked for 60 minutes in 1% BSA containing 0.1% Triton X-100 and incubated overnight at 4°C with primary antibody mouse anti-influenza B nucleoprotein (NP, MCA403, 1:100, BioRad, California, USA). After three PBS washes, cells were incubated for 1 hour at room temperature with an Alexa Fluor 555-conjugated secondary antibody (1:100, ThermoFisher Scientific, Massachusetts, USA) followed by counterstained with Hoechst, DAPI (1:500, ThermoFisher Scientific, Massachusetts, USA) and phalloidin (1;1000, ThermoFisher Scientific, Massachusetts, USA). Samples were mounted in Prolong Antifade (ThermoFisher Scientific, Massachusetts, USA) and imaged on a Andor BC43 confocal microscope using Imaris Viewer Software (Oxford Instruments, UK).

### Statistical analysis

All statistical analyses were performed using GraphPad Prism software version 10 (GraphPad Software, La Jolla, USA) unless otherwise specified. Differences between experimental groups were assessed by one-way analysis of variance (ANOVA) and post hoc Tukey’s test, while animal survival was analysed using the log-rank (Mantel-Cox) test. A P-value of <0.05 was considered statistically significant. Receptor-binding specificity was visualized in R using ggplot2. Mean absorbance across three biological replicates for each mutant was displayed as bar charts with standard deviation error bars. Total receptor binding to 3’-sialyllactosamine (3′ SLN) and 6’-sialyllactosamine (6′ SLN) was quantified by summing absorbance values across all concentrations per replicate and plotted as boxplots. Temperature and pH response data were log₂-transformed and analysed using a linear mixed-effects model fitted using the lmer function from the lme4 package in R. In this model, *virus mutants*, *temperature/pH*, and their interaction were treated as fixed effects, while *replicate* nested within *strain* was included as a random effect to account for variability among repeated measurements. This approach allowed us to assess both how each virus mutant responded across different conditions and whether significant differences existed between strains under the same condition. Model significance was evaluated using ANOVA implemented in the lmerTest package, and post hoc pairwise comparisons were performed with the emmeans package using Tukey’s adjustment for multiple testing. Heatmaps were generated in R using the ComplexHeatmap package. Comparisons between two groups in ^18^F-FDG-PET/CT imaging were performed using Welch’s t-test.

## Acknowledgments

This study was supported by Singapore Ministry of Health’s National Medical Research Council (NMRC) under Clinical Science-Investigator Research Grant CS-IRG/MOH-000374 (G.J.D.S & Y.C.F.S.); Open-Fund Large Collaborative Grant OF-LCG/MOH-000505-05; and by the Duke-NUS Signature Research Programme funded by Ministry of Health, Singapore. We thank GISAID and all submitters for access to their databases and all submitters of data and sequences to these databases.

## Author contributions

G.J.D.S. and Y.C.F.S. conceived and designed research. G.G.K.N. and W.F.Y. performed experiments. G.G.K.N., P.C., X.L., D.Q.C. and R.Z. analysed data. G.G.K.N., P.C., X.L. and Y.C.F.S. designed Figures. C.B.L.V., A.G. and A.M.C. performed PET/CT. R.J.W. provided reagents. G.G.K.N. and Y.C.F.S. co-drafted the manuscript with input from R.J.W., P.A.T. and G.J.D.S. All authors contributed to reviewing and editing of the manuscript.

## Data and materials availability

All data are available in the main text or the supplementary materials.

## Supplementary Materials

Figs. S1 to S8

Tables S1 to S19

