## Supplementary Files for "Haemagglutinin 162–164 deletions enhance influenza B/Victoria virus fitness and virulence in vivo"

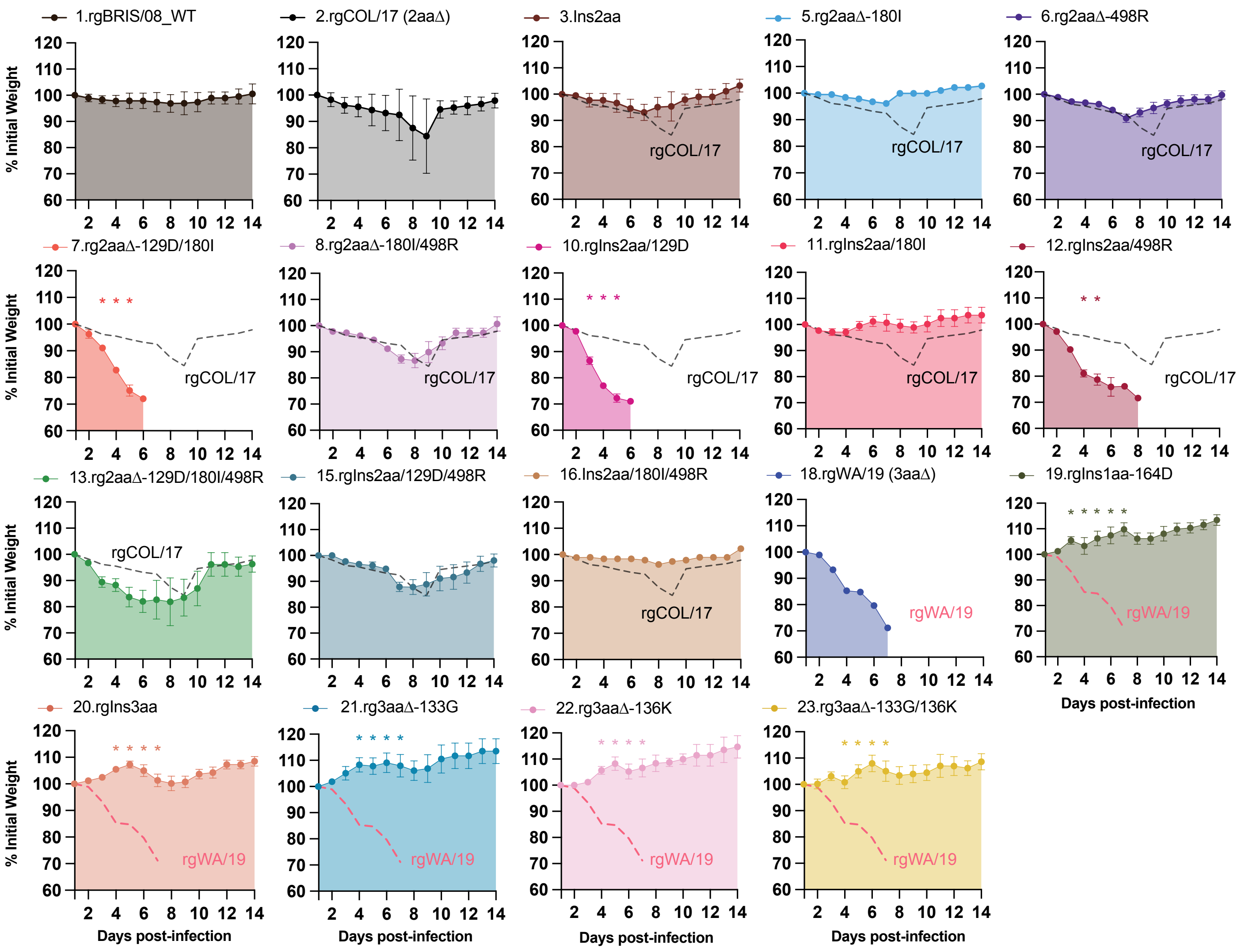


**Fig. S1. The effects of B/Victoria HA amino acid substitutions on changes in body weight in mice.** Body weight trajectories of mice intranasally infected with $10^5$ $TCID_{50}$ of recombinant B/Victoria viruses. Each panel represents a individual recombinant virus, and percent baseline body weight (mean ± SEM) is shown over a 14-day monitoring period. Black and pink dotted lines denote rgCOL/17 and rgWA/19 respectively. Each group consisted of n=5 mice. Statistical significance was assessed using one-way ANOVA followed by Tukey test.*P< 0.05 indicates significance.

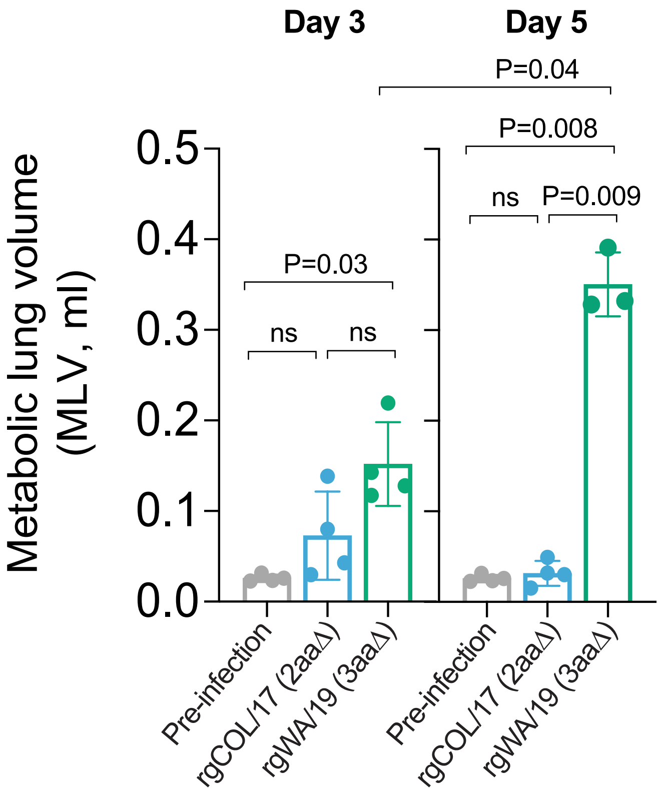


**Fig. S2. Metabolic lung volume in infected mice.** Metabolic lung volume (MLV) was quantified from $^{18}$F-FDG-PET/CT images at days 3 and 5 post-infection in rgCOL/17 (2aa Δ), and rgWA/19 (3aaΔ) mice. Grey bars indicate basal level of the same mice prior to infection. Data represent mean ± SEM and were compared using the Mann-Whitney test. ns, not significant. Each group consisted of *n*=4 mice.

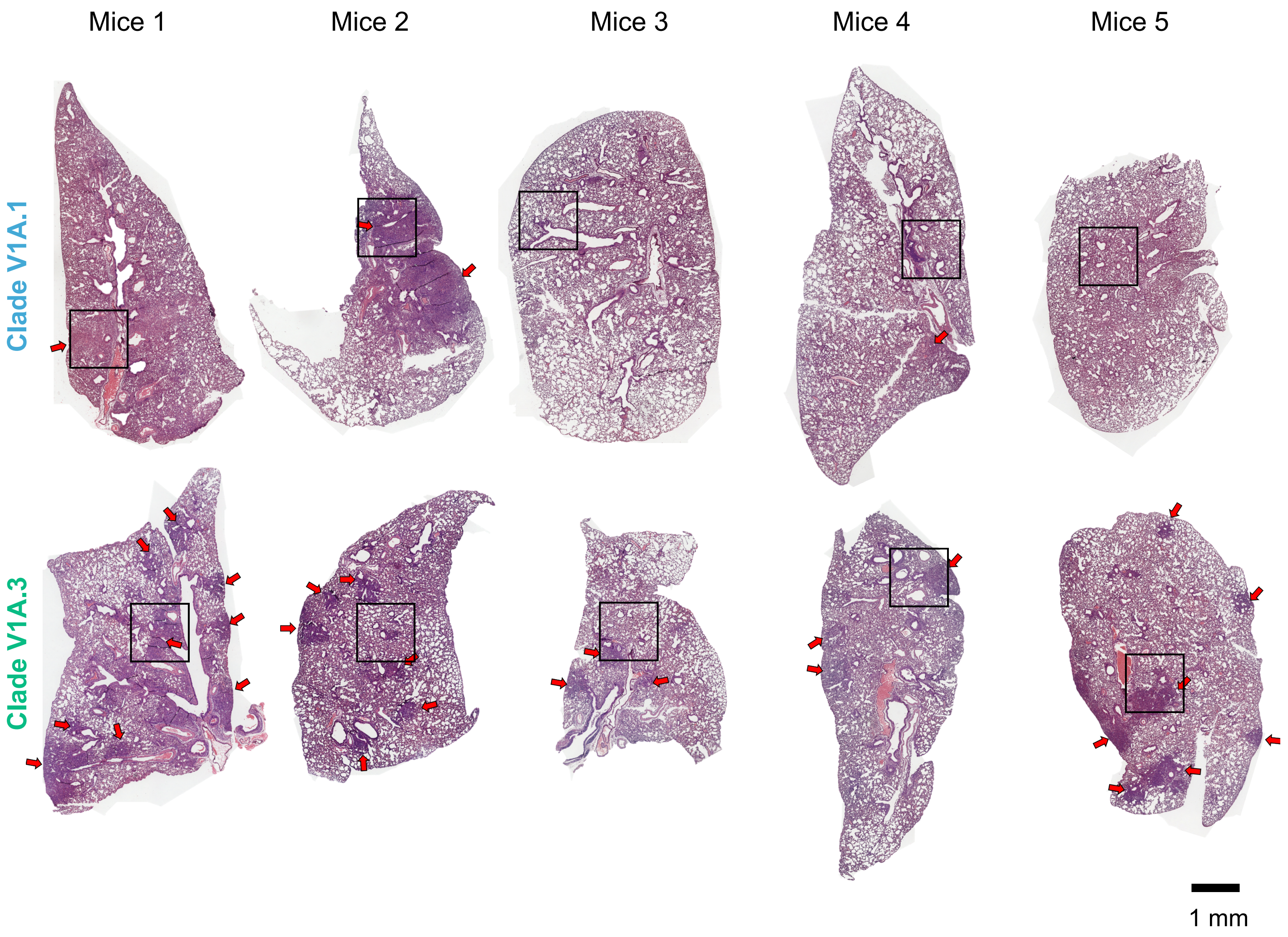


**Fig. S3. Histopathological examination of mouse lungs following infection with Clade V1A.1 (rgCOL/17) and Clade V1A.3 (rgWA/19) viruses.** Hematoxylin and eosin (H&E)-stained sections of the left lung lobe collected on day 5 post-infection from mice infected with either rgCOL/17 or rgWA/19 virus. Broad regions of inflammation are marked with red arrows and outlined boxed indicate areas selected for higher magnification (as shown in Fig. S4). Scale bar: 1mm. Each group included *n*=5 mice.

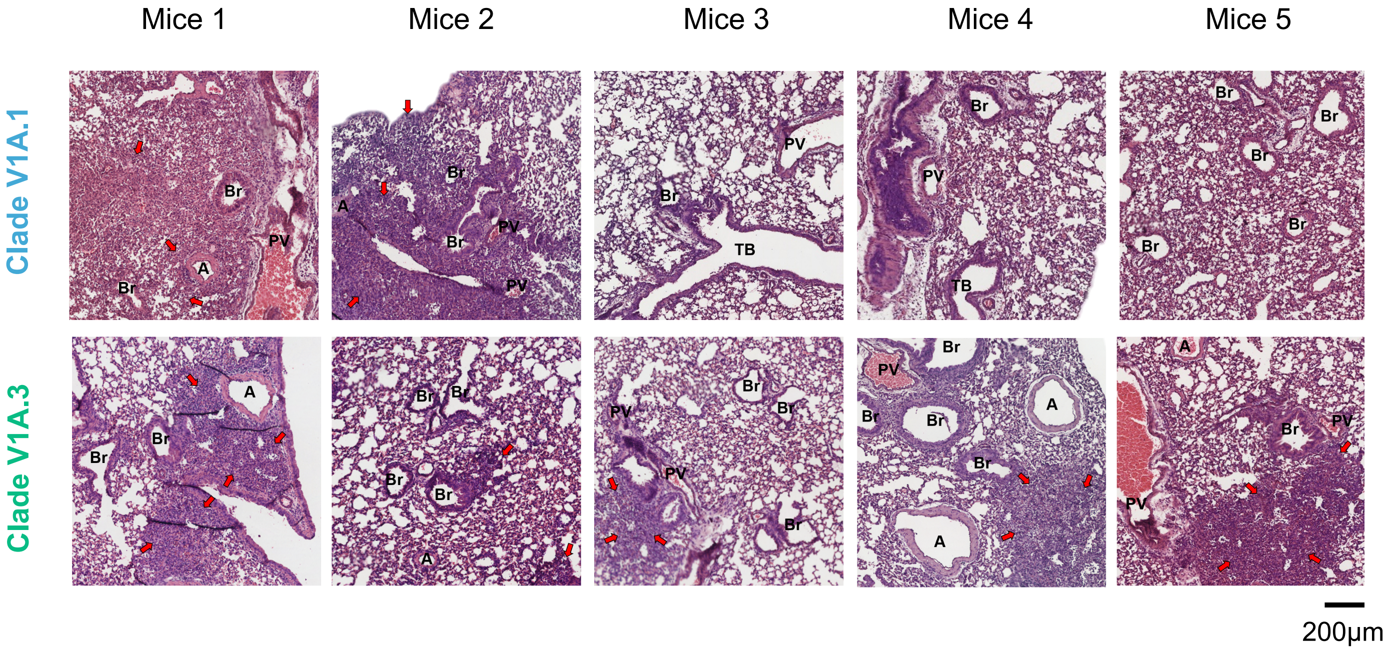


**Fig. S4. Assessment of lung pathology in mice at higher magnification.** Hematoxylin and eosin (H&E)-stained images highlight structural features within affected regions, including artery (A), bronchiole (Br), pulmonary vein (PV), and terminal bronchi (TB). Red arrows indicate rominent inflammation. Scale bar: 200μm. Each group consisted of of *n*=5 mice.

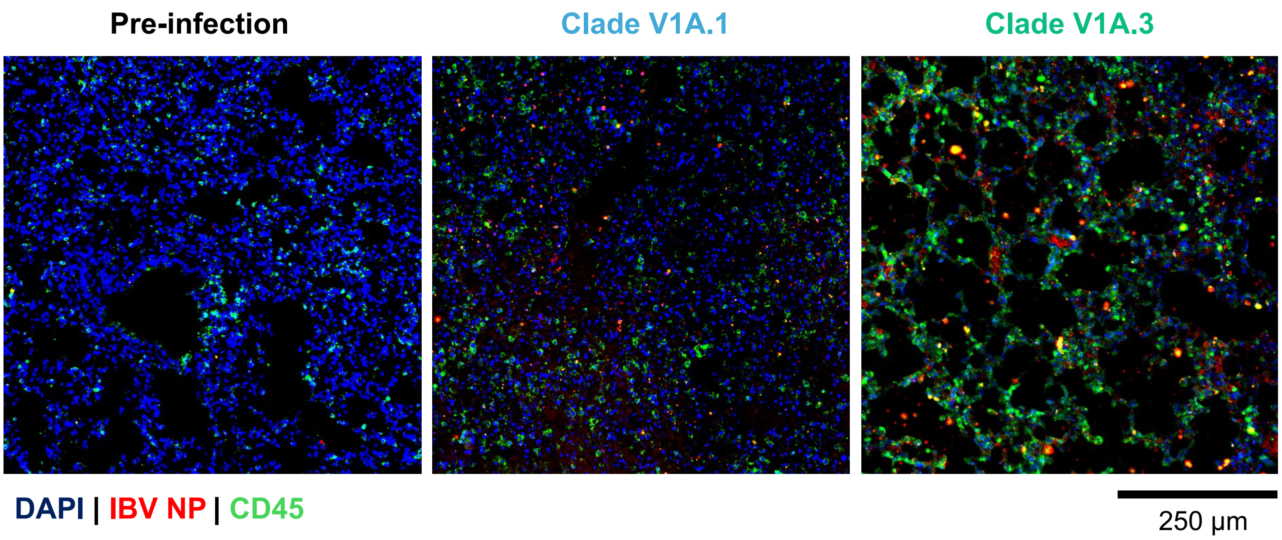


**Fig. S5. Immunohistopathology images of lungs from mice infected with Clade V1A.1 and V1A.3 viruses.** Immunofluorescence staining of mouse lungs infected with Clade 1A.1 (rgCOL/17) and Clade 1A.3 (rgWA/19) viruses at day 5 post-infection. Viral nucleoprotein (NP) is shown in red, immune cell marker CD45 is shown in green and nuclei in blue. Scale bar: 250µm.

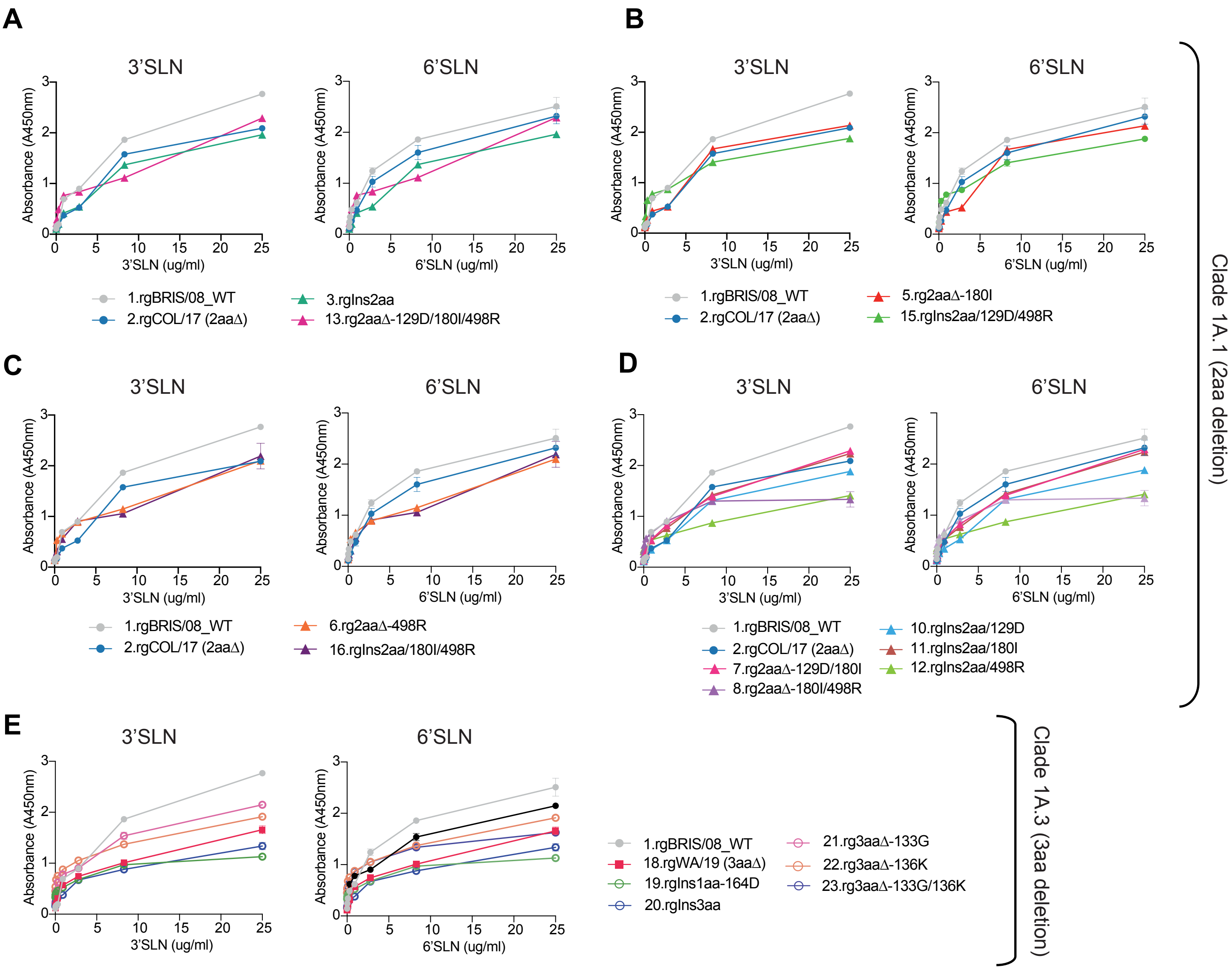


**Fig. S6. Receptor binding profiles of recombinant B/Victoria viruses.** Receptor binding affinities to α2,3-linked (3'SLN-PAA) and α2,6-linked (6'SLN-PAA) sialylglycopolymers were measured using solid-phase direct binding assays. Serial dilutions of biotinylated glycans were incubated with 64 HAU of each virus, and bound glycans were detected using HRP-conjugated streptavidin and TMB subtrate, with absorbance recorded at 450 nm. (**A-D**) Panels correspond to Clade V1A.1 (2aaΔ) viruses and (**E**) panel corresponds to Clade V1A.3 (3aaΔ) viruses. Statistical significance was evaluated by one-way ANOVA. Data represent mean ± SEM from three independent experiments.

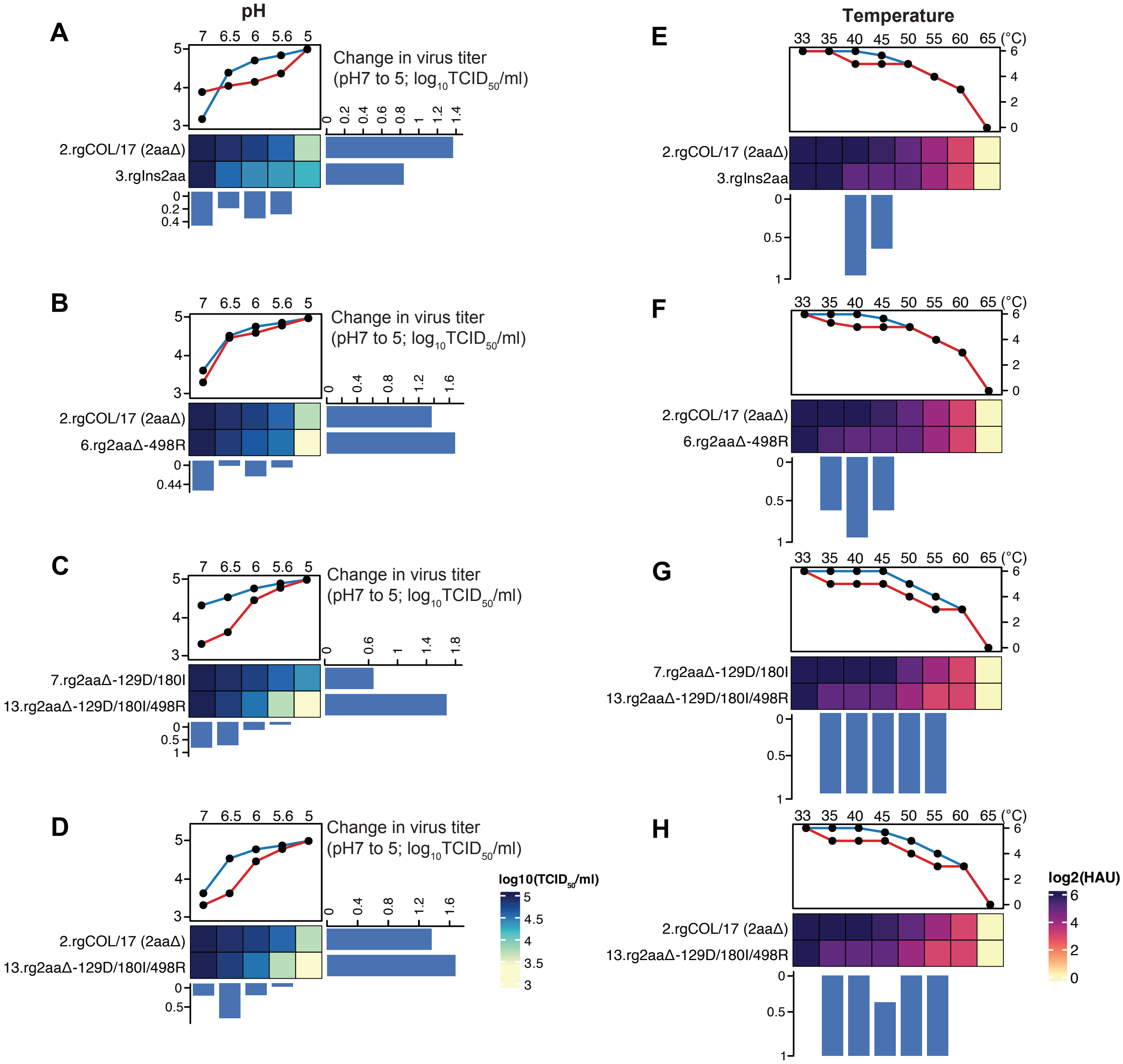


**Fig. S7. Acid and thermal stability profiles of Clade V1A.1 recombinant viruses.** (**A-D**) Recombinant viruses from Clades V1A.1 were incubated in buffers with pH values ranging from 5.0 to 7.0, followed by infection of MDCK cells. At 72 hours post-infection (hpi), cells were fixed and stained for viral NS1 protein, and viral titers were quantified by $TCID_{50}$ assay. (**E-H**) For thermal stability analysis, viruses were standardized to 64 hemagglutination units (HAU) and incubated at temperatures between 33°C and 65°C for 20 minutes. Hemagglutination activity was assessed using 1% guinea pig red blood cells, and titers were expressed as $log_2$HAU. Statistical significance was determined using linear mixed-effects model and one-way ANOVA with Tukey's post hoc test. Data represent the mean ± SEM (pH assay: *n*=2; temperature assay: *n*=3).

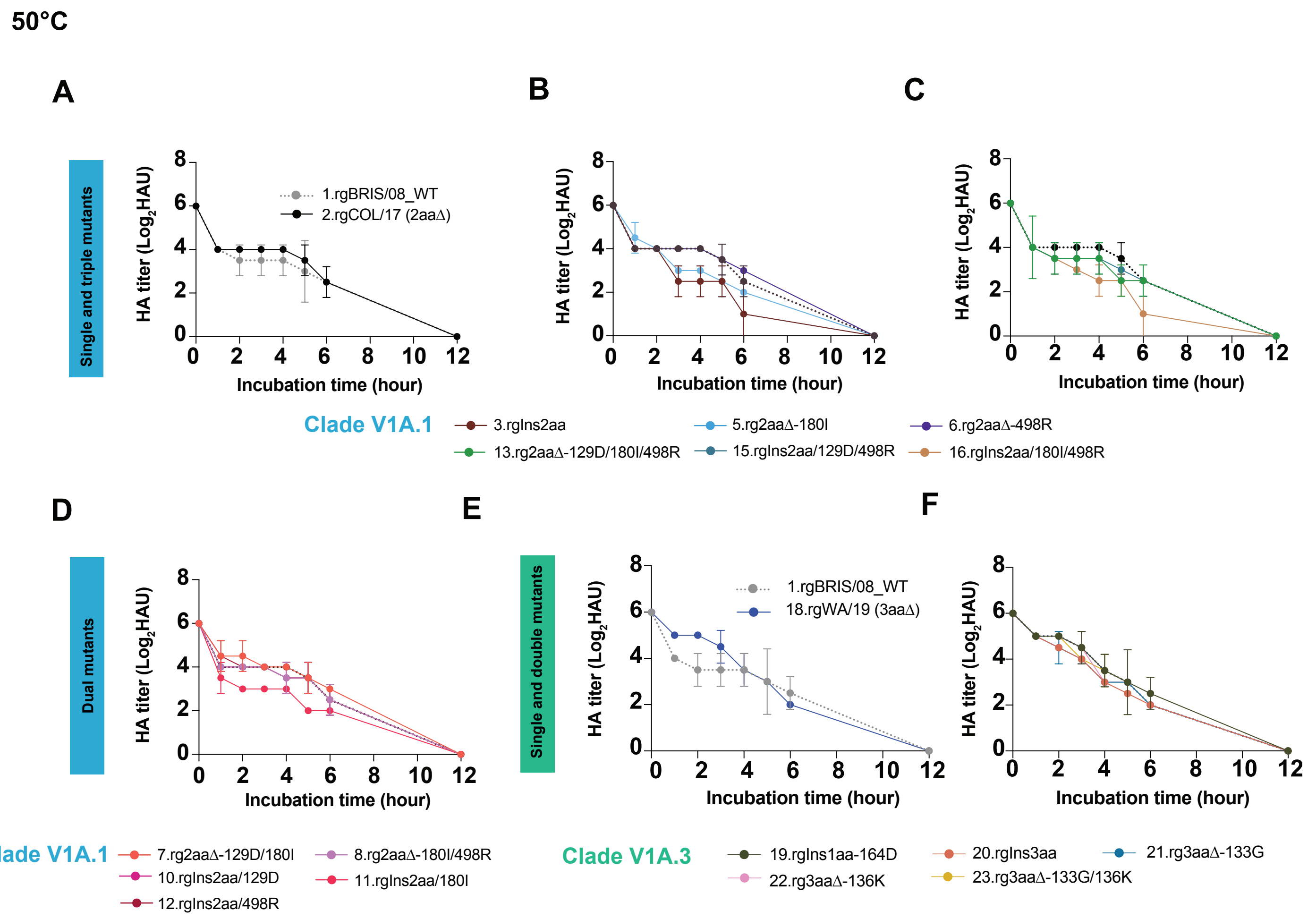


**Fig. S8. Time-course thermal stability profiles of Clades V1A.1 and V1A.3 recombinant viruses.** (**A-D**) Recombinant viruses from Clade V1A.1 and (**E-F**) Clade V1A.3 were incubated at 50°C and sampled at multiple time points over a 12-hour period. Hemagglutination activity was measured using 1% guinea pig red blood cells, and titers were expressed as $log_2$HAU. Statistical significance was assessed by one-way ANOVA. Data represent mean ± SEM from three biologically independent experiments (*n*=3).

**Table S1. TMRCA estimates for HA gene segment in Victoria clade V1A.1 and V1A.3.**

| Clade | Mean TMRCA | Lower 95% HPD | Upper 95% HPD | PP |
|---|---|---|---|---|
| V1A.1 (2aaΔ) | 2016.62 (13 Aug 2016) | 2016.33 (30 Apr 2016) | 2016.92 (02 Dec 2016) | 1 |
| V1A.3 (3aaΔ) | 2017.50 (01 Jul 2017) | 2023.79 (15 Oct 2023) | 2024.77 (08 Oct 2024) | 1 |

**PP, posterior probability**

**Table S2. Average lung $^{18}$F-FDG uptake during disease progression.**

| Day | rgCOL/17 | rgWA/19 |
|---|---|---|
| 0 | 5.30 ± 0.39 | 5.30 ± 0.39 |
| 3.5 | 5.09 ± 0.87 | 7.19 ± 1.17 |
| 5.5 | 4.06 ± 0.70 | 10.23 ± 1.17 |

**Table S3. Receptor binding profile for the 3'SLN glycan in three replicates.**

| | | 3'SLN glycan concentration (μg/mL) | | | | | | | |
|---|---|---|---|---|---|---|---|---|---|
| No | Recombinant virus | 25ug/ml | 8.3ug/ml | 2.8ug/ml | 0.9ug/ml | 0.3ug/ml | 0.1ug/ml | 0.03ug/ml | 0.01ug/ml |
| 1 | rgBRIS/08_WT | 2.683 | 1.868 | 0.906 | 0.694 | 0.199 | 0.160 | 0.152 | 0.111 |
| | | 2.885 | 1.805 | 0.900 | 0.696 | 0.206 | 0.179 | 0.146 | 0.113 |
| | | 2.735 | 1.915 | 0.884 | 0.681 | 0.183 | 0.173 | 0.141 | 0.111 |
| 2 | rgCOL/17 (2aaΔ) | 2.008 | 1.563 | 0.527 | 0.390 | 0.172 | 0.136 | 0.139 | 0.108 |
| | | 2.063 | 1.501 | 0.496 | 0.306 | 0.170 | 0.154 | 0.143 | 0.111 |
| | | 2.198 | 1.666 | 0.550 | 0.418 | 0.176 | 0.145 | 0.133 | 0.116 |
| 3 | rgIns2aa | 2.035 | 1.459 | 0.532 | 0.428 | 0.202 | 0.175 | 0.157 | 0.103 |
| | | 1.969 | 1.389 | 0.525 | 0.386 | 0.206 | 0.185 | 0.152 | 0.113 |
| | | 1.883 | 1.255 | 0.575 | 0.442 | 0.209 | 0.175 | 0.157 | 0.100 |
| 5 | rg2aaΔ-180I | 2.135 | 1.742 | 0.544 | 0.417 | 0.281 | 0.161 | 0.158 | 0.112 |
| | | 2.158 | 1.689 | 0.537 | 0.437 | 0.232 | 0.160 | 0.145 | 0.124 |
| | | 2.117 | 1.581 | 0.493 | 0.463 | 0.284 | 0.183 | 0.139 | 0.113 |
| 6 | rg2aaΔ-498R | 2.158 | 1.987 | 0.814 | 0.741 | 0.568 | 0.181 | 0.174 | 0.136 |
| | | 2.059 | 1.844 | 0.928 | 0.690 | 0.525 | 0.202 | 0.164 | 0.145 |
| | | 2.128 | 1.876 | 0.840 | 0.659 | 0.515 | 0.181 | 0.173 | 0.136 |
| 7 | rg2aaΔ-129D/180I | 2.224 | 1.336 | 0.858 | 0.492 | 0.296 | 0.226 | 0.177 | 0.120 |
| | | 2.328 | 1.466 | 0.759 | 0.534 | 0.371 | 0.217 | 0.188 | 0.126 |
| | | 2.322 | 1.368 | 0.867 | 0.591 | 0.285 | 0.208 | 0.188 | 0.140 |
| 8 | rg2aaΔ-180I/498R | 1.518 | 1.351 | 0.937 | 0.657 | 0.598 | 0.467 | 0.350 | 0.226 |
| | | 1.451 | 1.319 | 0.915 | 0.676 | 0.594 | 0.451 | 0.306 | 0.260 |
| | | 1.035 | 1.236 | 0.861 | 0.688 | 0.505 | 0.406 | 0.345 | 0.295 |
| 10 | rgIns2aa/129D | 1.437 | 0.908 | 0.621 | 0.522 | 0.399 | 0.380 | 0.347 | 0.342 |
| | | 1.418 | 0.839 | 0.673 | 0.544 | 0.354 | 0.317 | 0.369 | 0.254 |
| | | 1.374 | 0.864 | 0.587 | 0.541 | 0.338 | 0.356 | 0.369 | 0.357 |
| 11 | rgIns2aa/180I | 2.363 | 1.402 | 0.741 | 0.481 | 0.258 | 0.172 | 0.160 | 0.133 |
| | | 2.137 | 1.392 | 0.761 | 0.556 | 0.285 | 0.182 | 0.163 | 0.139 |
| | | 2.204 | 1.470 | 0.808 | 0.545 | 0.262 | 0.175 | 0.171 | 0.129 |
| 12 | rgIns2aa/498R | 1.437 | 0.908 | 0.621 | 0.522 | 0.399 | 0.380 | 0.347 | 0.342 |
| | | 1.418 | 0.839 | 0.673 | 0.544 | 0.354 | 0.317 | 0.369 | 0.254 |
| | | 1.374 | 0.864 | 0.587 | 0.541 | 0.338 | 0.356 | 0.369 | 0.357 |
| 13 | rg2aaΔ-129D/180I/498R | 2.228 | 1.107 | 0.845 | 0.759 | 0.497 | 0.288 | 0.215 | 0.137 |
| | | 2.275 | 1.133 | 0.851 | 0.760 | 0.499 | 0.290 | 0.237 | 0.132 |
| | | 2.366 | 1.097 | 0.819 | 0.774 | 0.475 | 0.299 | 0.214 | 0.142 |
| 15 | rgIns2aa/129D/498R | 1.914 | 1.317 | 0.920 | 0.790 | 0.656 | 0.329 | 0.241 | 0.138 |
| | | 1.854 | 1.359 | 0.835 | 0.759 | 0.698 | 0.327 | 0.241 | 0.152 |
| | | 1.869 | 1.544 | 0.861 | 0.791 | 0.611 | 0.345 | 0.214 | 0.112 |
| 16 | rgIns2aa/180I/498R | 1.823 | 1.078 | 0.936 | 0.529 | 0.314 | 0.222 | 0.212 | 0.112 |
| | | 2.677 | 1.076 | 0.947 | 0.530 | 0.303 | 0.242 | 0.214 | 0.178 |
| | | 2.071 | 1.013 | 0.841 | 0.611 | 0.317 | 0.237 | 0.198 | 0.133 |
| 18 | rgWa/19 (3aaΔ) | 1.768 | 1.024 | 0.718 | 0.541 | 0.301 | 0.210 | 0.174 | 0.122 |
| | | 1.511 | 1.046 | 0.784 | 0.571 | 0.312 | 0.251 | 0.178 | 0.113 |
| | | 1.690 | 0.959 | 0.732 | 0.593 | 0.321 | 0.203 | 0.157 | 0.110 |
| 19 | rgIns1aa-164D | 1.203 | 0.991 | 0.669 | 0.520 | 0.477 | 0.422 | 0.379 | 0.326 |
| | | 1.153 | 0.956 | 0.728 | 0.523 | 0.466 | 0.415 | 0.360 | 0.309 |
| | | 1.031 | 0.965 | 0.650 | 0.534 | 0.433 | 0.410 | 0.387 | 0.362 |
| 20 | rgIns3aa | 1.314 | 0.890 | 0.678 | 0.404 | 0.319 | 0.270 | 0.187 | 0.150 |
| | | 1.229 | 0.963 | 0.658 | 0.380 | 0.367 | 0.272 | 0.184 | 0.161 |
| | | 1.469 | 0.783 | 0.676 | 0.356 | 0.301 | 0.252 | 0.189 | 0.158 |
| 21 | rg3aaΔ-133G | 2.146 | 1.402 | 0.876 | 0.770 | 0.594 | 0.457 | 0.357 | 0.280 |
| | | 2.159 | 1.633 | 0.943 | 0.767 | 0.638 | 0.470 | 0.387 | 0.214 |
| | | 2.134 | 1.579 | 0.890 | 0.799 | 0.633 | 0.430 | 0.368 | 0.209 |
| 22 | rg3aaΔ-136K | 1.910 | 1.313 | 1.051 | 0.852 | 0.752 | 0.699 | 0.523 | 0.496 |
| | | 1.919 | 1.347 | 1.089 | 0.897 | 0.765 | 0.698 | 0.582 | 0.404 |
| | | 1.910 | 1.456 | 1.018 | 0.880 | 0.734 | 0.636 | 0.504 | 0.458 |
| 23 | rg3aaΔ-133G/136K | 1.700 | 1.314 | 1.054 | 0.805 | 0.631 | 0.571 | 0.523 | 0.423 |
| | | 1.536 | 1.367 | 1.041 | 0.899 | 0.664 | 0.600 | 0.544 | 0.456 |
| | | 1.647 | 1.328 | 1.060 | 0.888 | 0.614 | 0.544 | 0.522 | 0.471 |

**Table S4. Receptor binding profile for the 6'SLN glycan in three replicates.**

| | | 6'SLN glycan concentration (μg/mL) | | | | | | | |
|---|---|---|---|---|---|---|---|---|---|
| No | Recombinant virus | 25ug/ml | 8.3ug/ml | 2.8ug/ml | 0.9ug/ml | 0.3ug/ml | 0.1ug/ml | 0.03ug/ml | 0.01ug/ml |
| 1 | rgBRIS/08_WT | 2.569 | 1.804 | 1.217 | 0.602 | 0.457 | 0.382 | 0.298 | 0.150 |
| | | 2.180 | 1.859 | 1.142 | 0.669 | 0.495 | 0.280 | 0.128 | 0.138 |
| | | 2.779 | 1.915 | 1.366 | 0.588 | 0.503 | 0.359 | 0.295 | 0.149 |
| 2 | rgCOL/17 (2aaΔ) | 2.403 | 1.878 | 1.176 | 0.349 | 0.250 | 0.151 | 0.132 | 0.110 |
| | | 2.528 | 1.477 | 1.091 | 0.602 | 0.281 | 0.146 | 0.145 | 0.125 |
| | | 2.027 | 1.452 | 0.826 | 0.468 | 0.260 | 0.124 | 0.127 | 0.098 |
| 3 | rgIns2aa | 2.627 | 1.480 | 1.215 | 0.645 | 0.261 | 0.248 | 0.205 | 0.182 |
| | | 2.377 | 1.679 | 1.008 | 0.527 | 0.241 | 0.269 | 0.179 | 0.162 |
| | | 2.136 | 1.723 | 1.177 | 0.621 | 0.233 | 0.237 | 0.197 | 0.170 |
| 5 | rg2aaΔ-180I | 2.345 | 1.597 | 1.081 | 0.692 | 0.249 | 0.191 | 0.195 | 0.149 |
| | | 2.420 | 1.626 | 1.043 | 0.660 | 0.208 | 0.162 | 0.174 | 0.138 |
| | | 2.068 | 1.332 | 0.972 | 0.588 | 0.212 | 0.214 | 0.166 | 0.144 |
| 6 | rg2aaΔ-498R | 2.158 | 1.144 | 0.876 | 0.690 | 0.515 | 0.202 | 0.164 | 0.134 |
| | | 2.059 | 1.171 | 0.928 | 0.641 | 0.525 | 0.173 | 0.174 | 0.145 |
| | | 2.087 | 1.128 | 0.874 | 0.659 | 0.568 | 0.186 | 0.181 | 0.136 |
| 7 | rg2aaΔ-129D/180I | 2.563 | 1.961 | 0.827 | 0.484 | 0.225 | 0.205 | 0.160 | 0.103 |
| | | 2.150 | 1.894 | 0.803 | 0.350 | 0.285 | 0.200 | 0.172 | 0.152 |
| | | 2.177 | 1.760 | 0.871 | 0.463 | 0.266 | 0.205 | 0.191 | 0.140 |
| 8 | rg2aaΔ-180I/498R | 1.318 | 1.194 | 0.823 | 0.735 | 0.539 | 0.462 | 0.311 | 0.371 |
| | | 1.318 | 1.206 | 0.824 | 0.740 | 0.544 | 0.474 | 0.314 | 0.319 |
| | | 1.352 | 1.209 | 0.867 | 0.629 | 0.572 | 0.463 | 0.362 | 0.360 |
| 10 | rgIns2aa/129D | 2.234 | 1.596 | 0.593 | 0.454 | 0.206 | 0.116 | 0.111 | 0.097 |
| | | 2.246 | 1.721 | 0.689 | 0.432 | 0.200 | 0.151 | 0.154 | 0.137 |
| | | 2.111 | 1.862 | 0.654 | 0.373 | 0.248 | 0.166 | 0.118 | 0.127 |
| 11 | rgIns2aa/180I | 2.803 | 1.714 | 0.992 | 0.495 | 0.272 | 0.201 | 0.150 | 0.123 |
| | | 2.779 | 1.820 | 0.803 | 0.597 | 0.261 | 0.189 | 0.126 | 0.103 |
| | | 2.843 | 1.991 | 0.869 | 0.428 | 0.294 | 0.286 | 0.152 | 0.133 |
| 12 | rgIns2aa/498R | 1.832 | 1.339 | 0.786 | 0.623 | 0.552 | 0.473 | 0.313 | 0.390 |
| | | 1.829 | 1.230 | 0.804 | 0.644 | 0.558 | 0.427 | 0.394 | 0.400 |
| | | 1.750 | 1.361 | 0.763 | 0.622 | 0.563 | 0.443 | 0.325 | 0.367 |
| 13 | rg2aaΔ-129D/180I/498R | 2.875 | 2.269 | 1.198 | 0.970 | 0.538 | 0.231 | 0.124 | 0.095 |
| | | 2.426 | 2.205 | 1.099 | 0.876 | 0.499 | 0.259 | 0.129 | 0.101 |
| | | 2.357 | 2.160 | 1.160 | 0.870 | 0.531 | 0.242 | 0.130 | 0.118 |
| 15 | rgIns2aa/129D/498R | 2.454 | 2.097 | 1.513 | 0.914 | 0.509 | 0.236 | 0.126 | 0.111 |
| | | 2.538 | 2.088 | 1.413 | 0.928 | 0.512 | 0.241 | 0.150 | 0.114 |
| | | 2.587 | 2.076 | 1.648 | 0.881 | 0.596 | 0.218 | 0.132 | 0.126 |
| 16 | rgIns2aa/180I/498R | 2.287 | 1.737 | 1.162 | 0.789 | 0.201 | 0.123 | 0.125 | 0.096 |
| | | 2.397 | 1.772 | 1.011 | 0.869 | 0.185 | 0.120 | 0.126 | 0.084 |
| | | 2.501 | 1.941 | 0.624 | 0.754 | 0.199 | 0.123 | 0.128 | 0.117 |
| 18 | rgWa/19 (3aaΔ) | 2.010 | 1.715 | 1.336 | 0.926 | 0.312 | 0.199 | 0.122 | 0.108 |
| | | 2.023 | 1.706 | 1.224 | 0.766 | 0.301 | 0.212 | 0.132 | 0.109 |
| | | 2.029 | 1.663 | 1.293 | 0.849 | 0.286 | 0.219 | 0.122 | 0.115 |
| 19 | rgIns1aa-164D | 1.527 | 1.307 | 1.020 | 0.920 | 0.700 | 0.492 | 0.352 | 0.408 |
| | | 1.684 | 1.256 | 1.116 | 0.975 | 0.672 | 0.519 | 0.380 | 0.401 |
| | | 1.773 | 1.167 | 1.124 | 0.976 | 0.715 | 0.470 | 0.311 | 0.412 |
| 20 | rgIns3aa | 2.576 | 1.375 | 1.112 | 1.044 | 0.929 | 0.779 | 0.523 | 0.416 |
| | | 2.553 | 1.323 | 1.172 | 1.032 | 0.916 | 0.721 | 0.537 | 0.411 |
| | | 2.653 | 1.552 | 1.164 | 0.977 | 0.882 | 0.753 | 0.572 | 0.442 |
| 21 | rg3aaΔ-133G | 2.108 | 1.159 | 0.917 | 0.784 | 0.713 | 0.556 | 0.342 | 0.210 |
| | | 2.274 | 1.374 | 0.928 | 0.768 | 0.790 | 0.599 | 0.334 | 0.269 |
| | | 2.126 | 1.187 | 0.936 | 0.778 | 0.709 | 0.553 | 0.358 | 0.285 |
| 22 | rg3aaΔ-136K | 1.777 | 1.299 | 0.858 | 0.776 | 0.698 | 0.511 | 0.462 | 0.270 |
| | | 1.885 | 1.356 | 0.856 | 0.707 | 0.676 | 0.599 | 0.494 | 0.384 |
| | | 1.889 | 1.238 | 0.894 | 0.762 | 0.656 | 0.503 | 0.486 | 0.379 |
| 23 | rg3aaΔ-133G/136K | 1.761 | 1.338 | 1.189 | 0.890 | 0.681 | 0.514 | 0.496 | 0.248 |
| | | 1.870 | 1.291 | 1.065 | 0.881 | 0.744 | 0.599 | 0.393 | 0.353 |
| | | 1.701 | 1.248 | 1.023 | 0.895 | 0.681 | 0.513 | 0.393 | 0.342 |

**Table S5. Virus infectivity in MDCK cells after pH treatment in two replicates.**

| No | Recombinant virus | pH | TCID50 | TCID50 |
|---|---|---|---|---|
| 1 | rgBRIS/08_WT | pH5 | 3.16E+03 | 2.21E+03 |
| | | pH5.6 | 1.26E+04 | 2.80E+04 |
| | | pH6 | 3.16E+04 | 3.83E+04 |
| | | pH6.5 | 4.22E+04 | 4.14E+04 |
| | | pH7 | 1.00E+05 | 1.00E+05 |
| 2 | rgCOL/17 (2aaΔ) | pH5 | 4.44E+03 | 4.49E+03 |
| | | pH5.6 | 3.16E+04 | 3.16E+04 |
| | | pH6 | 4.64E+04 | 6.96E+04 |
| | | pH6.5 | 7.39E+04 | 7.29E+04 |
| | | pH7 | 1.00E+05 | 1.00E+05 |
| 3 | rgIns2aa | pH5 | 1.00E+04 | 1.78E+04 |
| | | pH5.6 | 1.58E+04 | 1.90E+04 |
| | | pH6 | 2.15E+04 | 2.00E+04 |
| | | pH6.5 | 3.16E+04 | 2.91E+04 |
| | | pH7 | 1.00E+05 | 1.00E+05 |
| 5 | rg2aaΔ-180I | pH5 | 1.70E+04 | 3.35E+04 |
| | | pH5.6 | 3.16E+04 | 3.69E+04 |
| | | pH6 | 4.64E+04 | 4.73E+04 |
| | | pH6.5 | 6.31E+04 | 6.11E+04 |
| | | pH7 | 1.00E+05 | 1.00E+05 |
| 6 | rg2aaΔ-498R | pH5 | 3.16E+03 | 2.00E+03 |
| | | pH5.6 | 2.98E+04 | 2.58E+04 |
| | | pH6 | 3.16E+04 | 4.56E+04 |
| | | pH6.5 | 6.81E+04 | 5.44E+04 |
| | | pH7 | 1.00E+05 | 1.00E+05 |
| 7 | rg2aaΔ-129D/180I | pH5 | 1.93E+04 | 1.93E+04 |
| | | pH5.6 | 3.16E+04 | 3.16E+04 |
| | | pH6 | 5.88E+04 | 5.25E+04 |
| | | pH6.5 | 8.12E+04 | 7.46E+04 |
| | | pH7 | 1.00E+05 | 1.00E+05 |
| 8 | rg2aaΔ-180I/498R | pH5 | 2.77E+03 | 5.10E+03 |
| | | pH5.6 | 5.88E+03 | 8.24E+03 |
| | | pH6 | 3.16E+04 | 2.69E+04 |
| | | pH6.5 | 6.71E+04 | 7.63E+04 |
| | | pH7 | 1.00E+05 | 1.00E+05 |
| 10 | rgIns2aa/129D | pH5 | 6.31E+03 | 6.38E+03 |
| | | pH5.6 | 3.16E+04 | 3.62E+04 |
| | | pH6 | 4.64E+04 | 5.18E+04 |
| | | pH6.5 | 7.50E+04 | 7.73E+04 |
| | | pH7 | 1.00E+05 | 1.00E+05 |
| 11 | rgIns2aa/180I | pH5 | 5.62E+03 | 3.75E+03 |
| | | pH5.6 | 1.58E+04 | 1.43E+04 |
| | | pH6 | 3.73E+04 | 3.07E+04 |
| | | pH6.5 | 5.46E+04 | 5.35E+04 |
| | | pH7 | 1.00E+05 | 1.00E+05 |
| 12 | rgIns2aa/498R | pH5 | 3.16E+03 | 1.96E+03 |
| | | pH5.6 | 3.16E+04 | 3.16E+04 |
| | | pH6 | 6.31E+04 | 6.40E+04 |
| | | pH6.5 | 7.63E+04 | 7.28E+04 |
| | | pH7 | 1.00E+05 | 1.00E+05 |
| 13 | rg2aaΔ-129D/180I/498R | pH5 | 3.16E+03 | 2.00E+03 |
| | | pH5.6 | 4.64E+03 | 4.27E+03 |
| | | pH6 | 2.68E+04 | 2.61E+04 |
| | | pH6.5 | 6.31E+04 | 5.32E+04 |
| | | pH7 | 1.00E+05 | 1.00E+05 |
| 15 | rgIns2aa/129D/498R | pH5 | 6.31E+03 | 4.39E+03 |
| | | pH5.6 | 1.89E+04 | 1.97E+04 |
| | | pH6 | 2.15E+04 | 2.61E+04 |

| | | | | |
|---|---|---|---|---|
| | | pH6.5 | 3.73E+04 | 4.75E+04 |
| | | pH7 | 1.00E+05 | 1.00E+05 |
| 16 | rgIns2aa/180I/498R | pH5 | 1.31E+04 | 1.00E+04 |
| | | pH5.6 | 5.58E+04 | 6.44E+04 |
| | | pH6 | 7.65E+04 | 7.10E+04 |
| | | pH6.5 | 8.25E+04 | 8.27E+04 |
| | | pH7 | 1.00E+05 | 1.00E+05 |
| 18 | rgWa/19 (3aaΔ) | pH5 | 1.54E+04 | 2.13E+04 |
| | | pH5.6 | 2.67E+04 | 2.94E+04 |
| | | pH6 | 4.96E+04 | 5.62E+04 |
| | | pH6.5 | 6.42E+04 | 7.63E+04 |
| | | pH7 | 1.00E+05 | 1.00E+05 |
| 19 | rgIns1aa-164D | pH5 | 1.25E+04 | 1.31E+04 |
| | | pH5.6 | 1.90E+04 | 1.85E+04 |
| | | pH6 | 2.15E+04 | 2.37E+04 |
| | | pH6.5 | 7.50E+04 | 7.87E+04 |
| | | pH7 | 1.00E+05 | 1.00E+05 |
| 20 | rgIns3aa | pH5 | 1.93E+04 | 1.36E+04 |
| | | pH5.6 | 2.51E+04 | 2.29E+04 |
| | | pH6 | 4.11E+04 | 5.46E+04 |
| | | pH6.5 | 6.66E+04 | 7.80E+04 |
| | | pH7 | 1.00E+05 | 1.00E+05 |
| 21 | rg3aaΔ-133G | pH5 | 1.96E+04 | 1.58E+04 |
| | | pH5.6 | 2.57E+04 | 2.92E+04 |
| | | pH6 | 3.94E+04 | 3.11E+04 |
| | | pH6.5 | 6.31E+04 | 6.96E+04 |
| | | pH7 | 1.00E+05 | 1.00E+05 |
| 22 | rg3aaΔ-136K | pH5 | 1.58E+04 | 1.29E+04 |
| | | pH5.6 | 2.68E+04 | 3.31E+04 |
| | | pH6 | 5.50E+04 | 5.88E+04 |
| | | pH6.5 | 7.17E+04 | 7.28E+04 |
| | | pH7 | 1.00E+05 | 1.00E+05 |
| 23 | rg3aaΔ-133G/136K | pH5 | 1.78E+04 | 1.25E+04 |
| | | pH5.6 | 2.68E+04 | 2.94E+04 |
| | | pH6 | 4.64E+04 | 4.99E+04 |
| | | pH6.5 | 7.94E+04 | 6.46E+04 |
| | | pH7 | 1.00E+05 | 1.00E+05 |

**Table S6. Haemagglutination activity after temperature incubation in three replicates.**

| No | Recombinant virus | 33°C | 35°C | 40°C | 45°C | 50°C | 55°C | 60°C | 65°C |
|---|---|---|---|---|---|---|---|---|---|
| 1 | rgBRIS/08_WT | 64 | 64 | 32 | 32 | 16 | 8 | 8 | 0 |
| | | 64 | 64 | 64 | 32 | 32 | 16 | 8 | 0 |
| | | 64 | 64 | 32 | 32 | 16 | 8 | 8 | 0 |
| 2 | rgCOL/17 (2aaΔ) | 64 | 64 | 64 | 32 | 32 | 16 | 8 | 0 |
| | | 64 | 64 | 64 | 64 | 32 | 16 | 8 | 0 |
| | | 64 | 64 | 64 | 64 | 32 | 16 | 8 | 0 |
| 3 | rgIns2aa | 64 | 64 | 32 | 32 | 32 | 16 | 8 | 0 |
| | | 64 | 64 | 32 | 32 | 32 | 16 | 8 | 0 |
| | | 64 | 64 | 32 | 32 | 32 | 16 | 8 | 0 |
| 5 | rg2aaΔ-180I | 64 | 64 | 64 | 32 | 32 | 16 | 8 | 0 |
| | | 64 | 32 | 32 | 32 | 32 | 16 | 8 | 0 |
| | | 64 | 32 | 32 | 32 | 32 | 16 | 8 | 0 |
| 6 | rg2aaΔ-498R | 64 | 64 | 32 | 32 | 32 | 16 | 8 | 0 |
| | | 64 | 32 | 32 | 32 | 32 | 16 | 8 | 0 |
| | | 64 | 32 | 32 | 32 | 32 | 16 | 8 | 0 |
| 7 | rg2aaΔ-129D/180I | 64 | 64 | 64 | 64 | 32 | 16 | 8 | 0 |
| | | 64 | 64 | 64 | 64 | 32 | 16 | 8 | 0 |
| | | 64 | 64 | 64 | 64 | 32 | 16 | 8 | 0 |
| 8 | rg2aaΔ-180I/498R | 64 | 64 | 64 | 32 | 32 | 16 | 8 | 0 |
| | | 64 | 64 | 32 | 32 | 32 | 16 | 8 | 0 |
| | | 64 | 64 | 32 | 32 | 32 | 16 | 8 | 0 |
| 10 | rgIns2aa/129D | 64 | 64 | 32 | 32 | 16 | 8 | 4 | 0 |
| | | 64 | 64 | 64 | 32 | 32 | 16 | 8 | 0 |
| | | 64 | 64 | 64 | 32 | 32 | 16 | 8 | 0 |
| 11 | rgIns2aa/180I | 64 | 64 | 64 | 32 | 32 | 16 | 8 | 0 |
| | | 64 | 32 | 32 | 32 | 16 | 8 | 4 | 0 |
| | | 64 | 32 | 32 | 32 | 16 | 8 | 4 | 0 |
| 12 | rgIns2aa/498R | 64 | 64 | 32 | 32 | 16 | 16 | 4 | 0 |
| | | 64 | 64 | 64 | 32 | 32 | 16 | 4 | 0 |
| | | 64 | 64 | 64 | 32 | 32 | 16 | 4 | 0 |
| 13 | rg2aaΔ-129D/180I/498R | 64 | 32 | 32 | 32 | 16 | 8 | 8 | 0 |
| | | 64 | 32 | 32 | 32 | 16 | 8 | 8 | 0 |
| | | 64 | 32 | 32 | 32 | 16 | 8 | 8 | 0 |
| 15 | rgIns2aa/129D/498R | 64 | 64 | 32 | 32 | 32 | 16 | 8 | 0 |
| | | 64 | 64 | 64 | 32 | 32 | 16 | 8 | 0 |
| | | 64 | 64 | 32 | 32 | 32 | 16 | 8 | 0 |
| 16 | rgIns2aa/180I/498R | 64 | 64 | 32 | 32 | 16 | 16 | 8 | 0 |
| | | 64 | 64 | 64 | 64 | 16 | 16 | 8 | 0 |
| | | 64 | 64 | 32 | 32 | 16 | 16 | 8 | 0 |
| 18 | rgWa/19 (3aaΔ) | 64 | 64 | 32 | 32 | 16 | 8 | 4 | 0 |
| | | 64 | 64 | 32 | 32 | 32 | 8 | 4 | 0 |
| | | 64 | 64 | 32 | 32 | 32 | 8 | 4 | 0 |
| 19 | rgIns1aa-164D | 64 | 64 | 32 | 32 | 32 | 8 | 4 | 0 |
| | | 64 | 64 | 32 | 32 | 32 | 8 | 4 | 0 |
| | | 64 | 64 | 32 | 32 | 32 | 8 | 4 | 0 |
| 20 | rgIns3aa | 64 | 32 | 32 | 16 | 16 | 8 | 4 | 0 |
| | | 64 | 32 | 32 | 16 | 16 | 8 | 4 | 0 |
| | | 64 | 32 | 32 | 16 | 16 | 8 | 4 | 0 |
| 21 | rg3aaΔ-133G | 64 | 64 | 32 | 32 | 16 | 8 | 4 | 0 |
| | | 64 | 64 | 32 | 32 | 16 | 8 | 4 | 0 |
| | | 64 | 64 | 32 | 32 | 16 | 8 | 4 | 0 |
| 22 | rg3aaΔ-136K | 64 | 64 | 32 | 32 | 16 | 8 | 4 | 0 |
| | | 64 | 64 | 64 | 32 | 16 | 8 | 4 | 0 |
| | | 64 | 64 | 32 | 32 | 16 | 8 | 4 | 0 |
| 23 | rg3aaΔ-133G/136K | 64 | 64 | 32 | 32 | 16 | 8 | 4 | 0 |
| | | 64 | 32 | 32 | 16 | 16 | 8 | 4 | 0 |
| | | 64 | 64 | 32 | 32 | 16 | 8 | 4 | 0 |

**Table S7. Haemagglutination activity following incubation at 50°C for varying durations in three replicates.**

| No | Recombinant virus | 0H | 1H | 2H | 3H | 4H | 5H | 6H | 12H |
|---|---|---|---|---|---|---|---|---|---|
| 1 | rgBRIS/08_WT | 64 | 16 | 16 | 16 | 16 | 16 | 8 | 0 |
| | | 64 | 16 | 8 | 8 | 8 | 4 | 4 | 0 |
| | | 64 | 16 | 16 | 16 | 8 | 8 | 4 | 0 |
| 2 | rgCOL/17 (2aaΔ) | 64 | 16 | 16 | 16 | 16 | 16 | 8 | 0 |
| | | 64 | 16 | 16 | 16 | 16 | 8 | 4 | 0 |
| | | 64 | 16 | 16 | 16 | 16 | 16 | 8 | 0 |
| 3 | rgIns2aa | 64 | 16 | 16 | 4 | 4 | 4 | 0 | 0 |
| | | 64 | 16 | 16 | 8 | 8 | 8 | 4 | 0 |
| | | 64 | 16 | 16 | 4 | 4 | 4 | 0 | 0 |
| 5 | rg2aaΔ-180I | 64 | 32 | 16 | 8 | 8 | 4 | 4 | 0 |
| | | 64 | 16 | 16 | 8 | 8 | 8 | 4 | 0 |
| | | 64 | 16 | 16 | 8 | 8 | 8 | 4 | 0 |
| 6 | rg2aaΔ-498R | 64 | 16 | 16 | 16 | 16 | 8 | 8 | 0 |
| | | 64 | 16 | 16 | 16 | 16 | 16 | 8 | 0 |
| | | 64 | 16 | 16 | 16 | 16 | 16 | 8 | 0 |
| 7 | rg2aaΔ-129D/180I | 64 | 32 | 32 | 16 | 16 | 8 | 8 | 0 |
| | | 64 | 16 | 16 | 16 | 16 | 16 | 8 | 0 |
| | | 64 | 32 | 32 | 16 | 16 | 8 | 8 | 0 |
| 8 | rg2aaΔ-180I/498R | 64 | 16 | 16 | 16 | 8 | 8 | 4 | 0 |
| | | 64 | 16 | 16 | 16 | 16 | 16 | 8 | 0 |
| | | 64 | 16 | 16 | 16 | 16 | 16 | 8 | 0 |
| 10 | rgIns2aa/129D | 64 | 16 | 16 | 16 | 8 | 8 | 4 | 0 |
| | | 64 | 16 | 16 | 16 | 16 | 16 | 8 | 0 |
| | | 64 | 16 | 16 | 16 | 16 | 16 | 8 | 0 |
| 11 | rgIns2aa/180I | 64 | 16 | 8 | 8 | 8 | 4 | 4 | 0 |
| | | 64 | 8 | 8 | 8 | 8 | 4 | 4 | 0 |
| | | 64 | 8 | 8 | 8 | 8 | 4 | 4 | 0 |
| 12 | rgIns2aa/498R | 64 | 32 | 16 | 16 | 8 | 8 | 4 | 0 |
| | | 64 | 16 | 16 | 16 | 16 | 16 | 8 | 0 |
| | | 64 | 16 | 16 | 16 | 16 | 16 | 8 | 0 |
| 13 | rg2aaΔ-129D/180I/498R | 64 | 32 | 16 | 16 | 16 | 8 | 8 | 0 |
| | | 64 | 8 | 8 | 8 | 8 | 4 | 4 | 0 |
| | | 64 | 32 | 16 | 16 | 16 | 8 | 8 | 0 |
| 15 | rgIns2aa/129D/498R | 64 | 16 | 16 | 16 | 16 | 8 | 8 | 0 |
| | | 64 | 16 | 8 | 8 | 8 | 8 | 4 | 0 |
| | | 64 | 16 | 8 | 8 | 8 | 8 | 4 | 0 |
| 16 | rgIns2aa/180I/498R | 64 | 16 | 8 | 8 | 4 | 4 | 0 | 0 |
| | | 64 | 16 | 16 | 8 | 8 | 8 | 4 | 0 |
| | | 64 | 16 | 16 | 8 | 8 | 8 | 4 | 0 |
| 18 | rgWa/19 (3aaΔ) | 64 | 32 | 32 | 32 | 16 | 8 | 4 | 0 |
| | | 64 | 32 | 32 | 16 | 8 | 8 | 4 | 0 |
| | | 64 | 32 | 32 | 16 | 8 | 8 | 4 | 0 |
| 19 | rgIns1aa-164D | 64 | 32 | 32 | 32 | 16 | 16 | 8 | 0 |
| | | 64 | 32 | 32 | 16 | 8 | 4 | 4 | 0 |
| | | 64 | 32 | 32 | 16 | 8 | 4 | 4 | 0 |
| 20 | rgIns3aa | 64 | 32 | 16 | 16 | 8 | 8 | 4 | 0 |
| | | 64 | 32 | 32 | 16 | 8 | 4 | 4 | 0 |
| | | 64 | 32 | 32 | 16 | 8 | 4 | 4 | 0 |
| 21 | rg3aaΔ-133G | 64 | 32 | 16 | 16 | 8 | 8 | 4 | 0 |
| | | 64 | 32 | 32 | 16 | 8 | 8 | 4 | 0 |
| | | 64 | 32 | 16 | 16 | 8 | 8 | 4 | 0 |
| 22 | rg3aaΔ-136K | 64 | 32 | 32 | 32 | 8 | 8 | 4 | 0 |

| | | | | | | | | |
|---|---|---|---|---|---|---|---|---|
| | 64 | 32 | 32 | 16 | 8 | 8 | 4 | 0 |
| | 64 | 32 | 32 | 32 | 8 | 8 | 4 | 0 |
| 23 rg3aaΔ-133G/136K | 64 | 32 | 32 | 16 | 16 | 8 | 4 | 0 |
| | 64 | 32 | 32 | 16 | 8 | 8 | 4 | 0 |
| | 64 | 32 | 32 | 16 | 16 | 8 | 4 | 0 |

**Table S8. Statistical analysis of significance differences among viruses treated at pH 5.**

| Recombinant virus | rgIns2aa/129D/498R | rgIns2aa/129D | rgIns2aa/180I/498R | rgIns2aa/180I | rgIns2aa/498R | rg2aaΔ-129D/180I | rg2aaΔ-129D/180I/498R | rg2aaΔ-180I | rg2aaΔ-180I/498R | rg2aaΔ-498R | rg3aaΔ-133G | rg3aaΔ-133G/136K | rg3aaΔ-136K | rgIns1aa-164D | rgIns2aa | rgIns3aa | rgBRIS/08 wild-type | rgCOL/17 (2aaΔ) | rgWA/19 (3aaΔ) |
|---|---|---|---|---|---|---|---|---|---|---|---|---|---|---|---|---|---|---|---|
| rgIns2aa/129D/498R | | 0.999 | 0.000 | 1.000 | 0.000 | 0.000 | 0.000 | 0.000 | 0.634 | 0.000 | 0.000 | 0.000 | 0.000 | 0.000 | 0.000 | 0.000 | 0.000 | 1.000 | 0.000 |
| rgIns2aa/129D | 0.999 | | 0.025 | 0.746 | 0.000 | 0.000 | 0.000 | 0.000 | 0.041 | 0.000 | 0.000 | 0.000 | 0.000 | 0.002 | 0.001 | 0.000 | 0.000 | 0.619 | 0.000 |
| rgIns2aa/180I/498R | 0.000 | 0.025 | | 0.000 | 0.000 | 0.144 | 0.000 | 0.003 | 0.000 | 0.000 | 0.443 | 0.980 | 0.996 | 1.000 | 1.000 | 0.797 | 0.000 | 0.000 | 0.334 |
| rgIns2aa/180I | 1.000 | 0.746 | 0.000 | | 0.003 | 0.000 | 0.004 | 0.000 | 0.994 | 0.004 | 0.000 | 0.000 | 0.000 | 0.000 | 0.000 | 0.000 | 0.015 | 1.000 | 0.000 |
| rgIns2aa/498R | 0.000 | 0.000 | 0.000 | 0.003 | | 0.000 | 1.000 | 0.000 | 0.226 | 1.000 | 0.000 | 0.000 | 0.000 | 0.000 | 0.000 | 0.000 | 1.000 | 0.006 | 0.000 |
| rg2aaΔ-129D/180I | 0.000 | 0.000 | 0.144 | 0.000 | 0.000 | | 0.000 | 0.999 | 0.000 | 0.000 | 1.000 | 0.984 | 0.940 | 0.547 | 0.716 | 1.000 | 0.000 | 0.000 | 1.000 |
| rg2aaΔ-129D/180I/498R | 0.000 | 0.000 | 0.000 | 0.004 | 1.000 | 0.000 | | 0.000 | 0.265 | 1.000 | 0.000 | 0.000 | 0.000 | 0.000 | 0.000 | 0.000 | 1.000 | 0.008 | 0.000 |
| rg2aaΔ-180I | 0.000 | 0.000 | 0.003 | 0.000 | 0.000 | 0.999 | 0.000 | | 0.000 | 0.000 | 0.953 | 0.343 | 0.211 | 0.034 | 0.068 | 0.716 | 0.000 | 0.000 | 0.981 |
| rg2aaΔ-180I/498R | 0.634 | 0.041 | 0.000 | 0.994 | 0.226 | 0.000 | 0.265 | 0.000 | | 0.265 | 0.000 | 0.000 | 0.000 | 0.000 | 0.000 | 0.000 | 0.522 | 0.999 | 0.000 |
| rg2aaΔ-498R | 0.000 | 0.000 | 0.000 | 0.004 | 1.000 | 0.000 | 1.000 | 0.000 | 0.265 | | 0.000 | 0.000 | 0.000 | 0.000 | 0.000 | 0.000 | 1.000 | 0.007 | 0.000 |
| rg3aaΔ-133G | 0.000 | 0.000 | 0.443 | 0.000 | 0.000 | 1.000 | 0.000 | 0.953 | 0.000 | 0.000 | | 1.000 | 0.999 | 0.894 | 0.964 | 1.000 | 0.000 | 0.000 | 1.000 |
| rg3aaΔ-133G/136K | 0.000 | 0.000 | 0.980 | 0.000 | 0.000 | 0.984 | 0.000 | 0.343 | 0.000 | 0.000 | 1.000 | | 1.000 | 1.000 | 1.000 | 1.000 | 0.000 | 0.000 | 0.999 |
| rg3aaΔ-136K | 0.000 | 0.000 | 0.996 | 0.000 | 0.000 | 0.940 | 0.000 | 0.211 | 0.000 | 0.000 | 0.999 | 1.000 | | 1.000 | 1.000 | 1.000 | 0.000 | 0.000 | 0.995 |
| rgIns1aa-164D | 0.000 | 0.002 | 1.000 | 0.000 | 0.000 | 0.547 | 0.000 | 0.034 | 0.000 | 0.000 | 0.894 | 1.000 | 1.000 | | 1.000 | 0.994 | 0.000 | 0.000 | 0.814 |
| rgIns2aa | 0.000 | 0.001 | 1.000 | 0.000 | 0.000 | 0.716 | 0.000 | 0.068 | 0.000 | 0.000 | 0.964 | 1.000 | 1.000 | 1.000 | | 0.999 | 0.000 | 0.000 | 0.921 |
| rgIns3aa | 0.000 | 0.000 | 0.797 | 0.000 | 0.000 | 1.000 | 0.000 | 0.716 | 0.000 | 0.000 | 1.000 | 1.000 | 1.000 | 0.994 | 0.999 | | 0.000 | 0.000 | 1.000 |
| rgBRIS/08 wild-type | 0.000 | 0.000 | 0.000 | 0.015 | 1.000 | 0.000 | 1.000 | 0.000 | 0.522 | 1.000 | 0.000 | 0.000 | 0.000 | 0.000 | 0.000 | 0.000 | | 0.027 | 0.000 |
| rgCOL/17 (2aaΔ) | 1.000 | 0.619 | 0.000 | 1.000 | 0.006 | 0.000 | 0.008 | 0.000 | 0.999 | 0.007 | 0.000 | 0.000 | 0.000 | 0.000 | 0.000 | 0.000 | 0.027 | | 0.000 |
| rgWA/19 (3aaΔ) | 0.000 | 0.000 | 0.334 | 0.000 | 0.000 | 1.000 | 0.000 | 0.981 | 0.000 | 0.000 | 1.000 | 0.999 | 0.995 | 0.814 | 0.921 | 1.000 | 0.000 | 0.000 | |

**Table S9. Statistical analysis of significance differences among viruses treated at pH 5.6.**

| Recombinant virus | rgIns2aa/129D/498R | rgIns2aa/129D | rgIns2aa/180I/498R | rgIns2aa/180I | rgIns2aa/498R | rg2aaΔ-129D/180I | rg2aaΔ-129D/180I/498R | rg2aaΔ-180I | rg2aaΔ-180I/498R | rg2aaΔ-498R | rg3aaΔ-133G | rg3aaΔ-133G/136K | rg3aaΔ-136K | rgIns1aa-164D | rgIns2aa | rgIns3aa | rgBRIS/08 wild-type | rgCOL/17 (2aaΔ) | rgWA/19 (3aaΔ) |
|---|---|---|---|---|---|---|---|---|---|---|---|---|---|---|---|---|---|---|---|
| rgIns2aa/129D/498R | | 0.126 | 0.000 | 0.992 | 0.297 | 0.297 | 0.000 | 0.110 | 0.000 | 0.812 | 0.851 | 0.764 | 0.526 | 1.000 | 1.000 | 0.999 | 1.000 | 0.297 | 0.774 |
| rgIns2aa/129D | 0.126 | | 0.170 | 0.001 | 1.000 | 1.000 | 0.000 | 1.000 | 0.000 | 1.000 | 0.999 | 1.000 | 1.000 | 0.084 | 0.020 | 0.887 | 0.072 | 1.000 | 1.000 |
| rgIns2aa/180I/498R | 0.000 | 0.170 | | 0.000 | 0.063 | 0.063 | 0.000 | 0.191 | 0.000 | 0.006 | 0.004 | 0.007 | 0.023 | 0.000 | 0.000 | 0.000 | 0.000 | 0.063 | 0.007 |
| rgIns2aa/180I | 0.992 | 0.001 | 0.000 | | 0.004 | 0.004 | 0.000 | 0.001 | 0.000 | 0.048 | 0.059 | 0.038 | 0.013 | 0.998 | 1.000 | 0.359 | 0.999 | 0.004 | 0.040 |
| rgIns2aa/498R | 0.297 | 1.000 | 0.063 | 0.004 | | 1.000 | 0.000 | 1.000 | 0.000 | 1.000 | 1.000 | 1.000 | 1.000 | 0.215 | 0.063 | 0.983 | 0.190 | 1.000 | 1.000 |
| rg2aaΔ-129D/180I | 0.297 | 1.000 | 0.063 | 0.004 | 1.000 | | 0.000 | 1.000 | 0.000 | 1.000 | 1.000 | 1.000 | 1.000 | 0.215 | 0.063 | 0.983 | 0.190 | 1.000 | 1.000 |
| rg2aaΔ-129D/180I/498R | 0.000 | 0.000 | 0.000 | 0.000 | 0.000 | 0.000 | | 0.000 | 0.211 | 0.000 | 0.000 | 0.000 | 0.000 | 0.000 | 0.000 | 0.000 | 0.000 | 0.000 | 0.000 |
| rg2aaΔ-180I | 0.110 | 1.000 | 0.191 | 0.001 | 1.000 | 1.000 | 0.000 | | 0.000 | 0.999 | 0.999 | 1.000 | 1.000 | 0.073 | 0.017 | 0.863 | 0.063 | 1.000 | 1.000 |
| rg2aaΔ-180I/498R | 0.000 | 0.000 | 0.000 | 0.000 | 0.000 | 0.000 | 0.211 | 0.000 | | 0.000 | 0.000 | 0.000 | 0.000 | 0.000 | 0.000 | 0.000 | 0.000 | 0.000 | 0.000 |
| rg2aaΔ-498R | 0.812 | 1.000 | 0.006 | 0.048 | 1.000 | 1.000 | 0.000 | 0.999 | 0.000 | | 1.000 | 1.000 | 1.000 | 0.714 | 0.367 | 1.000 | 0.676 | 1.000 | 1.000 |
| rg3aaΔ-133G | 0.851 | 0.999 | 0.004 | 0.059 | 1.000 | 1.000 | 0.000 | 0.999 | 0.000 | 1.000 | | 1.000 | 1.000 | 0.761 | 0.415 | 1.000 | 0.725 | 1.000 | 1.000 |
| rg3aaΔ-133G/136K | 0.764 | 1.000 | 0.007 | 0.038 | 1.000 | 1.000 | 0.000 | 1.000 | 0.000 | 1.000 | 1.000 | | 1.000 | 0.658 | 0.317 | 1.000 | 0.619 | 1.000 | 1.000 |
| rg3aaΔ-136K | 0.526 | 1.000 | 0.023 | 0.013 | 1.000 | 1.000 | 0.000 | 1.000 | 0.000 | 1.000 | 1.000 | 1.000 | | 0.415 | 0.154 | 0.999 | 0.378 | 1.000 | 1.000 |
| rgIns1aa-164D | 1.000 | 0.084 | 0.000 | 0.998 | 0.215 | 0.215 | 0.000 | 0.073 | 0.000 | 0.714 | 0.761 | 0.658 | 0.415 | | 1.000 | 0.994 | 1.000 | 0.215 | 0.669 |
| rgIns2aa | 1.000 | 0.020 | 0.000 | 1.000 | 0.063 | 0.063 | 0.000 | 0.017 | 0.000 | 0.367 | 0.415 | 0.317 | 0.154 | 1.000 | | 0.905 | 1.000 | 0.063 | 0.326 |
| rgIns3aa | 0.999 | 0.887 | 0.000 | 0.359 | 0.983 | 0.983 | 0.000 | 0.863 | 0.000 | 1.000 | 1.000 | 1.000 | 0.999 | 0.994 | 0.905 | | 0.991 | 0.983 | 1.000 |
| rgBRIS/08 wild-type | 1.000 | 0.072 | 0.000 | 0.999 | 0.190 | 0.190 | 0.000 | 0.063 | 0.000 | 0.676 | 0.725 | 0.619 | 0.378 | 1.000 | 1.000 | 0.991 | | 0.190 | 0.630 |
| rgCOL/17 (2aaΔ) | 0.297 | 1.000 | 0.063 | 0.004 | 1.000 | 1.000 | 0.000 | 1.000 | 0.000 | 1.000 | 1.000 | 1.000 | 1.000 | 0.215 | 0.063 | 0.983 | 0.190 | | 1.000 |
| rgWA/19 (3aaΔ) | 0.774 | 1.000 | 0.007 | 0.040 | 1.000 | 1.000 | 0.000 | 1.000 | 0.000 | 1.000 | 1.000 | 1.000 | 1.000 | 0.669 | 0.326 | 1.000 | 0.630 | 1.000 | |

**Table S10. Statistical analysis of significance differences among viruses treated at pH 6.**

| Recombinant virus | rgIns2aa/129D/498R | rgIns2aa/129D | rgIns2aa/180I/498R | rgIns2aa/180I | rgIns2aa/498R | rg2aaΔ-129D/180I | rg2aaΔ-129D/180I/498R | rg2aaΔ-180I | rg2aaΔ-180I/498R | rg2aaΔ-498R | rg3aaΔ-133G | rg3aaΔ-133G/136K | rg3aaΔ-136K | rgIns1aa-164D | rgIns2aa | rgIns3aa | rgBRIS/08 wild-type | rgCOL/17 (2aaΔ) | rgWA/19 (3aaΔ) |
|---|---|---|---|---|---|---|---|---|---|---|---|---|---|---|---|---|---|---|---|
| rgIns2aa/129D/498R | | 0.010 | 0.000 | 0.854 | 0.000 | 0.001 | 1.000 | 0.023 | 0.999 | 0.425 | 0.746 | 0.015 | 0.001 | 1.000 | 1.000 | 0.020 | 0.766 | 0.001 | 0.002 |
| rgIns2aa/129D | 0.010 | | 0.777 | 0.849 | 0.996 | 1.000 | 0.075 | 1.000 | 0.286 | 0.995 | 0.927 | 1.000 | 1.000 | 0.004 | 0.001 | 1.000 | 0.916 | 1.000 | 1.000 |
| rgIns2aa/180I/498R | 0.000 | 0.777 | | 0.006 | 1.000 | 0.991 | 0.000 | 0.607 | 0.000 | 0.050 | 0.012 | 0.709 | 0.997 | 0.000 | 0.000 | 0.646 | 0.011 | 0.996 | 0.952 |
| rgIns2aa/180I | 0.854 | 0.849 | 0.006 | | 0.077 | 0.390 | 0.996 | 0.944 | 1.000 | 1.000 | 1.000 | 0.896 | 0.308 | 0.683 | 0.325 | 0.928 | 1.000 | 0.316 | 0.584 |
| rgIns2aa/498R | 0.000 | 0.996 | 1.000 | 0.077 | | 1.000 | 0.001 | 0.977 | 0.005 | 0.335 | 0.125 | 0.991 | 1.000 | 0.000 | 0.000 | 0.984 | 0.116 | 1.000 | 1.000 |
| rg2aaΔ-129D/180I | 0.001 | 1.000 | 0.991 | 0.390 | 1.000 | | 0.009 | 1.000 | 0.054 | 0.828 | 0.523 | 1.000 | 1.000 | 0.000 | 0.000 | 1.000 | 0.500 | 1.000 | 1.000 |
| rg2aaΔ-129D/180I/498R | 1.000 | 0.075 | 0.000 | 0.996 | 0.001 | 0.009 | | 0.142 | 1.000 | 0.857 | 0.983 | 0.099 | 0.006 | 1.000 | 0.995 | 0.124 | 0.986 | 0.006 | 0.023 |
| rg2aaΔ-180I | 0.023 | 1.000 | 0.607 | 0.944 | 0.977 | 1.000 | 0.142 | | 0.444 | 1.000 | 0.980 | 1.000 | 1.000 | 0.009 | 0.001 | 1.000 | 0.976 | 1.000 | 1.000 |
| rg2aaΔ-180I/498R | 0.999 | 0.286 | 0.000 | 1.000 | 0.005 | 0.054 | 1.000 | 0.444 | | 0.992 | 1.000 | 0.348 | 0.037 | 0.992 | 0.880 | 0.407 | 1.000 | 0.039 | 0.114 |
| rg2aaΔ-498R | 0.425 | 0.995 | 0.050 | 1.000 | 0.335 | 0.828 | 0.857 | 1.000 | 0.992 | | 1.000 | 0.998 | 0.750 | 0.253 | 0.073 | 0.999 | 1.000 | 0.759 | 0.940 |
| rg3aaΔ-133G | 0.746 | 0.927 | 0.012 | 1.000 | 0.125 | 0.523 | 0.983 | 0.980 | 1.000 | 1.000 | | 0.955 | 0.430 | 0.547 | 0.222 | 0.972 | 1.000 | 0.440 | 0.719 |
| rg3aaΔ-133G/136K | 0.015 | 1.000 | 0.709 | 0.896 | 0.991 | 1.000 | 0.099 | 1.000 | 0.348 | 0.998 | 0.955 | | 1.000 | 0.006 | 0.001 | 1.000 | 0.948 | 1.000 | 1.000 |
| rg3aaΔ-136K | 0.001 | 1.000 | 0.997 | 0.308 | 1.000 | 1.000 | 0.006 | 1.000 | 0.037 | 0.750 | 0.430 | 1.000 | | 0.000 | 0.000 | 1.000 | 0.408 | 1.000 | 1.000 |
| rgIns1aa-164D | 1.000 | 0.004 | 0.000 | 0.683 | 0.000 | 0.000 | 1.000 | 0.009 | 0.992 | 0.253 | 0.547 | 0.006 | 0.000 | | 1.000 | 0.008 | 0.570 | 0.000 | 0.001 |
| rgIns2aa | 1.000 | 0.001 | 0.000 | 0.325 | 0.000 | 0.000 | 0.995 | 0.001 | 0.880 | 0.073 | 0.222 | 0.001 | 0.000 | 1.000 | | 0.001 | 0.237 | 0.000 | 0.000 |
| rgIns3aa | 0.020 | 1.000 | 0.646 | 0.928 | 0.984 | 1.000 | 0.124 | 1.000 | 0.407 | 0.999 | 0.972 | 1.000 | 1.000 | 0.008 | 0.001 | | 0.967 | 1.000 | 1.000 |
| rgBRIS/08 wild-type | 0.766 | 0.916 | 0.011 | 1.000 | 0.116 | 0.500 | 0.986 | 0.976 | 1.000 | 1.000 | 1.000 | 0.948 | 0.408 | 0.570 | 0.237 | 0.967 | | 0.418 | 0.697 |
| rgCOL/17 (2aaΔ) | 0.001 | 1.000 | 0.996 | 0.316 | 1.000 | 1.000 | 0.006 | 1.000 | 0.039 | 0.759 | 0.440 | 1.000 | 1.000 | 0.000 | 0.000 | 1.000 | 0.418 | | 1.000 |
| rgWA/19 (3aaΔ) | 0.002 | 1.000 | 0.952 | 0.584 | 1.000 | 1.000 | 0.023 | 1.000 | 0.114 | 0.940 | 0.719 | 1.000 | 1.000 | 0.001 | 0.000 | 1.000 | 0.697 | 1.000 | |

**Table S11. Statistical analysis of significance differences among viruses treated at pH 6.5.**

| Recombinant virus | rgIns2aa/129D/498R | rgIns2aa/129D | rgIns2aa/180I/498R | rgIns2aa/180I | rgIns2aa/498R | rg2aaΔ-129D/180I | rg2aaΔ-129D/180I/498R | rg2aaΔ-180I | rg2aaΔ-180I/498R | rg2aaΔ-498R | rg3aaΔ-133G | rg3aaΔ-133G/136K | rg3aaΔ-136K | rgIns1aa-164D | rgIns2aa | rgIns3aa | rgBRIS/08 wild-type | rgCOL/17 (2aaΔ) | rgWA/19 (3aaΔ) |
|---|---|---|---|---|---|---|---|---|---|---|---|---|---|---|---|---|---|---|---|
| rgIns2aa/129D/498R | | 0.150 | 0.048 | 0.997 | 0.195 | 0.112 | 0.960 | 0.818 | 0.306 | 0.872 | 0.584 | 0.303 | 0.274 | 0.133 | 0.937 | 0.282 | 1.000 | 0.232 | 0.378 |
| rgIns2aa/129D | 0.150 | | 1.000 | 0.940 | 1.000 | 1.000 | 0.994 | 1.000 | 1.000 | 0.999 | 1.000 | 1.000 | 1.000 | 1.000 | 0.000 | 1.000 | 0.137 | 1.000 | 1.000 |
| rgIns2aa/180I/498R | 0.048 | 1.000 | | 0.734 | 1.000 | 1.000 | 0.923 | 0.991 | 1.000 | 0.981 | 1.000 | 1.000 | 1.000 | 1.000 | 0.000 | 1.000 | 0.043 | 1.000 | 1.000 |
| rgIns2aa/180I | 0.997 | 0.940 | 0.734 | | 0.966 | 0.901 | 1.000 | 1.000 | 0.991 | 1.000 | 1.000 | 0.991 | 0.987 | 0.925 | 0.146 | 0.989 | 0.996 | 0.979 | 0.997 |
| rgIns2aa/498R | 0.195 | 1.000 | 1.000 | 0.966 | | 1.000 | 0.998 | 1.000 | 1.000 | 1.000 | 1.000 | 1.000 | 1.000 | 1.000 | 0.001 | 1.000 | 0.179 | 1.000 | 1.000 |
| rg2aaΔ-129D/180I | 0.112 | 1.000 | 1.000 | 0.901 | 1.000 | | 0.986 | 0.999 | 1.000 | 0.998 | 1.000 | 1.000 | 1.000 | 1.000 | 0.000 | 1.000 | 0.102 | 1.000 | 1.000 |
| rg2aaΔ-129D/180I/498R | 0.960 | 0.994 | 0.923 | 1.000 | 0.998 | 0.986 | | 1.000 | 1.000 | 1.000 | 1.000 | 1.000 | 1.000 | 0.991 | 0.054 | 1.000 | 0.952 | 0.999 | 1.000 |
| rg2aaΔ-180I | 0.818 | 1.000 | 0.991 | 1.000 | 1.000 | 0.999 | 1.000 | | 1.000 | 1.000 | 1.000 | 1.000 | 1.000 | 1.000 | 0.017 | 1.000 | 0.797 | 1.000 | 1.000 |
| rg2aaΔ-180I/498R | 0.306 | 1.000 | 1.000 | 0.991 | 1.000 | 1.000 | 1.000 | 1.000 | | 1.000 | 1.000 | 1.000 | 1.000 | 1.000 | 0.001 | 1.000 | 0.285 | 1.000 | 1.000 |
| rg2aaΔ-498R | 0.872 | 0.999 | 0.981 | 1.000 | 1.000 | 0.998 | 1.000 | 1.000 | 1.000 | | 1.000 | 1.000 | 1.000 | 0.999 | 0.024 | 1.000 | 0.854 | 1.000 | 1.000 |
| rg3aaΔ-133G | 0.584 | 1.000 | 1.000 | 1.000 | 1.000 | 1.000 | 1.000 | 1.000 | 1.000 | 1.000 | | 1.000 | 1.000 | 1.000 | 0.005 | 1.000 | 0.557 | 1.000 | 1.000 |
| rg3aaΔ-133G/136K | 0.303 | 1.000 | 1.000 | 0.991 | 1.000 | 1.000 | 1.000 | 1.000 | 1.000 | 1.000 | 1.000 | | 1.000 | 1.000 | 0.001 | 1.000 | 0.282 | 1.000 | 1.000 |
| rg3aaΔ-136K | 0.274 | 1.000 | 1.000 | 0.987 | 1.000 | 1.000 | 1.000 | 1.000 | 1.000 | 1.000 | 1.000 | 1.000 | | 1.000 | 0.001 | 1.000 | 0.254 | 1.000 | 1.000 |
| rgIns1aa-164D | 0.133 | 1.000 | 1.000 | 0.925 | 1.000 | 1.000 | 0.991 | 1.000 | 1.000 | 0.999 | 1.000 | 1.000 | 1.000 | | 0.000 | 1.000 | 0.121 | 1.000 | 1.000 |
| rgIns2aa | 0.937 | 0.000 | 0.000 | 0.146 | 0.001 | 0.000 | 0.054 | 0.017 | 0.001 | 0.024 | 0.005 | 0.001 | 0.001 | 0.000 | | 0.001 | 0.947 | 0.001 | 0.002 |
| rgIns3aa | 0.282 | 1.000 | 1.000 | 0.989 | 1.000 | 1.000 | 1.000 | 1.000 | 1.000 | 1.000 | 1.000 | 1.000 | 1.000 | 1.000 | 0.001 | | 0.262 | 1.000 | 1.000 |
| rgBRIS/08 wild-type | 1.000 | 0.137 | 0.043 | 0.996 | 0.179 | 0.102 | 0.952 | 0.797 | 0.285 | 0.854 | 0.557 | 0.282 | 0.254 | 0.121 | 0.947 | 0.262 | | 0.214 | 0.354 |
| rgCOL/17 (2aaΔ) | 0.232 | 1.000 | 1.000 | 0.979 | 1.000 | 1.000 | 0.999 | 1.000 | 1.000 | 1.000 | 1.000 | 1.000 | 1.000 | 1.000 | 0.001 | 1.000 | 0.214 | | 1.000 |
| rgWA/19 (3aaΔ) | 0.378 | 1.000 | 1.000 | 0.997 | 1.000 | 1.000 | 1.000 | 1.000 | 1.000 | 1.000 | 1.000 | 1.000 | 1.000 | 1.000 | 0.002 | 1.000 | 0.354 | 1.000 | |

**Table S12. Statistical signficance of pH-dependent differences within each virus.**

1.rgBRIS/08 wild-type

| pH level | pH5 | pH5.6 | pH6 | pH6.5 | pH7 |
|---|---|---|---|---|---|
| pH5 | | <0.0001 | <0.0001 | <0.0001 | <0.0001 |
| pH5.6 | <0.0001 | | 0.004 | 0.000 | <0.0001 |
| pH6 | <0.0001 | 0.004 | | 0.840 | 0.000 |
| pH6.5 | <0.0001 | 0.000 | 0.840 | | 0.000 |
| pH7 | <0.0001 | <0.0001 | 0.000 | 0.000 | |

2.rgCOL/17 (2aaΔ)

| pH level | pH5 | pH5.6 | pH6 | pH6.5 | pH7 |
|---|---|---|---|---|---|
| pH5 | | <0.0001 | <0.0001 | <0.0001 | <0.0001 |
| pH5.6 | <0.0001 | | 0.014 | 0.000 | 0.000 |
| pH6 | <0.0001 | 0.014 | | 0.642 | 0.029 |
| pH6.5 | <0.0001 | 0.000 | 0.642 | | 0.490 |
| pH7 | <0.0001 | 0.000 | 0.029 | 0.490 | |

3.rgIns2aa

| pH level | pH5 | pH5.6 | pH6 | pH6.5 | pH7 |
|---|---|---|---|---|---|
| pH5 | | 0.475 | 0.059 | 0.000 | <0.0001 |
| pH5.6 | 0.475 | | 0.815 | 0.012 | <0.0001 |
| pH6 | 0.059 | 0.815 | | 0.178 | <0.0001 |
| pH6.5 | 0.000 | 0.012 | 0.178 | | 0.000 |
| pH7 | <0.0001 | <0.0001 | <0.0001 | 0.000 | |

5.rg2aaΔ-180I

| pH level | pH5 | pH5.6 | pH6 | pH6.5 | pH7 |
|---|---|---|---|---|---|
| pH5 | | 0.228 | 0.002 | 0.000 | 0.000 |
| pH5.6 | 0.228 | | 0.396 | 0.012 | 0.000 |
| pH6 | 0.002 | 0.396 | | 0.545 | 0.001 |
| pH6.5 | 0.000 | 0.012 | 0.545 | | 0.098 |
| pH7 | 0.000 | 0.000 | 0.001 | 0.098 | |

6.rg2aaΔ-498R

| pH level | pH5 | pH5.6 | pH6 | pH6.5 | pH7 |
|---|---|---|---|---|---|
| pH5 | | <0.0001 | <0.0001 | <0.0001 | <0.0001 |
| pH5.6 | <0.0001 | | 0.386 | 0.000 | 0.000 |
| pH6 | <0.0001 | 0.386 | | 0.078 | 0.000 |
| pH6.5 | <0.0001 | 0.000 | 0.078 | | 0.077 |
| pH7 | <0.0001 | 0.000 | 0.000 | 0.077 | |

7.rg2aaΔ-129D/180I

| pH level | pH5 | pH5.6 | pH6 | pH6.5 | pH7 |
|---|---|---|---|---|---|
| pH5 | | 0.038 | 0.000 | 0.000 | <0.0001 |
| pH5.6 | 0.038 | | 0.020 | 0.000 | 0.000 |
| pH6 | 0.000 | 0.020 | | 0.376 | 0.021 |
| pH6.5 | 0.000 | 0.000 | 0.376 | | 0.688 |
| pH7 | <0.0001 | 0.000 | 0.021 | 0.688 | |

8.rg2aaΔ-180I/498R

| pH level | pH5 | pH5.6 | pH6 | pH6.5 | pH7 |
|---|---|---|---|---|---|
| pH5 | | 0.000 | <0.0001 | <0.0001 | <0.0001 |
| pH5.6 | 0.000 | | <0.0001 | <0.0001 | <0.0001 |
| pH6 | <0.0001 | <0.0001 | | 0.000 | 0.000 |
| pH6.5 | <0.0001 | <0.0001 | 0.000 | | 0.407 |
| pH7 | <0.0001 | <0.0001 | 0.000 | 0.407 | |

10.rgIns2aa/129D

| pH level | pH5 | pH5.6 | pH6 | pH6.5 | pH7 |
|---|---|---|---|---|---|
| pH5 | | <0.0001 | <0.0001 | <0.0001 | <0.0001 |
| pH5.6 | <0.0001 | | 0.241 | 0.000 | 0.000 |
| pH6 | <0.0001 | 0.241 | | 0.134 | 0.003 |
| pH6.5 | <0.0001 | 0.000 | 0.134 | | 0.614 |
| pH7 | <0.0001 | 0.000 | 0.003 | 0.614 | |

**Table S13. Statistical signficance of differences among viruses at 35°C.**

| Recombinant virus | rgIns2aa/129D /498R | rgIns2aa/129D | rgIns2aa/180I /498R | rgIns2aa/180I | rgIns2aa/498R | rg2aaΔ-129D/180I | rg2aaΔ-129D/180I/498R | rg2aaΔ-180I | rg2aaΔ-180I/498R | rg2aaΔ-498R | rg3aaΔ-133G | rg3aaΔ-133G/136K | rg3aaΔ-136K | rgIns1aa-164D | rgIns2aa | rgIns3aa | rgBRIS/08 wild-type | rgCOL/17 (2aaΔ) | rgWA/19 (3aaΔ) |
|---|---|---|---|---|---|---|---|---|---|---|---|---|---|---|---|---|---|---|---|
| rgIns2aa/129D/498R | | 1.000 | 1.000 | 0.077 | 1.000 | 1.000 | 0.000 | 0.077 | 1.000 | 0.077 | 1.000 | 0.970 | 1.000 | 1.000 | 1.000 | 0.000 | 1.000 | 1.000 | 1.000 |
| rgIns2aa/129D | 1.000 | | 1.000 | 0.077 | 1.000 | 1.000 | 0.000 | 0.077 | 1.000 | 0.077 | 1.000 | 0.970 | 1.000 | 1.000 | 1.000 | 0.000 | 1.000 | 1.000 | 1.000 |
| rgIns2aa/180I/498R | 1.000 | 1.000 | | 0.077 | 1.000 | 1.000 | 0.000 | 0.077 | 1.000 | 0.077 | 1.000 | 0.970 | 1.000 | 1.000 | 1.000 | 0.000 | 1.000 | 1.000 | 1.000 |
| rgIns2aa/180I | 0.077 | 0.077 | 0.077 | | 0.077 | 0.077 | 0.970 | 1.000 | 0.077 | 1.000 | 0.077 | 0.970 | 0.077 | 0.077 | 0.077 | 0.970 | 0.077 | 0.077 | 0.077 |
| rgIns2aa/498R | 1.000 | 1.000 | 1.000 | 0.077 | | 1.000 | 0.000 | 0.077 | 1.000 | 0.077 | 1.000 | 0.970 | 1.000 | 1.000 | 1.000 | 0.000 | 1.000 | 1.000 | 1.000 |
| rg2aaΔ-129D/180I | 1.000 | 1.000 | 1.000 | 0.077 | 1.000 | | 0.000 | 0.077 | 1.000 | 0.077 | 1.000 | 0.970 | 1.000 | 1.000 | 1.000 | 0.000 | 1.000 | 1.000 | 1.000 |
| rg2aaΔ-129D/180I/498R | 0.000 | 0.000 | 0.000 | 0.970 | 0.000 | 0.000 | | 0.970 | 0.000 | 0.970 | 0.000 | 0.077 | 0.000 | 0.000 | 0.000 | 1.000 | 0.000 | 0.000 | 0.000 |
| rg2aaΔ-180I | 0.077 | 0.077 | 0.077 | 1.000 | 0.077 | 0.077 | 0.970 | | 0.077 | 1.000 | 0.077 | 0.970 | 0.077 | 0.077 | 0.077 | 0.970 | 0.077 | 0.077 | 0.077 |
| rg2aaΔ-180I/498R | 1.000 | 1.000 | 1.000 | 0.077 | 1.000 | 1.000 | 0.000 | 0.077 | | 0.077 | 1.000 | 0.970 | 1.000 | 1.000 | 1.000 | 0.000 | 1.000 | 1.000 | 1.000 |
| rg2aaΔ-498R | 0.077 | 0.077 | 0.077 | 1.000 | 0.077 | 0.077 | 0.970 | 1.000 | 0.077 | | 0.077 | 0.970 | 0.077 | 0.077 | 0.077 | 0.970 | 0.077 | 0.077 | 0.077 |
| rg3aaΔ-133G | 1.000 | 1.000 | 1.000 | 0.077 | 1.000 | 1.000 | 0.000 | 0.077 | 1.000 | 0.077 | | 0.970 | 1.000 | 1.000 | 1.000 | 0.000 | 1.000 | 1.000 | 1.000 |
| rg3aaΔ-133G/136K | 0.970 | 0.970 | 0.970 | 0.970 | 0.970 | 0.970 | 0.077 | 0.970 | 0.970 | 0.970 | 0.970 | | 0.970 | 0.970 | 0.970 | 0.077 | 0.970 | 0.970 | 0.970 |
| rg3aaΔ-136K | 1.000 | 1.000 | 1.000 | 0.077 | 1.000 | 1.000 | 0.000 | 0.077 | 1.000 | 0.077 | 1.000 | 0.970 | | 1.000 | 1.000 | 0.000 | 1.000 | 1.000 | 1.000 |
| rgIns1aa-164D | 1.000 | 1.000 | 1.000 | 0.077 | 1.000 | 1.000 | 0.000 | 0.077 | 1.000 | 0.077 | 1.000 | 0.970 | 1.000 | | 1.000 | 0.000 | 1.000 | 1.000 | 1.000 |
| rgIns2aa | 1.000 | 1.000 | 1.000 | 0.077 | 1.000 | 1.000 | 0.000 | 0.077 | 1.000 | 0.077 | 1.000 | 0.970 | 1.000 | 1.000 | | 0.000 | 1.000 | 1.000 | 1.000 |
| rgIns3aa | 0.000 | 0.000 | 0.000 | 0.970 | 0.000 | 0.000 | 1.000 | 0.970 | 0.000 | 0.970 | 0.000 | 0.077 | 0.000 | 0.000 | 0.000 | | 0.000 | 0.000 | 0.000 |
| rgBRIS/08 wild-type | 1.000 | 1.000 | 1.000 | 0.077 | 1.000 | 1.000 | 0.000 | 0.077 | 1.000 | 0.077 | 1.000 | 0.970 | 1.000 | 1.000 | 1.000 | 0.000 | | 1.000 | 1.000 |
| rgCOL/17 (2aaΔ) | 1.000 | 1.000 | 1.000 | 0.077 | 1.000 | 1.000 | 0.000 | 0.077 | 1.000 | 0.077 | 1.000 | 0.970 | 1.000 | 1.000 | 1.000 | 0.000 | 1.000 | | 1.000 |
| rgWA/19 (3aaΔ) | 1.000 | 1.000 | 1.000 | 0.077 | 1.000 | 1.000 | 0.000 | 0.077 | 1.000 | 0.077 | 1.000 | 0.970 | 1.000 | 1.000 | 1.000 | 0.000 | 1.000 | 1.000 | |

**Table S14. Statistical signficance of differences among viruses at 40°C.**

| Recombinant virus | rgIns2aa/129D /498R | rgIns2aa/129D | rgIns2aa/180I /498R | rgIns2aa/180I | rgIns2aa/498R | rg2aaΔ-129D/180I | rg2aaΔ-129D/180I/498R | rg2aaΔ-180I | rg2aaΔ-180I/498R | rg2aaΔ-498R | rg3aaΔ-133G | rg3aaΔ-133G/136K | rg3aaΔ-136K | rgIns1aa-164D | rgIns2aa | rgIns3aa | rgBRIS/08 wild-type | rgCOL/17 (2aaΔ) | rgWA/19 (3aaΔ) |
|---|---|---|---|---|---|---|---|---|---|---|---|---|---|---|---|---|---|---|---|
| rgIns2aa/129D/498R | | 0.970 | 1.000 | 1.000 | 0.970 | 0.077 | 0.970 | 1.000 | 1.000 | 0.970 | 0.970 | 0.970 | 1.000 | 0.970 | 0.970 | 0.970 | 1.000 | 0.077 | 0.970 |
| rgIns2aa/129D | 0.970 | | 0.970 | 0.970 | 1.000 | 0.970 | 0.077 | 0.970 | 0.970 | 0.077 | 0.077 | 0.077 | 0.970 | 0.077 | 0.077 | 0.077 | 0.970 | 0.970 | 0.077 |
| rgIns2aa/180I/498R | 1.000 | 0.970 | | 1.000 | 0.970 | 0.077 | 0.970 | 1.000 | 1.000 | 0.970 | 0.970 | 0.970 | 1.000 | 0.970 | 0.970 | 0.970 | 1.000 | 0.077 | 0.970 |
| rgIns2aa/180I | 1.000 | 0.970 | 1.000 | | 0.970 | 0.077 | 0.970 | 1.000 | 1.000 | 0.970 | 0.970 | 0.970 | 1.000 | 0.970 | 0.970 | 0.970 | 1.000 | 0.077 | 0.970 |
| rgIns2aa/498R | 0.970 | 1.000 | 0.970 | 0.970 | | 0.970 | 0.077 | 0.970 | 0.970 | 0.077 | 0.077 | 0.077 | 0.970 | 0.077 | 0.077 | 0.077 | 0.970 | 0.970 | 0.077 |
| rg2aaΔ-129D/180I | 0.077 | 0.970 | 0.077 | 0.077 | 0.970 | | 0.000 | 0.077 | 0.077 | 0.000 | 0.000 | 0.000 | 0.077 | 0.000 | 0.000 | 0.000 | 0.077 | 1.000 | 0.000 |
| rg2aaΔ-129D/180I/498R | 0.970 | 0.077 | 0.970 | 0.970 | 0.077 | 0.000 | | 0.970 | 0.970 | 1.000 | 1.000 | 1.000 | 0.970 | 1.000 | 1.000 | 1.000 | 0.970 | 0.000 | 1.000 |
| rg2aaΔ-180I | 1.000 | 0.970 | 1.000 | 1.000 | 0.970 | 0.077 | 0.970 | | 1.000 | 0.970 | 0.970 | 0.970 | 1.000 | 0.970 | 0.970 | 0.970 | 1.000 | 0.077 | 0.970 |
| rg2aaΔ-180I/498R | 1.000 | 0.970 | 1.000 | 1.000 | 0.970 | 0.077 | 0.970 | 1.000 | | 0.970 | 0.970 | 0.970 | 1.000 | 0.970 | 0.970 | 0.970 | 1.000 | 0.077 | 0.970 |
| rg2aaΔ-498R | 0.970 | 0.077 | 0.970 | 0.970 | 0.077 | 0.000 | 1.000 | 0.970 | 0.970 | | 1.000 | 1.000 | 0.970 | 1.000 | 1.000 | 1.000 | 0.970 | 0.000 | 1.000 |
| rg3aaΔ-133G | 0.970 | 0.077 | 0.970 | 0.970 | 0.077 | 0.000 | 1.000 | 0.970 | 0.970 | 1.000 | | 1.000 | 0.970 | 1.000 | 1.000 | 1.000 | 0.970 | 0.000 | 1.000 |
| rg3aaΔ-133G/136K | 0.970 | 0.077 | 0.970 | 0.970 | 0.077 | 0.000 | 1.000 | 0.970 | 0.970 | 1.000 | 1.000 | | 0.970 | 1.000 | 1.000 | 1.000 | 0.970 | 0.000 | 1.000 |
| rg3aaΔ-136K | 1.000 | 0.970 | 1.000 | 1.000 | 0.970 | 0.077 | 0.970 | 1.000 | 1.000 | 0.970 | 0.970 | 0.970 | | 0.970 | 0.970 | 0.970 | 1.000 | 0.077 | 0.970 |
| rgIns1aa-164D | 0.970 | 0.077 | 0.970 | 0.970 | 0.077 | 0.000 | 1.000 | 0.970 | 0.970 | 1.000 | 1.000 | 1.000 | 0.970 | | 1.000 | 1.000 | 0.970 | 0.000 | 1.000 |
| rgIns2aa | 0.970 | 0.077 | 0.970 | 0.970 | 0.077 | 0.000 | 1.000 | 0.970 | 0.970 | 1.000 | 1.000 | 1.000 | 0.970 | 1.000 | | 1.000 | 0.970 | 0.000 | 1.000 |
| rgIns3aa | 0.970 | 0.077 | 0.970 | 0.970 | 0.077 | 0.000 | 1.000 | 0.970 | 0.970 | 1.000 | 1.000 | 1.000 | 0.970 | 1.000 | 1.000 | | 0.970 | 0.000 | 1.000 |
| rgBRIS/08 wild-type | 1.000 | 0.970 | 1.000 | 1.000 | 0.970 | 0.077 | 0.970 | 1.000 | 1.000 | 0.970 | 0.970 | 0.970 | 1.000 | 0.970 | 0.970 | 0.970 | | 0.077 | 0.970 |
| rgCOL/17 (2aaΔ) | 0.077 | 0.970 | 0.077 | 0.077 | 0.970 | 1.000 | 0.000 | 0.077 | 0.077 | 0.000 | 0.000 | 0.000 | 0.077 | 0.000 | 0.000 | 0.000 | 0.077 | | 0.000 |
| rgWA/19 (3aaΔ) | 0.970 | 0.077 | 0.970 | 0.970 | 0.077 | 0.000 | 1.000 | 0.970 | 0.970 | 1.000 | 1.000 | 1.000 | 0.970 | 1.000 | 1.000 | 1.000 | 0.970 | 0.000 | |

**Table S15. Statistical signficance of differences among viruses at 45°C.**

| Recombinant virus | rgIns2aa/129D /498R | rgIns2aa/129D | rgIns2aa/180I /498R | rgIns2aa/180I | rgIns2aa/498R | rg2aaΔ-129D/180I | rg2aaΔ-129D/180I/498R | rg2aaΔ-180I | rg2aaΔ-180I/498R | rg2aaΔ-498R | rg3aaΔ-133G | rg3aaΔ-133G/136K | rg3aaΔ-136K | rgIns1aa-164D | rgIns2aa | rgIns3aa | rgBRIS/08 wild-type | rgCOL/17 (2aaΔ) | rgWA/19 (3aaΔ) |
|---|---|---|---|---|---|---|---|---|---|---|---|---|---|---|---|---|---|---|---|
| rgIns2aa/129D/498R | | 1.000 | 0.970 | 1.000 | 1.000 | 0.000 | 1.000 | 1.000 | 1.000 | 1.000 | 1.000 | 0.970 | 1.000 | 1.000 | 1.000 | 0.000 | 1.000 | 0.077 | 1.000 |
| rgIns2aa/129D | 1.000 | | 0.970 | 1.000 | 1.000 | 0.000 | 1.000 | 1.000 | 1.000 | 1.000 | 1.000 | 0.970 | 1.000 | 1.000 | 1.000 | 0.000 | 1.000 | 0.077 | 1.000 |
| rgIns2aa/180I/498R | 0.970 | 0.970 | | 0.970 | 0.970 | 0.077 | 0.970 | 0.970 | 0.970 | 0.970 | 0.970 | 0.077 | 0.970 | 0.970 | 0.970 | 0.000 | 0.970 | 0.970 | 0.970 |
| rgIns2aa/180I | 1.000 | 1.000 | 0.970 | | 1.000 | 0.000 | 1.000 | 1.000 | 1.000 | 1.000 | 1.000 | 0.970 | 1.000 | 1.000 | 1.000 | 0.000 | 1.000 | 0.077 | 1.000 |
| rgIns2aa/498R | 1.000 | 1.000 | 0.970 | 1.000 | | 0.000 | 1.000 | 1.000 | 1.000 | 1.000 | 1.000 | 0.970 | 1.000 | 1.000 | 1.000 | 0.000 | 1.000 | 0.077 | 1.000 |
| rg2aaΔ-129D/180I | 0.000 | 0.000 | 0.077 | 0.000 | 0.000 | | 0.000 | 0.000 | 0.000 | 0.000 | 0.000 | 0.000 | 0.000 | 0.000 | 0.000 | 0.000 | 0.000 | 0.970 | 0.000 |
| rg2aaΔ-129D/180I/498R | 1.000 | 1.000 | 0.970 | 1.000 | 1.000 | 0.000 | | 1.000 | 1.000 | 1.000 | 1.000 | 0.970 | 1.000 | 1.000 | 1.000 | 0.000 | 1.000 | 0.077 | 1.000 |
| rg2aaΔ-180I | 1.000 | 1.000 | 0.970 | 1.000 | 1.000 | 0.000 | 1.000 | | 1.000 | 1.000 | 1.000 | 0.970 | 1.000 | 1.000 | 1.000 | 0.000 | 1.000 | 0.077 | 1.000 |
| rg2aaΔ-180I/498R | 1.000 | 1.000 | 0.970 | 1.000 | 1.000 | 0.000 | 1.000 | 1.000 | | 1.000 | 1.000 | 0.970 | 1.000 | 1.000 | 1.000 | 0.000 | 1.000 | 0.077 | 1.000 |
| rg2aaΔ-498R | 1.000 | 1.000 | 0.970 | 1.000 | 1.000 | 0.000 | 1.000 | 1.000 | 1.000 | | 1.000 | 0.970 | 1.000 | 1.000 | 1.000 | 0.000 | 1.000 | 0.077 | 1.000 |
| rg3aaΔ-133G | 1.000 | 1.000 | 0.970 | 1.000 | 1.000 | 0.000 | 1.000 | 1.000 | 1.000 | 1.000 | | 0.970 | 1.000 | 1.000 | 1.000 | 0.000 | 1.000 | 0.077 | 1.000 |
| rg3aaΔ-133G/136K | 0.970 | 0.970 | 0.077 | 0.970 | 0.970 | 0.000 | 0.970 | 0.970 | 0.970 | 0.970 | 0.970 | | 0.970 | 0.970 | 0.970 | 0.077 | 0.970 | 0.000 | 0.970 |
| rg3aaΔ-136K | 1.000 | 1.000 | 0.970 | 1.000 | 1.000 | 0.000 | 1.000 | 1.000 | 1.000 | 1.000 | 1.000 | 0.970 | | 1.000 | 1.000 | 0.000 | 1.000 | 0.077 | 1.000 |
| rgIns1aa-164D | 1.000 | 1.000 | 0.970 | 1.000 | 1.000 | 0.000 | 1.000 | 1.000 | 1.000 | 1.000 | 1.000 | 0.970 | 1.000 | | 1.000 | 0.000 | 1.000 | 0.077 | 1.000 |
| rgIns2aa | 1.000 | 1.000 | 0.970 | 1.000 | 1.000 | 0.000 | 1.000 | 1.000 | 1.000 | 1.000 | 1.000 | 0.970 | 1.000 | 1.000 | | 0.000 | 1.000 | 0.077 | 1.000 |
| rgIns3aa | 0.000 | 0.000 | 0.000 | 0.000 | 0.000 | 0.000 | 0.000 | 0.000 | 0.000 | 0.000 | 0.000 | 0.077 | 0.000 | 0.000 | 0.000 | | 0.000 | 0.000 | 0.000 |
| rgBRIS/08 wild-type | 1.000 | 1.000 | 0.970 | 1.000 | 1.000 | 0.000 | 1.000 | 1.000 | 1.000 | 1.000 | 1.000 | 0.970 | 1.000 | 1.000 | 1.000 | 0.000 | | 0.077 | 1.000 |
| rgCOL/17 (2aaΔ) | 0.077 | 0.077 | 0.970 | 0.077 | 0.077 | 0.970 | 0.077 | 0.077 | 0.077 | 0.077 | 0.077 | 0.000 | 0.077 | 0.077 | 0.077 | 0.000 | 0.077 | | 0.077 |
| rgWA/19 (3aaΔ) | 1.000 | 1.000 | 0.970 | 1.000 | 1.000 | 0.000 | 1.000 | 1.000 | 1.000 | 1.000 | 1.000 | 0.970 | 1.000 | 1.000 | 1.000 | 0.000 | 1.000 | 0.077 | |

**Table S16. Statistical signficance of differences among viruses at 50°C.**

| Recombinant virus | rgIns2aa/129D /498R | rgIns2aa/129D | rgIns2aa/180I /498R | rgIns2aa/180I | rgIns2aa/498R | rg2aaΔ-129D/180I | rg2aaΔ-129D/180I/498R | rg2aaΔ-180I | rg2aaΔ-180I/498R | rg2aaΔ-498R | rg3aaΔ-133G | rg3aaΔ-133G/136K | rg3aaΔ-136K | rgIns1aa-164D | rgIns2aa | rgIns3aa | rgBRIS/08 wild-type | rgCOL/17 (2aaΔ) | rgWA/19 (3aaΔ) |
|---|---|---|---|---|---|---|---|---|---|---|---|---|---|---|---|---|---|---|---|
| rgIns2aa/129D/498R | | 0.970 | 0.000 | 0.077 | 0.970 | 1.000 | 0.000 | 1.000 | 1.000 | 1.000 | 0.000 | 0.000 | 0.000 | 1.000 | 1.000 | 0.000 | 0.077 | 1.000 | 0.970 |
| rgIns2aa/129D | 0.970 | | 0.077 | 0.970 | 1.000 | 0.970 | 0.077 | 0.970 | 0.970 | 0.970 | 0.077 | 0.077 | 0.077 | 0.970 | 0.970 | 0.077 | 0.970 | 0.970 | 1.000 |
| rgIns2aa/180I/498R | 0.000 | 0.077 | | 0.970 | 0.077 | 0.000 | 1.000 | 0.000 | 0.000 | 0.000 | 1.000 | 1.000 | 1.000 | 0.000 | 0.000 | 1.000 | 0.970 | 0.000 | 0.077 |
| rgIns2aa/180I | 0.077 | 0.970 | 0.970 | | 0.970 | 0.077 | 0.970 | 0.077 | 0.077 | 0.077 | 0.970 | 0.970 | 0.970 | 0.077 | 0.077 | 0.970 | 1.000 | 0.077 | 0.970 |
| rgIns2aa/498R | 0.970 | 1.000 | 0.077 | 0.970 | | 0.970 | 0.077 | 0.970 | 0.970 | 0.970 | 0.077 | 0.077 | 0.077 | 0.970 | 0.970 | 0.077 | 0.970 | 0.970 | 1.000 |
| rg2aaΔ-129D/180I | 1.000 | 0.970 | 0.000 | 0.077 | 0.970 | | 0.000 | 1.000 | 1.000 | 1.000 | 0.000 | 0.000 | 0.000 | 1.000 | 1.000 | 0.000 | 0.077 | 1.000 | 0.970 |
| rg2aaΔ-129D/180I/498R | 0.000 | 0.077 | 1.000 | 0.970 | 0.077 | 0.000 | | 0.000 | 0.000 | 0.000 | 1.000 | 1.000 | 1.000 | 0.000 | 0.000 | 1.000 | 0.970 | 0.000 | 0.077 |
| rg2aaΔ-180I | 1.000 | 0.970 | 0.000 | 0.077 | 0.970 | 1.000 | 0.000 | | 1.000 | 1.000 | 0.000 | 0.000 | 0.000 | 1.000 | 1.000 | 0.000 | 0.077 | 1.000 | 0.970 |
| rg2aaΔ-180I/498R | 1.000 | 0.970 | 0.000 | 0.077 | 0.970 | 1.000 | 0.000 | 1.000 | | 1.000 | 0.000 | 0.000 | 0.000 | 1.000 | 1.000 | 0.000 | 0.077 | 1.000 | 0.970 |
| rg2aaΔ-498R | 1.000 | 0.970 | 0.000 | 0.077 | 0.970 | 1.000 | 0.000 | 1.000 | 1.000 | | 0.000 | 0.000 | 0.000 | 1.000 | 1.000 | 0.000 | 0.077 | 1.000 | 0.970 |
| rg3aaΔ-133G | 0.000 | 0.077 | 1.000 | 0.970 | 0.077 | 0.000 | 1.000 | 0.000 | 0.000 | 0.000 | | 1.000 | 1.000 | 0.000 | 0.000 | 1.000 | 0.970 | 0.000 | 0.077 |
| rg3aaΔ-133G/136K | 0.000 | 0.077 | 1.000 | 0.970 | 0.077 | 0.000 | 1.000 | 0.000 | 0.000 | 0.000 | 1.000 | | 1.000 | 0.000 | 0.000 | 1.000 | 0.970 | 0.000 | 0.077 |
| rg3aaΔ-136K | 0.000 | 0.077 | 1.000 | 0.970 | 0.077 | 0.000 | 1.000 | 0.000 | 0.000 | 0.000 | 1.000 | 1.000 | | 0.000 | 0.000 | 1.000 | 0.970 | 0.000 | 0.077 |
| rgIns1aa-164D | 1.000 | 0.970 | 0.000 | 0.077 | 0.970 | 1.000 | 0.000 | 1.000 | 1.000 | 1.000 | 0.000 | 0.000 | 0.000 | | 1.000 | 0.000 | 0.077 | 1.000 | 0.970 |
| rgIns2aa | 1.000 | 0.970 | 0.000 | 0.077 | 0.970 | 1.000 | 0.000 | 1.000 | 1.000 | 1.000 | 0.000 | 0.000 | 0.000 | 1.000 | | 0.000 | 0.077 | 1.000 | 0.970 |
| rgIns3aa | 0.000 | 0.077 | 1.000 | 0.970 | 0.077 | 0.000 | 1.000 | 0.000 | 0.000 | 0.000 | 1.000 | 1.000 | 1.000 | 0.000 | 0.000 | | 0.970 | 0.000 | 0.077 |
| rgBRIS/08 wild-type | 0.077 | 0.970 | 0.970 | 1.000 | 0.970 | 0.077 | 0.970 | 0.077 | 0.077 | 0.077 | 0.970 | 0.970 | 0.970 | 0.077 | 0.077 | 0.970 | | 0.077 | 0.970 |
| rgCOL/17 (2aaΔ) | 1.000 | 0.970 | 0.000 | 0.077 | 0.970 | 1.000 | 0.000 | 1.000 | 1.000 | 1.000 | 0.000 | 0.000 | 0.000 | 1.000 | 1.000 | 0.000 | 0.077 | | 0.970 |
| rgWA/19 (3aaΔ) | 0.970 | 1.000 | 0.077 | 0.970 | 1.000 | 0.970 | 0.077 | 0.970 | 0.970 | 0.970 | 0.077 | 0.077 | 0.077 | 0.970 | 0.970 | 0.077 | 0.970 | 0.970 | |

**Table S17. Statistical signficance of differences among viruses at 55°C.**

| Recombinant virus | rgIns2aa/129D/498R | rgIns2aa/129D | rgIns2aa/180I/498R | rgIns2aa/180I | rgIns2aa/498R | rg2aaΔ-129D/180I | rg2aaΔ-129D/180I/498R | rg2aaΔ-180I | rg2aaΔ-180I/498R | rg2aaΔ-498R | rg3aaΔ-133G | rg3aaΔ-133G/136K | rg3aaΔ-136K | rgIns1aa-164D | rgIns2aa | rgIns3aa | rgBRIS/08 wild-type | rgCOL/17 (2aaΔ) | rgWA/19 (3aaΔ) |
|---|---|---|---|---|---|---|---|---|---|---|---|---|---|---|---|---|---|---|---|
| rgIns2aa/129D/498R | | 0.970 | 1.000 | 0.077 | 1.000 | 1.000 | 0.000 | 1.000 | 1.000 | 1.000 | 0.000 | 0.000 | 0.000 | 0.000 | 1.000 | 0.000 | 0.077 | 1.000 | 0.000 |
| rgIns2aa/129D | 0.970 | | 0.970 | 0.970 | 0.970 | 0.970 | 0.077 | 0.970 | 0.970 | 0.970 | 0.077 | 0.077 | 0.077 | 0.077 | 0.970 | 0.077 | 0.970 | 0.970 | 0.077 |
| rgIns2aa/180I/498R | 1.000 | 0.970 | | 0.077 | 1.000 | 1.000 | 0.000 | 1.000 | 1.000 | 1.000 | 0.000 | 0.000 | 0.000 | 0.000 | 1.000 | 0.000 | 0.077 | 1.000 | 0.000 |
| rgIns2aa/180I | 0.077 | 0.970 | 0.077 | | 0.077 | 0.077 | 0.970 | 0.077 | 0.077 | 0.077 | 0.970 | 0.970 | 0.970 | 0.970 | 0.077 | 0.970 | 1.000 | 0.077 | 0.970 |
| rgIns2aa/498R | 1.000 | 0.970 | 1.000 | 0.077 | | 1.000 | 0.000 | 1.000 | 1.000 | 1.000 | 0.000 | 0.000 | 0.000 | 0.000 | 1.000 | 0.000 | 0.077 | 1.000 | 0.000 |
| rg2aaΔ-129D/180I | 1.000 | 0.970 | 1.000 | 0.077 | 1.000 | | 0.000 | 1.000 | 1.000 | 1.000 | 0.000 | 0.000 | 0.000 | 0.000 | 1.000 | 0.000 | 0.077 | 1.000 | 0.000 |
| rg2aaΔ-129D/180I/498R | 0.000 | 0.077 | 0.000 | 0.970 | 0.000 | 0.000 | | 0.000 | 0.000 | 0.000 | 1.000 | 1.000 | 1.000 | 1.000 | 0.000 | 1.000 | 0.970 | 0.000 | 1.000 |
| rg2aaΔ-180I | 1.000 | 0.970 | 1.000 | 0.077 | 1.000 | 1.000 | 0.000 | | 1.000 | 1.000 | 0.000 | 0.000 | 0.000 | 0.000 | 1.000 | 0.000 | 0.077 | 1.000 | 0.000 |
| rg2aaΔ-180I/498R | 1.000 | 0.970 | 1.000 | 0.077 | 1.000 | 1.000 | 0.000 | 1.000 | | 1.000 | 0.000 | 0.000 | 0.000 | 0.000 | 1.000 | 0.000 | 0.077 | 1.000 | 0.000 |
| rg2aaΔ-498R | 1.000 | 0.970 | 1.000 | 0.077 | 1.000 | 1.000 | 0.000 | 1.000 | 1.000 | | 0.000 | 0.000 | 0.000 | 0.000 | 1.000 | 0.000 | 0.077 | 1.000 | 0.000 |
| rg3aaΔ-133G | 0.000 | 0.077 | 0.000 | 0.970 | 0.000 | 0.000 | 1.000 | 0.000 | 0.000 | 0.000 | | 1.000 | 1.000 | 1.000 | 0.000 | 1.000 | 0.970 | 0.000 | 1.000 |
| rg3aaΔ-133G/136K | 0.000 | 0.077 | 0.000 | 0.970 | 0.000 | 0.000 | 1.000 | 0.000 | 0.000 | 0.000 | 1.000 | | 1.000 | 1.000 | 0.000 | 1.000 | 0.970 | 0.000 | 1.000 |
| rg3aaΔ-136K | 0.000 | 0.077 | 0.000 | 0.970 | 0.000 | 0.000 | 1.000 | 0.000 | 0.000 | 0.000 | 1.000 | 1.000 | | 1.000 | 0.000 | 1.000 | 0.970 | 0.000 | 1.000 |
| rgIns1aa-164D | 0.000 | 0.077 | 0.000 | 0.970 | 0.000 | 0.000 | 1.000 | 0.000 | 0.000 | 0.000 | 1.000 | 1.000 | 1.000 | | 0.000 | 1.000 | 0.970 | 0.000 | 1.000 |
| rgIns2aa | 1.000 | 0.970 | 1.000 | 0.077 | 1.000 | 1.000 | 0.000 | 1.000 | 1.000 | 1.000 | 0.000 | 0.000 | 0.000 | 0.000 | | 0.000 | 0.077 | 1.000 | 0.000 |
| rgIns3aa | 0.000 | 0.077 | 0.000 | 0.970 | 0.000 | 0.000 | 1.000 | 0.000 | 0.000 | 0.000 | 1.000 | 1.000 | 1.000 | 1.000 | 0.000 | | 0.970 | 0.000 | 1.000 |
| rgBRIS/08 wild-type | 0.077 | 0.970 | 0.077 | 1.000 | 0.077 | 0.077 | 0.970 | 0.077 | 0.077 | 0.077 | 0.970 | 0.970 | 0.970 | 0.970 | 0.077 | 0.970 | | 0.077 | 0.970 |
| rgCOL/17 (2aaΔ) | 1.000 | 0.970 | 1.000 | 0.077 | 1.000 | 1.000 | 0.000 | 1.000 | 1.000 | 1.000 | 0.000 | 0.000 | 0.000 | 0.000 | 1.000 | 0.000 | 0.077 | | 0.000 |
| rgWA/19 (3aaΔ) | 0.000 | 0.077 | 0.000 | 0.970 | 0.000 | 0.000 | 1.000 | 0.000 | 0.000 | 0.000 | 1.000 | 1.000 | 1.000 | 1.000 | 0.000 | 1.000 | 0.970 | 0.000 | |

**Table S18. Statistical signficance of differences among viruses at 60°C.**

| Recombinant virus | rgIns2aa/129D/498R | rgIns2aa/129D | rgIns2aa/180I/498R | rgIns2aa/180I | rgIns2aa/498R | rg2aaΔ-129D/180I | rg2aaΔ-129D/180I/498R | rg2aaΔ-180I | rg2aaΔ-180I/498R | rg2aaΔ-498R | rg3aaΔ-133G | rg3aaΔ-133G/136K | rg3aaΔ-136K | rgIns1aa-164D | rgIns2aa | rgIns3aa | rgBRIS/08 wild-type | rgCOL/17 (2aaΔ) | rgWA/19 (3aaΔ) |
|---|---|---|---|---|---|---|---|---|---|---|---|---|---|---|---|---|---|---|---|
| rgIns2aa/129D/498R | | 0.970 | 1.000 | 0.077 | 0.000 | 1.000 | 1.000 | 1.000 | 1.000 | 1.000 | 0.000 | 0.000 | 0.000 | 0.000 | 1.000 | 0.000 | 1.000 | 1.000 | 0.000 |
| rgIns2aa/129D | 0.970 | | 0.970 | 0.970 | 0.077 | 0.970 | 0.970 | 0.970 | 0.970 | 0.970 | 0.077 | 0.077 | 0.077 | 0.077 | 0.970 | 0.077 | 0.970 | 0.970 | 0.077 |
| rgIns2aa/180I/498R | 1.000 | 0.970 | | 0.077 | 0.000 | 1.000 | 1.000 | 1.000 | 1.000 | 1.000 | 0.000 | 0.000 | 0.000 | 0.000 | 1.000 | 0.000 | 1.000 | 1.000 | 0.000 |
| rgIns2aa/180I | 0.077 | 0.970 | 0.077 | | 0.970 | 0.077 | 0.077 | 0.077 | 0.077 | 0.077 | 0.970 | 0.970 | 0.970 | 0.970 | 0.077 | 0.970 | 0.077 | 0.077 | 0.970 |
| rgIns2aa/498R | 0.000 | 0.077 | 0.000 | 0.970 | | 0.000 | 0.000 | 0.000 | 0.000 | 0.000 | 1.000 | 1.000 | 1.000 | 1.000 | 0.000 | 1.000 | 0.000 | 0.000 | 1.000 |
| rg2aaΔ-129D/180I | 1.000 | 0.970 | 1.000 | 0.077 | 0.000 | | 1.000 | 1.000 | 1.000 | 1.000 | 0.000 | 0.000 | 0.000 | 0.000 | 1.000 | 0.000 | 1.000 | 1.000 | 0.000 |
| rg2aaΔ-129D/180I/498R | 1.000 | 0.970 | 1.000 | 0.077 | 0.000 | 1.000 | | 1.000 | 1.000 | 1.000 | 0.000 | 0.000 | 0.000 | 0.000 | 1.000 | 0.000 | 1.000 | 1.000 | 0.000 |
| rg2aaΔ-180I | 1.000 | 0.970 | 1.000 | 0.077 | 0.000 | 1.000 | 1.000 | | 1.000 | 1.000 | 0.000 | 0.000 | 0.000 | 0.000 | 1.000 | 0.000 | 1.000 | 1.000 | 0.000 |
| rg2aaΔ-180I/498R | 1.000 | 0.970 | 1.000 | 0.077 | 0.000 | 1.000 | 1.000 | 1.000 | | 1.000 | 0.000 | 0.000 | 0.000 | 0.000 | 1.000 | 0.000 | 1.000 | 1.000 | 0.000 |
| rg2aaΔ-498R | 1.000 | 0.970 | 1.000 | 0.077 | 0.000 | 1.000 | 1.000 | 1.000 | 1.000 | | 0.000 | 0.000 | 0.000 | 0.000 | 1.000 | 0.000 | 1.000 | 1.000 | 0.000 |
| rg3aaΔ-133G | 0.000 | 0.077 | 0.000 | 0.970 | 1.000 | 0.000 | 0.000 | 0.000 | 0.000 | 0.000 | | 1.000 | 1.000 | 1.000 | 0.000 | 1.000 | 0.000 | 0.000 | 1.000 |
| rg3aaΔ-133G/136K | 0.000 | 0.077 | 0.000 | 0.970 | 1.000 | 0.000 | 0.000 | 0.000 | 0.000 | 0.000 | 1.000 | | 1.000 | 1.000 | 0.000 | 1.000 | 0.000 | 0.000 | 1.000 |
| rg3aaΔ-136K | 0.000 | 0.077 | 0.000 | 0.970 | 1.000 | 0.000 | 0.000 | 0.000 | 0.000 | 0.000 | 1.000 | 1.000 | | 1.000 | 0.000 | 1.000 | 0.000 | 0.000 | 1.000 |
| rgIns1aa-164D | 0.000 | 0.077 | 0.000 | 0.970 | 1.000 | 0.000 | 0.000 | 0.000 | 0.000 | 0.000 | 1.000 | 1.000 | 1.000 | | 0.000 | 1.000 | 0.000 | 0.000 | 1.000 |
| rgIns2aa | 1.000 | 0.970 | 1.000 | 0.077 | 0.000 | 1.000 | 1.000 | 1.000 | 1.000 | 1.000 | 0.000 | 0.000 | 0.000 | 0.000 | | 0.000 | 1.000 | 1.000 | 0.000 |
| rgIns3aa | 0.000 | 0.077 | 0.000 | 0.970 | 1.000 | 0.000 | 0.000 | 0.000 | 0.000 | 0.000 | 1.000 | 1.000 | 1.000 | 1.000 | 0.000 | | 0.000 | 0.000 | 1.000 |
| rgBRIS/08 wild-type | 1.000 | 0.970 | 1.000 | 0.077 | 0.000 | 1.000 | 1.000 | 1.000 | 1.000 | 1.000 | 0.000 | 0.000 | 0.000 | 0.000 | 1.000 | 0.000 | | 1.000 | 0.000 |
| rgCOL/17 (2aaΔ) | 1.000 | 0.970 | 1.000 | 0.077 | 0.000 | 1.000 | 1.000 | 1.000 | 1.000 | 1.000 | 0.000 | 0.000 | 0.000 | 0.000 | 1.000 | 0.000 | 1.000 | | 0.000 |
| rgWA/19 (3aaΔ) | 0.000 | 0.077 | 0.000 | 0.970 | 1.000 | 0.000 | 0.000 | 0.000 | 0.000 | 0.000 | 1.000 | 1.000 | 1.000 | 1.000 | 0.000 | 1.000 | 0.000 | 0.000 | |

**Table S19. Statistical signficance of temperature-dependent differences within each virus.**

1.rgBRIS/08 wild-type

| Temperature | 33°C | 35°C | 40°C | 45°C | 50°C | 55°C | 60°C | 65°C |
|---|---|---|---|---|---|---|---|---|
| 33°C | | 1.0000 | 0.0025 | 0.0000 | 0.0000 | 0.0000 | 0.0000 | 0.0000 |
| 35°C | 1.0000 | | 0.0025 | 0.0000 | 0.0000 | 0.0000 | 0.0000 | 0.0000 |
| 40°C | 0.0025 | 0.0025 | | 0.4989 | 0.0000 | 0.0000 | 0.0000 | 0.0000 |
| 45°C | 0.0000 | 0.0000 | 0.4989 | | 0.0025 | 0.0000 | 0.0000 | 0.0000 |
| 50°C | 0.0000 | 0.0000 | 0.0000 | 0.0025 | | 0.0000 | 0.0000 | 0.0000 |
| 55°C | 0.0000 | 0.0000 | 0.0000 | 0.0000 | 0.0000 | | 0.4989 | 0.0000 |
| 60°C | 0.0000 | 0.0000 | 0.0000 | 0.0000 | 0.0000 | 0.4989 | | 0.0000 |
| 65°C | 0.0000 | 0.0000 | 0.0000 | 0.0000 | 0.0000 | 0.0000 | 0.0000 | |

2.rgCOL/17 (2aaΔ)

| Temperature | 33°C | 35°C | 40°C | 45°C | 50°C | 55°C | 60°C | 65°C |
|---|---|---|---|---|---|---|---|---|
| 33°C | | 1.0000 | 1.0000 | 0.4989 | 0.0000 | 0.0000 | 0.0000 | 0.0000 |
| 35°C | 1.0000 | | 1.0000 | 0.4989 | 0.0000 | 0.0000 | 0.0000 | 0.0000 |
| 40°C | 1.0000 | 1.0000 | | 0.4989 | 0.0000 | 0.0000 | 0.0000 | 0.0000 |
| 45°C | 0.4989 | 0.4989 | 0.4989 | | 0.0025 | 0.0000 | 0.0000 | 0.0000 |
| 50°C | 0.0000 | 0.0000 | 0.0000 | 0.0025 | | 0.0000 | 0.0000 | 0.0000 |
| 55°C | 0.0000 | 0.0000 | 0.0000 | 0.0000 | 0.0000 | | 0.0000 | 0.0000 |
| 60°C | 0.0000 | 0.0000 | 0.0000 | 0.0000 | 0.0000 | 0.0000 | | 0.0000 |
| 65°C | 0.0000 | 0.0000 | 0.0000 | 0.0000 | 0.0000 | 0.0000 | 0.0000 | |

3.rgIns2aa

| Temperature | 33°C | 35°C | 40°C | 45°C | 50°C | 55°C | 60°C | 65°C |
|---|---|---|---|---|---|---|---|---|
| 33°C | | 1.0000 | 0.0000 | 0.0000 | 0.0000 | 0.0000 | 0.0000 | 0.0000 |
| 35°C | 1.0000 | | 0.0000 | 0.0000 | 0.0000 | 0.0000 | 0.0000 | 0.0000 |
| 40°C | 0.0000 | 0.0000 | | 1.0000 | 1.0000 | 0.0000 | 0.0000 | 0.0000 |
| 45°C | 0.0000 | 0.0000 | 1.0000 | | 1.0000 | 0.0000 | 0.0000 | 0.0000 |
| 50°C | 0.0000 | 0.0000 | 1.0000 | 1.0000 | | 0.0000 | 0.0000 | 0.0000 |
| 55°C | 0.0000 | 0.0000 | 0.0000 | 0.0000 | 0.0000 | | 0.0000 | 0.0000 |
| 60°C | 0.0000 | 0.0000 | 0.0000 | 0.0000 | 0.0000 | 0.0000 | | 0.0000 |
| 65°C | 0.0000 | 0.0000 | 0.0000 | 0.0000 | 0.0000 | 0.0000 | 0.0000 | |

5.rg2aaΔ-180I

| Temperature | 33°C | 35°C | 40°C | 45°C | 50°C | 55°C | 60°C | 65°C |
|---|---|---|---|---|---|---|---|---|
| 33°C | | 0.0025 | 0.0025 | 0.0000 | 0.0000 | 0.0000 | 0.0000 | 0.0000 |
| 35°C | 0.0025 | | 1.0000 | 0.4989 | 0.4989 | 0.0000 | 0.0000 | 0.0000 |
| 40°C | 0.0025 | 1.0000 | | 0.4989 | 0.4989 | 0.0000 | 0.0000 | 0.0000 |
| 45°C | 0.0000 | 0.4989 | 0.4989 | | 1.0000 | 0.0000 | 0.0000 | 0.0000 |
| 50°C | 0.0000 | 0.4989 | 0.4989 | 1.0000 | | 0.0000 | 0.0000 | 0.0000 |
| 55°C | 0.0000 | 0.0000 | 0.0000 | 0.0000 | 0.0000 | | 0.0000 | 0.0000 |
| 60°C | 0.0000 | 0.0000 | 0.0000 | 0.0000 | 0.0000 | 0.0000 | | 0.0000 |
| 65°C | 0.0000 | 0.0000 | 0.0000 | 0.0000 | 0.0000 | 0.0000 | 0.0000 | |

6.rg2aaΔ-498R

| Temperature | 33°C | 35°C | 40°C | 45°C | 50°C | 55°C | 60°C | 65°C |
|---|---|---|---|---|---|---|---|---|
| 33°C | | 0.0025 | 0.0000 | 0.0000 | 0.0000 | 0.0000 | 0.0000 | 0.0000 |
| 35°C | 0.0025 | | 0.4989 | 0.4989 | 0.4989 | 0.0000 | 0.0000 | 0.0000 |
| 40°C | 0.0000 | 0.4989 | | 1.0000 | 1.0000 | 0.0000 | 0.0000 | 0.0000 |
| 45°C | 0.0000 | 0.4989 | 1.0000 | | 1.0000 | 0.0000 | 0.0000 | 0.0000 |
| 50°C | 0.0000 | 0.4989 | 1.0000 | 1.0000 | | 0.0000 | 0.0000 | 0.0000 |
| 55°C | 0.0000 | 0.0000 | 0.0000 | 0.0000 | 0.0000 | | 0.0000 | 0.0000 |
| 60°C | 0.0000 | 0.0000 | 0.0000 | 0.0000 | 0.0000 | 0.0000 | | 0.0000 |
| 65°C | 0.0000 | 0.0000 | 0.0000 | 0.0000 | 0.0000 | 0.0000 | 0.0000 | |

7.rgIns2aa/129D/498R

| Temperature | 33°C | 35°C | 40°C | 45°C | 50°C | 55°C | 60°C | 65°C |
|---|---|---|---|---|---|---|---|---|
| 33°C | | 1.0000 | 1.0000 | 1.0000 | 0.0000 | 0.0000 | 0.0000 | 0.0000 |
| 35°C | 1.0000 | | 1.0000 | 1.0000 | 0.0000 | 0.0000 | 0.0000 | 0.0000 |

| | | | | | | | | |
|---|---|---|---|---|---|---|---|---|
| 40°C | 1.0000 | 1.0000 | | 1.0000 | 0.0000 | 0.0000 | 0.0000 | 0.0000 |
| 45°C | 1.0000 | 1.0000 | 1.0000 | | 0.0000 | 0.0000 | 0.0000 | 0.0000 |
| 50°C | 0.0000 | 0.0000 | 0.0000 | 0.0000 | | 0.0000 | 0.0000 | 0.0000 |
| 55°C | 0.0000 | 0.0000 | 0.0000 | 0.0000 | 0.0000 | | 0.0000 | 0.0000 |
| 60°C | 0.0000 | 0.0000 | 0.0000 | 0.0000 | 0.0000 | 0.0000 | | 0.0000 |
| 65°C | 0.0000 | 0.0000 | 0.0000 | 0.0000 | 0.0000 | 0.0000 | 0.0000 | |

6.rg2aaΔ-498R

| Temperature | 33°C | 35°C | 40°C | 45°C | 50°C | 55°C | 60°C | 65°C |
|---|---|---|---|---|---|---|---|---|
| 33°C | | 0.0025 | 0.0000 | 0.0000 | 0.0000 | 0.0000 | 0.0000 | 0.0000 |
| 35°C | 0.0025 | | 0.4989 | 0.4989 | 0.4989 | 0.0000 | 0.0000 | 0.0000 |
| 40°C | 0.0000 | 0.4989 | | 1.0000 | 1.0000 | 0.0000 | 0.0000 | 0.0000 |
| 45°C | 0.0000 | 0.4989 | 1.0000 | | 1.0000 | 0.0000 | 0.0000 | 0.0000 |
| 50°C | 0.0000 | 0.4989 | 1.0000 | 1.0000 | | 0.0000 | 0.0000 | 0.0000 |
| 55°C | 0.0000 | 0.0000 | 0.0000 | 0.0000 | 0.0000 | | 0.0000 | 0.0000 |
| 60°C | 0.0000 | 0.0000 | 0.0000 | 0.0000 | 0.0000 | 0.0000 | | 0.0000 |
| 65°C | 0.0000 | 0.0000 | 0.0000 | 0.0000 | 0.0000 | 0.0000 | 0.0000 | |

7.rg2aaΔ-129D/180I

| Temperature | 33°C | 35°C | 40°C | 45°C | 50°C | 55°C | 60°C | 65°C |
|---|---|---|---|---|---|---|---|---|
| 33°C | | 1.0000 | 1.0000 | 1.0000 | 0.0000 | 0.0000 | 0.0000 | 0.0000 |
| 35°C | 1.0000 | | 1.0000 | 1.0000 | 0.0000 | 0.0000 | 0.0000 | 0.0000 |
| 40°C | 1.0000 | 1.0000 | | 1.0000 | 0.0000 | 0.0000 | 0.0000 | 0.0000 |
| 45°C | 1.0000 | 1.0000 | 1.0000 | | 0.0000 | 0.0000 | 0.0000 | 0.0000 |
| 50°C | 0.0000 | 0.0000 | 0.0000 | 0.0000 | | 0.0000 | 0.0000 | 0.0000 |
| 55°C | 0.0000 | 0.0000 | 0.0000 | 0.0000 | 0.0000 | | 0.0000 | 0.0000 |
| 60°C | 0.0000 | 0.0000 | 0.0000 | 0.0000 | 0.0000 | 0.0000 | | 0.0000 |
| 65°C | 0.0000 | 0.0000 | 0.0000 | 0.0000 | 0.0000 | 0.0000 | 0.0000 | |

8.rg2aaΔ-180I/498R

| Temperature | 33°C | 35°C | 40°C | 45°C | 50°C | 55°C | 60°C | 65°C |
|---|---|---|---|---|---|---|---|---|
| 33°C | | 1.0000 | 0.0025 | 0.0000 | 0.0000 | 0.0000 | 0.0000 | 0.0000 |
| 35°C | 1.0000 | | 0.0025 | 0.0000 | 0.0000 | 0.0000 | 0.0000 | 0.0000 |
| 40°C | 0.0025 | 0.0025 | | 0.4989 | 0.4989 | 0.0000 | 0.0000 | 0.0000 |
| 45°C | 0.0000 | 0.0000 | 0.4989 | | 1.0000 | 0.0000 | 0.0000 | 0.0000 |
| 50°C | 0.0000 | 0.0000 | 0.4989 | 1.0000 | | 0.0000 | 0.0000 | 0.0000 |
| 55°C | 0.0000 | 0.0000 | 0.0000 | 0.0000 | 0.0000 | | 0.0000 | 0.0000 |
| 60°C | 0.0000 | 0.0000 | 0.0000 | 0.0000 | 0.0000 | 0.0000 | | 0.0000 |
| 65°C | 0.0000 | 0.0000 | 0.0000 | 0.0000 | 0.0000 | 0.0000 | 0.0000 | |

10.rgIns2aa/129D

| Temperature | 33°C | 35°C | 40°C | 45°C | 50°C | 55°C | 60°C | 65°C |
|---|---|---|---|---|---|---|---|---|
| 33°C | | 1.0000 | 0.4989 | 0.0000 | 0.0000 | 0.0000 | 0.0000 | 0.0000 |
| 35°C | 1.0000 | | 0.4989 | 0.0000 | 0.0000 | 0.0000 | 0.0000 | 0.0000 |
| 40°C | 0.4989 | 0.4989 | | 0.0025 | 0.0000 | 0.0000 | 0.0000 | 0.0000 |
| 45°C | 0.0000 | 0.0000 | 0.0025 | | 0.4989 | 0.0000 | 0.0000 | 0.0000 |
| 50°C | 0.0000 | 0.0000 | 0.0000 | 0.4989 | | 0.0000 | 0.0000 | 0.0000 |
| 55°C | 0.0000 | 0.0000 | 0.0000 | 0.0000 | 0.0000 | | 0.0000 | 0.0000 |
| 60°C | 0.0000 | 0.0000 | 0.0000 | 0.0000 | 0.0000 | 0.0000 | | 0.0000 |
| 65°C | 0.0000 | 0.0000 | 0.0000 | 0.0000 | 0.0000 | 0.0000 | 0.0000 | |

11.rgIns2aa/180I

| Temperature | 33°C | 35°C | 40°C | 45°C | 50°C | 55°C | 60°C | 65°C |
|---|---|---|---|---|---|---|---|---|
| 33°C | | 0.0025 | 0.0025 | 0.0000 | 0.0000 | 0.0000 | 0.0000 | 0.0000 |
| 35°C | 0.0025 | | 1.0000 | 0.4989 | 0.0000 | 0.0000 | 0.0000 | 0.0000 |
| 40°C | 0.0025 | 1.0000 | | 0.4989 | 0.0000 | 0.0000 | 0.0000 | 0.0000 |
| 45°C | 0.0000 | 0.4989 | 0.4989 | | 0.0025 | 0.0000 | 0.0000 | 0.0000 |
| 50°C | 0.0000 | 0.0000 | 0.0000 | 0.0025 | | 0.0000 | 0.0000 | 0.0000 |
| 55°C | 0.0000 | 0.0000 | 0.0000 | 0.0000 | 0.0000 | | 0.0000 | 0.0000 |
| 60°C | 0.0000 | 0.0000 | 0.0000 | 0.0000 | 0.0000 | 0.0000 | | 0.0000 |
| 65°C | 0.0000 | 0.0000 | 0.0000 | 0.0000 | 0.0000 | 0.0000 | 0.0000 | |

12.rgIns2aa/498R

| Temperature | 33°C | 35°C | 40°C | 45°C | 50°C | 55°C | 60°C | 65°C |
|---|---|---|---|---|---|---|---|---|
| 33°C | | 1.0000 | 0.4989 | 0.0000 | 0.0000 | 0.0000 | 0.0000 | 0.0000 |
| 35°C | 1.0000 | | 0.4989 | 0.0000 | 0.0000 | 0.0000 | 0.0000 | 0.0000 |
| 40°C | 0.4989 | 0.4989 | | 0.0025 | 0.0000 | 0.0000 | 0.0000 | 0.0000 |
| 45°C | 0.0000 | 0.0000 | 0.0025 | | 0.4989 | 0.0000 | 0.0000 | 0.0000 |
| 50°C | 0.0000 | 0.0000 | 0.0000 | 0.4989 | | 0.0025 | 0.0000 | 0.0000 |
| 55°C | 0.0000 | 0.0000 | 0.0000 | 0.0000 | 0.0025 | | 0.0000 | 0.0000 |
| 60°C | 0.0000 | 0.0000 | 0.0000 | 0.0000 | 0.0000 | 0.0000 | | 0.0000 |
| 65°C | 0.0000 | 0.0000 | 0.0000 | 0.0000 | 0.0000 | 0.0000 | 0.0000 | |

15.rgIns2aa/129D/498R

| Temperature | 33°C | 35°C | 40°C | 45°C | 50°C | 55°C | 60°C | 65°C |
|---|---|---|---|---|---|---|---|---|
| 33°C | | 1.0000 | 0.4989 | 0.0000 | 0.0000 | 0.0000 | 0.0000 | 0.0000 |
| 35°C | 1.0000 | | 0.4989 | 0.0000 | 0.0000 | 0.0000 | 0.0000 | 0.0000 |
| 40°C | 0.4989 | 0.4989 | | 0.0025 | 0.0000 | 0.0000 | 0.0000 | 0.0000 |
| 45°C | 0.0000 | 0.0000 | 0.0025 | | 0.4989 | 0.0000 | 0.0000 | 0.0000 |
| 50°C | 0.0000 | 0.0000 | 0.0000 | 0.4989 | | 0.0000 | 0.0000 | 0.0000 |
| 55°C | 0.0000 | 0.0000 | 0.0000 | 0.0000 | 0.0000 | | 0.0000 | 0.0000 |
| 60°C | 0.0000 | 0.0000 | 0.0000 | 0.0000 | 0.0000 | 0.0000 | | 0.0000 |
| 65°C | 0.0000 | 0.0000 | 0.0000 | 0.0000 | 0.0000 | 0.0000 | 0.0000 | |

16.rgIns2aa/180I/498R

| Temperature | 33°C | 35°C | 40°C | 45°C | 50°C | 55°C | 60°C | 65°C |
|---|---|---|---|---|---|---|---|---|
| 33°C | | 1.0000 | 0.0025 | 0.0025 | 0.0000 | 0.0000 | 0.0000 | 0.0000 |
| 35°C | 1.0000 | | 0.0025 | 0.0025 | 0.0000 | 0.0000 | 0.0000 | 0.0000 |
| 40°C | 0.0025 | 0.0025 | | 1.0000 | 0.0000 | 0.0000 | 0.0000 | 0.0000 |
| 45°C | 0.0025 | 0.0025 | 1.0000 | | 0.0000 | 0.0000 | 0.0000 | 0.0000 |
| 50°C | 0.0000 | 0.0000 | 0.0000 | 0.0000 | | 1.0000 | 0.0000 | 0.0000 |
| 55°C | 0.0000 | 0.0000 | 0.0000 | 0.0000 | 1.0000 | | 0.0000 | 0.0000 |
| 60°C | 0.0000 | 0.0000 | 0.0000 | 0.0000 | 0.0000 | 0.0000 | | 0.0000 |
| 65°C | 0.0000 | 0.0000 | 0.0000 | 0.0000 | 0.0000 | 0.0000 | 0.0000 | |

18.rgWA/19 (3aaΔ)

| Temperature | 33°C | 35°C | 40°C | 45°C | 50°C | 55°C | 60°C | 65°C |
|---|---|---|---|---|---|---|---|---|
| 33°C | | 1.0000 | 0.0000 | 0.0000 | 0.0000 | 0.0000 | 0.0000 | 0.0000 |
| 35°C | 1.0000 | | 0.0000 | 0.0000 | 0.0000 | 0.0000 | 0.0000 | 0.0000 |
| 40°C | 0.0000 | 0.0000 | | 1.0000 | 0.4989 | 0.0000 | 0.0000 | 0.0000 |
| 45°C | 0.0000 | 0.0000 | 1.0000 | | 0.4989 | 0.0000 | 0.0000 | 0.0000 |
| 50°C | 0.0000 | 0.0000 | 0.4989 | 0.4989 | | 0.0000 | 0.0000 | 0.0000 |
| 55°C | 0.0000 | 0.0000 | 0.0000 | 0.0000 | 0.0000 | | 0.0000 | 0.0000 |
| 60°C | 0.0000 | 0.0000 | 0.0000 | 0.0000 | 0.0000 | 0.0000 | | 0.0000 |
| 65°C | 0.0000 | 0.0000 | 0.0000 | 0.0000 | 0.0000 | 0.0000 | 0.0000 | |

19.rgIns1aa-164D

| Temperature | 33°C | 35°C | 40°C | 45°C | 50°C | 55°C | 60°C | 65°C |
|---|---|---|---|---|---|---|---|---|
| 33°C | | 1.0000 | 0.0000 | 0.0000 | 0.0000 | 0.0000 | 0.0000 | 0.0000 |
| 35°C | 1.0000 | | 0.0000 | 0.0000 | 0.0000 | 0.0000 | 0.0000 | 0.0000 |
| 40°C | 0.0000 | 0.0000 | | 1.0000 | 1.0000 | 0.0000 | 0.0000 | 0.0000 |
| 45°C | 0.0000 | 0.0000 | 1.0000 | | 1.0000 | 0.0000 | 0.0000 | 0.0000 |
| 50°C | 0.0000 | 0.0000 | 1.0000 | 1.0000 | | 0.0000 | 0.0000 | 0.0000 |
| 55°C | 0.0000 | 0.0000 | 0.0000 | 0.0000 | 0.0000 | | 0.0000 | 0.0000 |
| 60°C | 0.0000 | 0.0000 | 0.0000 | 0.0000 | 0.0000 | 0.0000 | | 0.0000 |
| 65°C | 0.0000 | 0.0000 | 0.0000 | 0.0000 | 0.0000 | 0.0000 | 0.0000 | |

20.rgIns3aa

| Temperature | 33°C | 35°C | 40°C | 45°C | 50°C | 55°C | 60°C | 65°C |
|---|---|---|---|---|---|---|---|---|
| 33°C | | 0.0000 | 0.0000 | 0.0000 | 0.0000 | 0.0000 | 0.0000 | 0.0000 |
| 35°C | 0.0000 | | 1.0000 | 0.0000 | 0.0000 | 0.0000 | 0.0000 | 0.0000 |
| 40°C | 0.0000 | 1.0000 | | 0.0000 | 0.0000 | 0.0000 | 0.0000 | 0.0000 |

| | | | | | | | | |
|---|---|---|---|---|---|---|---|---|
| 45°C | 0.0000 | 0.0000 | 0.0000 | | 1.0000 | 0.0000 | 0.0000 | 0.0000 |
| 50°C | 0.0000 | 0.0000 | 0.0000 | 1.0000 | | 0.0000 | 0.0000 | 0.0000 |
| 55°C | 0.0000 | 0.0000 | 0.0000 | 0.0000 | 0.0000 | | 0.0000 | 0.0000 |
| 60°C | 0.0000 | 0.0000 | 0.0000 | 0.0000 | 0.0000 | 0.0000 | | 0.0000 |
| 65°C | 0.0000 | 0.0000 | 0.0000 | 0.0000 | 0.0000 | 0.0000 | 0.0000 | |

21.rg3aaΔ-133G

| Temperature | 33°C | 35°C | 40°C | 45°C | 50°C | 55°C | 60°C | 65°C |
|---|---|---|---|---|---|---|---|---|
| 33°C | | 1.0000 | 0.0000 | 0.0000 | 0.0000 | 0.0000 | 0.0000 | 0.0000 |
| 35°C | 1.0000 | | 0.0000 | 0.0000 | 0.0000 | 0.0000 | 0.0000 | 0.0000 |
| 40°C | 0.0000 | 0.0000 | | 1.0000 | 0.0000 | 0.0000 | 0.0000 | 0.0000 |
| 45°C | 0.0000 | 0.0000 | 1.0000 | | 0.0000 | 0.0000 | 0.0000 | 0.0000 |
| 50°C | 0.0000 | 0.0000 | 0.0000 | 0.0000 | | 0.0000 | 0.0000 | 0.0000 |
| 55°C | 0.0000 | 0.0000 | 0.0000 | 0.0000 | 0.0000 | | 0.0000 | 0.0000 |
| 60°C | 0.0000 | 0.0000 | 0.0000 | 0.0000 | 0.0000 | 0.0000 | | 0.0000 |
| 65°C | 0.0000 | 0.0000 | 0.0000 | 0.0000 | 0.0000 | 0.0000 | 0.0000 | |

22.rg3aaΔ-136K

| Temperature | 33°C | 35°C | 40°C | 45°C | 50°C | 55°C | 60°C | 65°C |
|---|---|---|---|---|---|---|---|---|
| 33°C | | 1.0000 | 0.0025 | 0.0000 | 0.0000 | 0.0000 | 0.0000 | 0.0000 |
| 35°C | 1.0000 | | 0.0025 | 0.0000 | 0.0000 | 0.0000 | 0.0000 | 0.0000 |
| 40°C | 0.0025 | 0.0025 | | 0.4989 | 0.0000 | 0.0000 | 0.0000 | 0.0000 |
| 45°C | 0.0000 | 0.0000 | 0.4989 | | 0.0000 | 0.0000 | 0.0000 | 0.0000 |
| 50°C | 0.0000 | 0.0000 | 0.0000 | 0.0000 | | 0.0000 | 0.0000 | 0.0000 |
| 55°C | 0.0000 | 0.0000 | 0.0000 | 0.0000 | 0.0000 | | 0.0000 | 0.0000 |
| 60°C | 0.0000 | 0.0000 | 0.0000 | 0.0000 | 0.0000 | 0.0000 | | 0.0000 |
| 65°C | 0.0000 | 0.0000 | 0.0000 | 0.0000 | 0.0000 | 0.0000 | 0.0000 | |

23.rg3aaΔ-133G/136K

| Temperature | 33°C | 35°C | 40°C | 45°C | 50°C | 55°C | 60°C | 65°C |
|---|---|---|---|---|---|---|---|---|
| 33°C | | 0.4989 | 0.0000 | 0.0000 | 0.0000 | 0.0000 | 0.0000 | 0.0000 |
| 35°C | 0.4989 | | 0.0025 | 0.0000 | 0.0000 | 0.0000 | 0.0000 | 0.0000 |
| 40°C | 0.0000 | 0.0025 | | 0.4989 | 0.0000 | 0.0000 | 0.0000 | 0.0000 |
| 45°C | 0.0000 | 0.0000 | 0.4989 | | 0.0025 | 0.0000 | 0.0000 | 0.0000 |
| 50°C | 0.0000 | 0.0000 | 0.0000 | 0.0025 | | 0.0000 | 0.0000 | 0.0000 |
| 55°C | 0.0000 | 0.0000 | 0.0000 | 0.0000 | 0.0000 | | 0.0000 | 0.0000 |
| 60°C | 0.0000 | 0.0000 | 0.0000 | 0.0000 | 0.0000 | 0.0000 | | 0.0000 |
| 65°C | 0.0000 | 0.0000 | 0.0000 | 0.0000 | 0.0000 | 0.0000 | 0.0000 | |
